# SCAN-ACT: Adoptive T Cell Therapy Target Discovery Through Single-Cell Transcriptomics

**DOI:** 10.1101/2025.01.18.633736

**Authors:** Stefano Testa, Aastha Pal, Ajay Subramanian, Sushama Varma, Jack Pengfei Tang, Danielle Graham, Sara Arfan, Minggui Pan, Nam Q. Bui, Kristen N. Ganjoo, Sarah Dry, Paul Huang, Matt van de Rijn, Wei Jiang, Anusha Kalbasi, Everett J. Moding

## Abstract

The FDA approval of T cell receptor-engineered T cells (TCR-T) for synovial sarcoma demonstrates the potential for adoptive T cell therapies (ACTs) in solid tumors. However, the paucity of tumor-specific targets without expression in normal tissues remains a major bottleneck, especially in rare cancer subtypes. Here, we present a comprehensive computational pipeline called SCAN-ACT that leverages single cell RNA sequencing and multi-omics data from tumor and normal tissues to nominate and prioritize targets for both chimeric antigen receptor (CAR)- and TCR-T cells. For surface membrane targets, SCAN-ACT proposes target pairs for bispecific Boolean logic-gated CAR T cells. For peptide-MHC targets, SCAN-ACT proposes intracellular peptides bound to a diverse set of human leukocyte antigens. We applied the SCAN-ACT pipeline to soft tissue sarcoma (STS), analyzing 986,749 single cells to identify and prioritize 395 monospecific CAR-T targets, 14,192 bispecific CAR-T targets, and 5,020 peptide-MHC targets for TCR-T cells. Selected targets were validated experimentally by protein expression and for peptide-MHC binding. Proposed targets and target pairs reflected the mesenchymal, neuronal, and hematopoietic ontogeny of STS. This work provides a robust data repository along with a web-based and user-friendly set of analysis tools to accelerate ACT development for solid tumors.

## Introduction

Despite profound success in the treatment of hematologic malignancies, engineered adoptive T cell therapies (ACT) have had limited success in solid tumors^1–3^. While novel engineering strategies can address the major hurdles related to T cell function^4^, including poor tumor infiltration^5^, terminal differentiation, effector dysfunction, and persistence or memory formation^6–9^, a lack of tumor-specific targets with uniformly high expression and low risk of on-target off-tumor toxicity remains a challenge^10,11^. Advances in bispecific antigen targeting, synthetic gene circuits, and Boolean logic-gating in engineered T cells have opened new opportunities to improve antigen-discrimination between normal and tumor cells and increase the targetable antigen pool^4,12,13^. However, there are no repositories of suitable target pairs for Boolean logic CAR-T cells in solid tumors.

Prior ACT target discovery has focused on bulk RNA-sequencing and protein expression from tumor and normal samples^3,14,15^, which is confounded by non-malignant cells in the tumor microenvironment. Similarly, analysis of cancer cell lines^16^ may not reflect gene expression in vivo or the heterogeneity of expression between individual cancer cells and across patients. Furthermore, robust bulk transcriptomic and proteomic datasets are not available for many less common human cancers. Compared to bulk RNA-seq and proteomics, scRNA-seq more accurately represents the transcriptomic profile and heterogeneity of individual tumor cells enabling the precise identification of tumor-specific genes^17,18^.

Soft tissue sarcomas (STS) are rare and heterogeneous mesenchymal cancers with high rates of metastatic disease and limited therapeutic options^19^. Although chemotherapy remains the primary treatment for metastatic STS, most patients do not respond and the median overall survival is less than two years^20,21^. Immune checkpoint blockers (ICBs) targeting programmed cell death protein 1 (PD-1) and cytotoxic T-lymphocyte antigen-4 (CTLA-4) have shown promise in specific STS histotypes^22–24^, but are ineffective for most patients^25,26^. Recent studies have shown promising efficacy for TCR-T cells in STS. Afamitresgene autoleucel (afami-cel) targeting the cancer-testis antigen MAGE-A4 in patients with an HLA*02:01 allele achieved a 37% overall response rate in heavily treated patients with synovial sarcoma (SS) and myxoid round cell liposarcoma (MRCLS)^27,28^, leading to the first FDA approval of a genetically engineered ACT in solid tumors. Likewise, the most promising results for TCR-T cell products targeting NY-ESO-1^29,30^ have been reported in patients with STS^31^. However, additional targets will be critical to expand the availability of ACTs to patients with STS that do not express MAGE-A4 and NY-ESO-1. Furthermore, most patients lack the HLA-A*02:01 allele, which is most frequently expressed in Europeans, raising concerns about the equity of current ACTs^32^. Taken together, STS represent a challenging disease model with known effective ACT targets that can be used to benchmark target identification pipelines.

Here, we leveraged single cell RNA sequencing (scRNA-seq) from patient tumors and collated normal tissue and tumor genetic, transcriptomic, and proteomic data across public databases to develop a comprehensive computational pipeline for discovery of ACT targets in solid tumors, called SCAN-ACT (<u>S</u>ingle <u>C</u>ell <u>A</u>nalysis <u>N</u>etwork for <u>A</u>doptive <u>C</u>ellular therapy <u>T</u>argets). We applied SCAN-ACT in STS to build CellSARCTx (<u>Cell</u>ular <u>SARC</u>oma <u>Therapy</u>), a database describing and prioritizing over 20,000 TCR-T and CAR-T cell targets, including those for Boolean logic-gated and peptide-centric CAR-T cells (PC-CAR-T). Proposed targets captured the shared mesenchymal, neuronal and hematopoietic gene expression signatures of STS, reflecting the intertwined developmental ontogeny of these three lineages from the mesoderm and neural crest. We validated protein expression of select targets by IHC and binding of peptides derived from intracellular proteins to HLA molecules with a novel yeast display technology. We present two interactive web interfaces to explore and evaluate the identified STS ACT targets (CellSARCTx) and to apply SCAN-ACT to new scRNA-Seq samples (SCAN-ACT Custom). SCAN-ACT can be applied to individual patient samples to aid in therapy selection or broader panels of patients to identify conserved targets and accelerate ACT development. SCAN-ACT is freely accessible for academic use at scanact.stanford.edu.

## Results

### Overview of the SCAN-ACT pipeline

SCAN-ACT identifies tumor up-regulated and down-regulated genes through differential gene expression (DGE) analysis using scRNA-seq (**Figure 1A**). Tumor up-regulated genes are then compared against clinically tested CAR-T targets to identify novel targets with predicted low risk of on-target off-tumor toxicity. SCAN-ACT then defines Boolean logic-gated CAR-T target pairs by measuring tumor and normal cell/tissue co-expression patterns. Tumor-specific MHC-restricted peptides (*pMHCs*) targetable by TCR-T or PC-CAR-T cells are identified by integrating peptide immunogenicity and HLA presentation predictions with *pMHC* cross-reactivity assessment. Lastly, SCAN-ACT prioritizes targets by integrating target homogeneity with normal tissue/cell expression, and functional, genomic, and proteomics tumor data.

**Figure 1.**
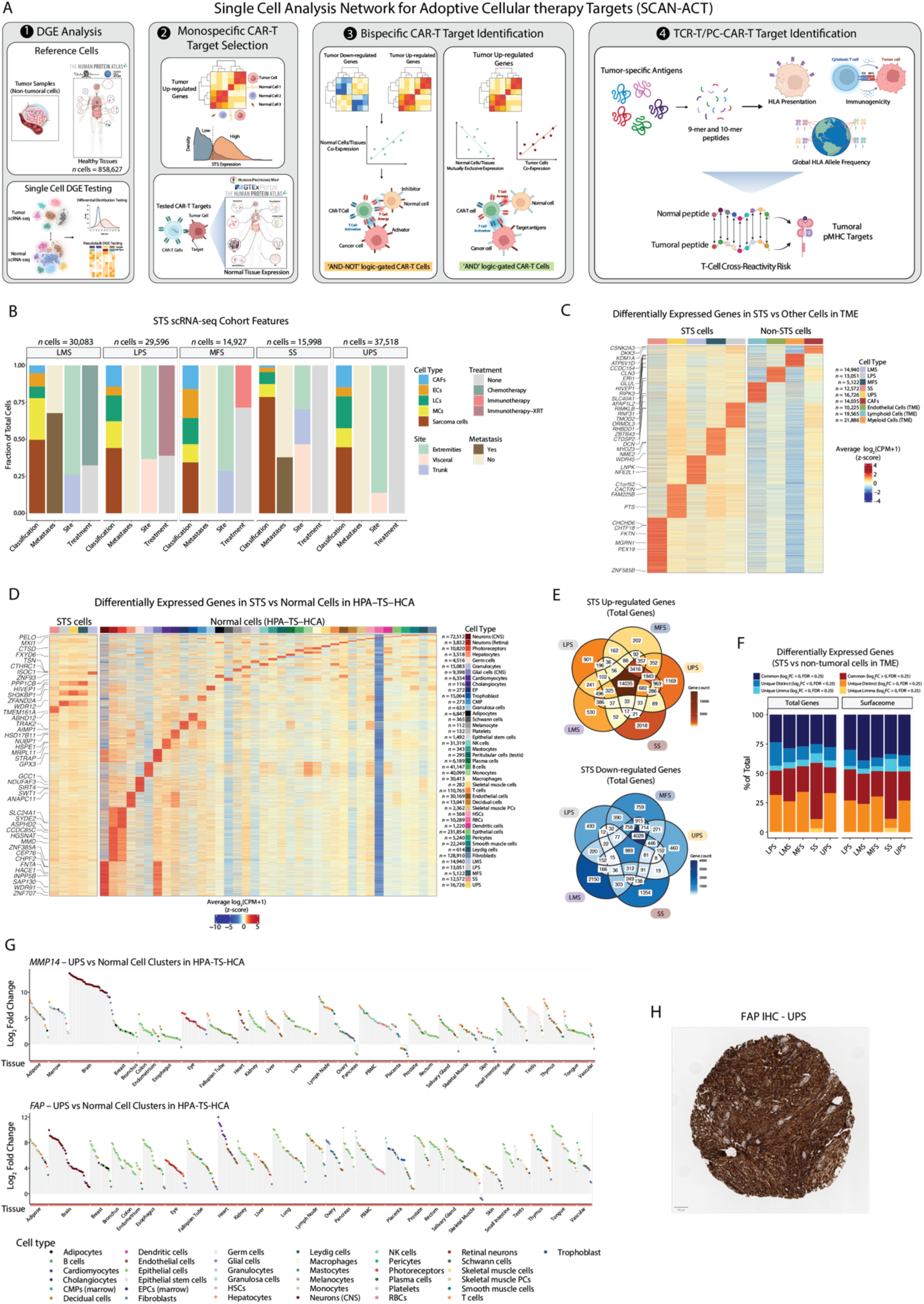
scRNA-seq for cellular therapy target identification. (A). Flow diagram showing the full SCAN-ACT pipeline. Stacked bar charts showing the fraction of cells identified as tumoral, endothelial, myeloid, lymphoid, or CAFs, sample site of origin, prior treatment, and metastatic or primary tumors. (C, D) Heatmaps showing average RNA expression in STS versus non-tumoral cells in the TME (C) and normal cells in the HPA, TS, and HCA (D). RNA expression (log2(CPM+1)) was averaged and median-centered across cell types, and z-scored across genes. The number of cells analyzed is shown in the legend. Only genes differentially expressed in each STS and measured in all samples are shown. (E) Venn diagrams showing the number of STS up-regulated and down-regulated genes shared between histotypes. (F) Stacked bar chart showing the fraction of DEGs in STS versus non-tumoral cells in the TME, identified with Limma-voom, distinct, or both methods, considering surfaceome and total genes separately. (G) Lollipop plots showing the log2FC in expression for MMP14 and FAP between UPS cells and the normal cell clusters in the HPA, TS, and HCA using Limma-voom. Only statistically significant comparisons are shown (FDR < 0.25). (H) FAP IHC expression on tissue microarrays of UPS samples. CAFs: cancer associated fibroblasts; ECs: endothelial cells; LCs: lymphoid cells; MCs: myeloid cells; Mφ: Macrophages.

### scRNA-seq for cellular therapy target identification in STS

To identify ACT targets across STS histologies, we applied SCAN-ACT to newly sequenced and publicly available scRNA-seq data^17,33^ from both translocation-driven and non-translocation-driven STS (**Table S1**). Our scRNA-seq dataset included 32 samples from 25 patients, totaling 128,122 cells after quality control (**Figure 1B, Figure S1A**, **S1B**, **S1C**, **S1E, S1G, S1I**). We included visceral (31%) and extremity/trunk (69%) tumor samples from male (72%) and female (28%) patients aged 22 to 77 years old. Samples included both primary (75%) and metastatic tumors (25%) collected before (84%) or after treatment (16%). Sarcoma cells were identified among the CD45^−^ cells (PTPRC count = 0), using inferred copy number abnormalities^34,35^ and transcriptional similarity to bulk samples of the same histology from The Cancer Genome Atlas (TCGA)^36^ (Similarity Score)^33^ (**Figure S1D, S1F**, **S1H**, **S1H**, **S1J**). Within the tumor microenvironment (TME), endothelial cells (ECs), myeloid cells (MCs), lymphoid cells (LCs), and cancer-associated fibroblasts (CAFs) were defined using canonical markers from the literature (**Table S2**).

### Identification of genes differentially expressed by malignant cells within the STS TME

To identify tumor-specific genes, SCAN-ACT performs DGE analysis between sarcoma cells and non-tumoral cells from the TME using Limma-voom^37^, a pseudo-bulk method, and distinct^38^, a differential distribution method (**Figure 1C**). We also compared sarcoma cells with normal cell clusters from 30 scRNA-seq datasets from the Human Protein Atlas (HPA)^39,40^, Tabula Sapiens (TS)^41^, and the Human Cell Atlas (HCA)^42^ covering the major normal human tissues (858,627 cells and 164 samples, **Figure 1D**). Unsupervised hierarchical clustering demonstrated that STS were most similar to CAFs and ECs within the TME (**Figure S2A**, **S2C**) and mesenchymal cells in normal tissues (**Figure S2B**, **S2D**), consistent with their putative mesenchymal origin. In total, we identified 29,322 up-regulated genes and 15,812 down-regulated genes in STS cells (**Table S3, S4**). SS and LMS had the most unique up-regulated and down-regulated genes, respectively, while UPS, LPS, and MFS shared the most up-regulated and down-regulated genes consistent with their biological similarity^36^ (**Figure 1E**, **Figure S2E**).

There was concordance between Limma-voom and distinct for DGE analysis in the TME (**Figure 1F**), with distinct identifying more unique differentially expressed genes. By measuring differences in gene expression distribution between cell types, distinct identifies subtle differences in expression^38^, which can be advantageous when comparing transcriptionally similar cells like STS and other mesenchymal cells in the TME. This approach identified a pool of sarcoma-specific genes, including many like *FAP* or *MMP14* in UPS which are up-regulated in STS compared to almost all normal cell types (**Figure 1G**). Notably, *FAP* has been targeted in solid tumors due to its TME expression^43^, though we found that *FAP* has intrinsic expression in STS cells, particularly LPS, MFS, and UPS, consistent with IHC data from UPS patient samples (**Figure 1H**), where 31/57 (54%) patients had *FAP* expression in ≥ 75% of cells (**Figure S2F, S2G, S2H**). Other targets classically associated with the stroma that were upregulated in malignant cells within the STS TME include *MRC2*, *CD248*, and *PDGFRB*.

### Monospecific CAR-T cell targets in STS

The ideal monospecific CAR-T cell target would have homogeneously high expression on cancer cells with low to no expression on normal cells to avoid on-target off-tumor toxicity^44^. However, development of CAR-T cells for solid tumors has been hampered by the scarcity of antigens that meet these conditions with few targets tested in humans^10^. To address this, we filtered the STS up-regulated genes to eliminate genes frequently expressed more highly in normal cells than STS cells. We then selected genes coding for surface proteins with high expression in STS by cross-referencing our scRNA-seq data with the TCGA STS bulk RNA-seq data (TCGA SARC) and the HPA transcriptomics data in STS cell lines. Finally, we removed targets with high RNA and protein expression in normal cells and tissues using CAR-T targets tested in humans as references for maximum tolerable on-target off-tumor toxicity risk (**Figure 2A**, **Table S4**, **Table S5**, **Table S6**, **Table S7**)^45^.

**Figure 2.**
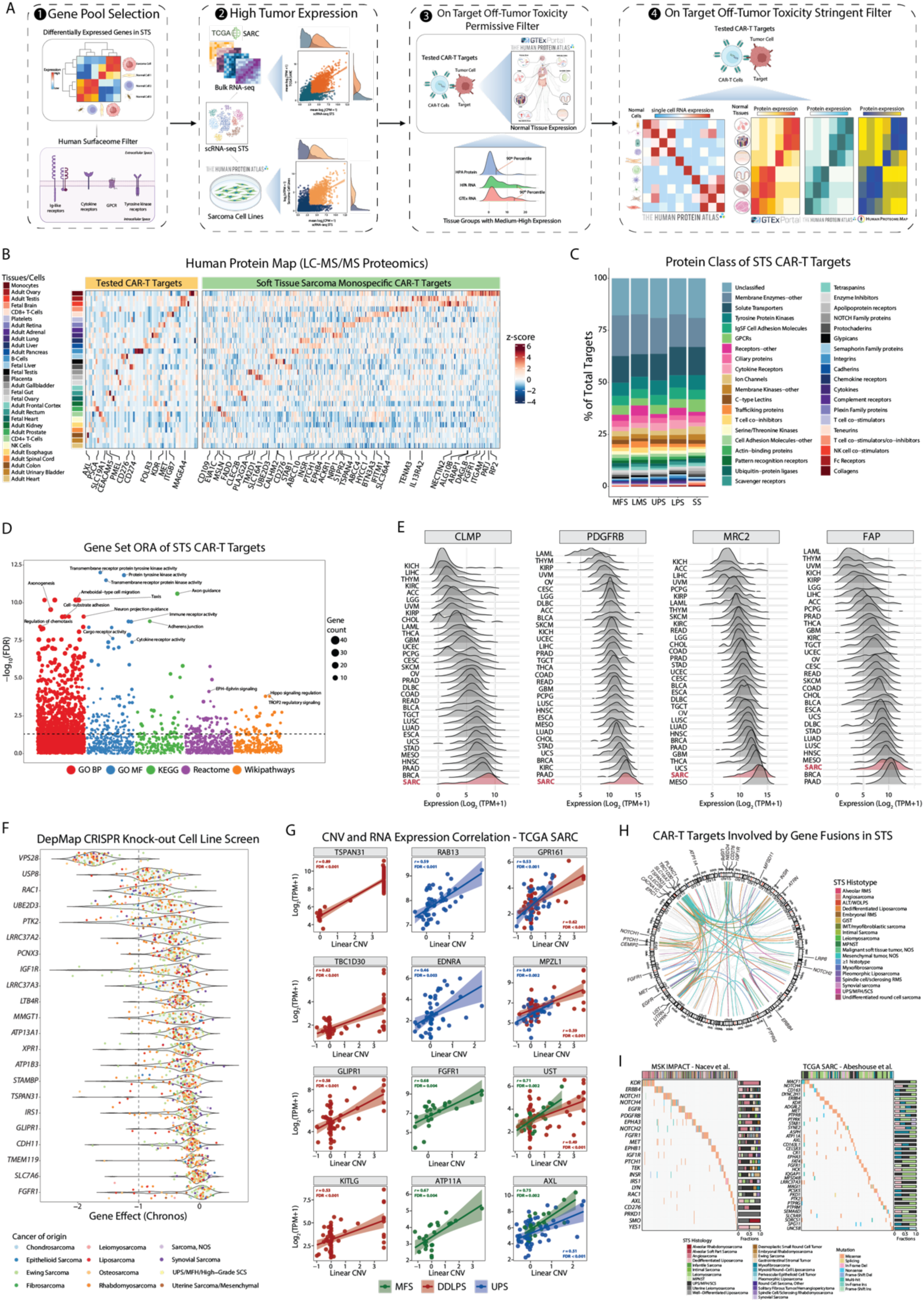
Monospecific CAR-T cell targets in STS. (A) Diagram showing discovery pipeline for monospecific STS CAR-T targets. (B) Heatmap showing the protein expression in normal tissues and cells or tissue groups for CAR-T targets tested in humans (left panel) and for the STS targets (right panel) in the HPM dataset. (C) Bar chart showing the proportion of STS CAR-T targets for each protein class/family. (D) Scatter plot showing gene set ORA on the 449 CAR-T targets, using the gene ontology (GO) molecular function (MF) and biological process (BP), as well as the KEGG, Reactome, and WikiPathways terms. The x-axis shows the –log10(FDR) while the dots size indicates the number of targets per gene set. The horizontal line intersects at the significance threshold (FDR = 0.05). (E) Ridge plots showing RNA expression (log2(TPM+1)) in the Pan-Cancer TCGA Atlas for STS CAR-T targets. STS samples (SARC) are highlighted in bold red. (F) Violin plots showing the effect of CAR-T targets KO on sarcoma cell line growth from DepMap, quantified using the Chronos algorithm. Negative values indicate genes essential for cell growth. The vertical line indicates the median Chronos value for known essential genes (–1). Dots are colored based on the cell line cancer of origin. (G) Pearson correlation between linear CNV level and RNA expression in TCGA SARC for monospecific CAR-T targets. P-values were adjusted using the BH method setting significance to FDR < 0.01. (H) Circos plot showing gene fusions involving a CAR-T target in STS from the Mitelman database, MSK-IMPACT, and TCGA SARC. Colored links show fusions for the top 30 most recurrently fused genes, with link colors indicating the STS histotype. (I) Oncoplots showing recurring SNVs/Indels for monospecific CAR-T targets in MSK-IMPACT (left) and TCGA SARC (right). ALT: Atypical Lipomatous Tumor; GIST: Gastrointestinal Stromal Tumor; MPNST: Malignant Peripheral Nerve Sheath Tumor; IMT: Inflammatory Myofibroblastic Tumor; MFH: Malignant Fibrous Histiocytoma; SCS: High-grade Spindle Cell Sarcoma.

We identified targets with RNA and protein expression profiles in normal tissues that mimic the normal tissue profiles of clinically tested CAR-T targets (**Figure 2B, Figure S3A**). Our approach reassuringly identified several targets currently in clinical trials for sarcomas, including *IGF1R*^46^*, CD163*, *EPHB4*, *FGFR1, KDR*, *NCAM1*, *SMO*, *PSEN1* and *PSEN2*^47^, and *NT5E*/CD73^48^. Overall, we found 395 unique targets, with 87 common to all sarcomas and 41 common to LPS, MFS, and UPS (**Figure S3B**, **Table S8, Figure S3C**, **S3D**). LPS and SS were the histotypes with the highest number of unique targets with 30 and 21, respectively.

In each sarcoma, transmembrane enzymes were the most common class (**Figure 2C**, **Table S9**), including metalloproteases of the ADAM family^49^ (*ADAM10*, *ADAM33*), metallopeptidases (*MME*), serine proteases (*PRSS21*, *FAP*), and protein tyrosine phosphatases (*PTPRJ*, *PTPRB*). Other common target classes were glypicans^50^ (*GPC3*, *GPC6*), immune checkpoint molecules^51^ (*CD276*, *NECTIN2*, *BTN2A2*), protein tyrosine kinases^52^ (*FGFR1*, *ERBB4*, *EFNA4*), NOTCH family receptors^53^ (*NOTCH2*, *NOTCH4*), G-protein coupled receptors – GPCRs^54^ (*ADGRL2*, *GPR107*), cell adhesion molecules of the Immunoglobulin superfamily – Ig-like CAMs (*CLMP*, *NEO1*), semaphorins^55^ (*SEMA4A*, *SEMA6D*), cytokine receptors (*IL1R1*, *IFNAR2*), c-type lectins^56^ (*MRC1*, *MRC2*), and solute transporters (*ABCA1*, *MMGT1*). Gene set over-representation analysis (ORA) of the 395 targets revealed enrichment in gene sets associated with cell–matrix interaction, axon and neural projections growth, cell mobility and migration, and solute transport across the plasma membrane (**Figure 2D**, **Figure S4A–E, Table S10, S11**). We observed moderate correlation between bulk RNA-seq expression of CAR-T targets in TCGA SARC and scRNA-seq expression (**Figure S3E**). For many targets like *CLMP, PDGFRB, NOTCH2,* and *GPC6*, TCGA SARC samples had the highest median expression compared with other cancer types in the TCGA Pan-Cancer Atlas (**Figure 2E**, **Figure S4F**).

To prioritize targets, we identified those essential for sarcoma cell line growth in vitro by analyzing publicly available CRISPR KO screening data from DepMap^57^ (**Figure 2F**, **Table S12**). Twenty-two genes were essential for at least one sarcoma cell line, with some being more selective for specific STS, like *TSPAN31* for LPS. Other genes were essential for multiple sarcoma cell lines, like *PTK2*, *VPS28*, and *UBE2D3*. We then studied the frequency of CNAs, gene fusions, and SNVs and small insertions/deletions (indels) involving each target, using publicly available DNA sequencing data (TCGA SARC^36^ and MSK-IMPACT, **Table S13, S14, S15, S16**)^58^. We found targets that were frequently amplified in specific STS histotypes in TCGA with high correlation between their CNA level and RNA expression, indicating that these events could have a driver effect^59,60^ (**Figure 2G**). For example, *TSPAN31* (*r* = 0.89) was amplified in 72.4% of DDLPS, and *FGFR1* (*r* = 0.68) was amplified in 12.5% of UPS. Integrating data from TCGA and MSK-IMPACT with the results of a search in the public Mitelman Database^61^, encompassing 52 STS types and 3283 unique fusions (**Figure 2H**, **Table S15**), we found 213 fusions across 20 STS histologies that involved at least one target. The CAR-T targets most altered by SNVs or indels included *MACF1* and *AXL* in UPS/MFH (10.2% and 6.1% in TCGA, **Figure 2I**, **Table S16**).

### Bispecific CAR-T targets in STS: ‘AND’ and ‘AND–NOT’ logic-gated CARs

To increase the surface antigen pool targetable by CAR-T cells in solid tumors, new bispecific CAR-T designs have been developed. Specifically, Boolean-logic ‘AND’-gated CAR-T cells^62^ like the logic-gated intracellular network (LINK) CAR-T^12^, and the synthetic NOTCH receptor (synNOTCH) CAR-T^63^ cause T-cell activation only when two separate antigens are recognized on target cells. Conversely, inhibitory CAR-T cells (iCARs)^64^ are an example of Boolean-logic ‘AND-NOT’-gated CAR-T cells, where an inhibitory domain prevents T-cell activation upon recognition of an antigen selectively expressed on normal but not on tumoral cells.

To discover targets for ‘AND’ CAR-T cells in STS, we sought to identify genes with high co-expression in STS cells and low to no co-expression in normal cells and tissues. (**Figure 3A, Figure S5A**). To this end, we measured co-expression of STS up-regulated genes in tumor cells at the single cell level using CS-CORE^65^ (**Figure S5B**), and in normal cells and tissues using the Genotype-Tissue Expression (GTEx)^66^ bulk RNA-seq and proteomics datasets, the Human Proteome Map (HMP)^67^ proteomics dataset, and the HPA scRNA-seq, bulk RNA-seq, and IHC datasets. We then compared tumor co-expression to normal tissue co-expression and calculated the *TS*_‘AND’_ score as a measure of targetability for each ‘AND’ logic-gated CAR-T gene pair, where pairs with *TS*_‘AND’_ ≥ 1 were those with better targetability. Although we found 3608 unique ‘AND’ CAR-T pairs (**Figure 3B, Table S17**), only 4 (0.1%) were common to all sarcoma histotypes, highlighting the histotype specificity of gene pairs as well as the challenge of identifying broadly applicable target pairs for ‘AND’ logic-gated CAR T cells.

**Figure 3.**
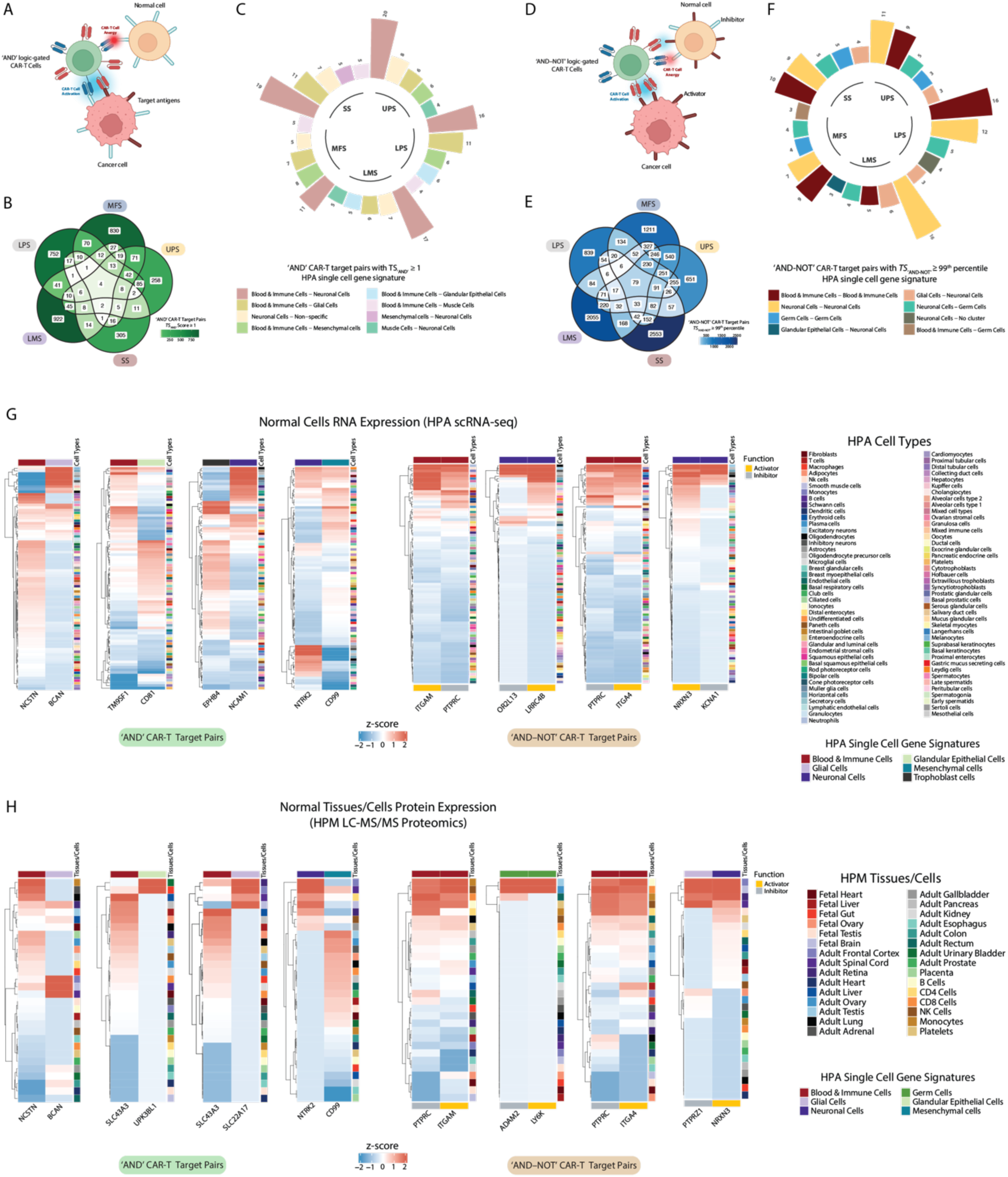
Bispecific CAR-T targets in STS: ‘AND’ and ‘AND–NOT’ logic-gated CARs. (A) Illustration of an ‘AND’ CAR-T cell activated by two different tumor antigens. (B) Venn diagram showing the number of ‘AND’ pairs in each STS. (C) Bar chart showing the top 5 most frequent ‘AND’ pairs by STS. Genes are classified based on their HPA single cell type signature. (D) Illustration of an ‘AND-NOT’ CAR-T cell activated by a tumor antigen, and inhibited by a normal cell antigen. (E) Venn diagram showing the number of shared and unique ‘AND-NOT’ pairs in each STS. (F) Bar chart showing the top 5 most frequent ‘AND-NOT’ pairs by STS. Genes are classified based on their HPA single cell signature, with the first indicating the activator and the second the inhibitor. (G, H) Heatmaps showing selected ‘AND’ and ‘AND-NOT’ pairs expression in the HPA scRNA-seq (G), and the HPM (H) datasets. Genes are annotated based on the HPA single-cell signature they belong to (top annotation), and their functional role as activators or inhibitors (bottom annotation, ‘AND-NOT’ pairs).

Most ‘AND’ CAR-T pairs consisted of genes with restricted expression in orthogonal normal cell types but shared expression in STS. Consistent with the mesenchymal cell origin of STS^68^, gene pairs from “Blood & Immune Cells” and “Neuronal Cells” were the most frequent in all STS histotypes reflecting the shared development of mesenchymal, hematopoietic, and neuronal cells from the embryonic mesoderm and neural crest^69^ (**Figure 3C, Figure S5C–F**). We previously observed expression of epithelial markers in malignant STS cells^17^, and gene pairs from both epithelial and mesenchymal cells were identified as targets for ‘AND’ CAR-T cells. These results highlight the potential for bispecific CAR-T cells to leverage the dysregulated differentiation of malignancies as a therapeutic target^70^.

For ‘AND-NOT’ CAR-T target pairs, the ideal combination would include an activator highly expressed on cancer cells, and an inhibitor with no expression on cancer cells but always co-expressed with the activator on normal cells (**Figure 3D, Figure S5A**). To find activator-inhibitor target pairs for ‘AND-NOT’ CAR-T cells, we used highly expressed STS up-regulated genes as activators and STS down-regulated genes with average scRNA-seq tumor expression ≤ 50^th^ percentile as inhibitors and measured their expression in normal cells and tissues. We combined the results of the co-expression analysis in each dataset to obtain a summative targetability measure, the *TS*_‘AND-NOT’_ score, with values raging from −1 to 1, where higher values indicate higher co-expression in normal tissues/cells. Overall, we found a total of 10,584 ‘AND-NOT’ CAR-T pairs across all sarcomas (**Figure 3E**, **Table S18**). Of these, 79 (0.7%) pairs were shared across all five STS, including *TSPAN6*–*CLDN17*, *PDGFC*–*GPR33*, and *IL17RD*–*ZNRF4.* Pairs with genes belonging to the same HPA single cell gene signatures were the most frequent, reflecting similar patterns of expression in normal cells (**Figure 3F**).

Considering the ‘AND’ and ‘AND-NOT’ CAR-T target pairs with high *TS* scores, again we found that most ‘AND’ pairs tend to have similar expression profiles in the same cells and tissues, while ‘AND-NOT’ pairs show opposite expression in the same tissues and cells, which was true both at the protein and RNA level (**Figure 3G-H**). Examples of ‘AND’ pairs include *NCSTN*– *BCAN* (“Blood & Immune Cells” – “Glial Cells”), *TM9SF1*–*CD81* (“Blood & Immune Cells” – “Glandular Epithelial Cells”), and *NTRK2*–*CD99* (“Neuronal Cells” – “Mesenchymal Cells”). Examples of ‘AND–NOT’ CAR-T target pairs include *PTPRC*–*ITGAM* (“Blood & Immune Cells”), *PTPRZ1*–*NRXN3* (“Glial Cells” – “Neuronal Cells”), and *ADAM2*–*LY6K* (“Germ Cells”). Many of the bispecific CAR-T targets were essential for STS cell line growth (**Figure S6A**) and/or were genetically altered in human STS (**Figure S6B-H**, **Table S19-S21**), suggesting their importance in STS oncogenesis.

### Targeting intracellular antigens in STS: TCR-T and PC-CAR-T cells

CAR-T cells target cell surface antigens but cannot directly recognize intra-cellular proteins, which constitutes a large fraction of cancer antigens^71^. To address this issue, TCR-T and peptide-centric CAR-T (PC-CAR-T)^72^ cells have been developed targeting peptides presented by the major histocompatibility complex (MHC) on tumor cells. Most peptides targeted by TCR-T cells tested in humans so far derive from cancer germline antigens (CGAs)^73^, which are highly expressed in cancer and reproductive organs, such as *MAGEA4* or *CTAG1B,* coding for the NY-ESO-1 protein^27,30^. SCAN-ACT uses DGE analysis to identify tumor cell up-regulated genes coding for either intra- or extracellular proteins, with normal tissue expression similar to CGAs targeted by TCR-T cells tested in humans to find tumor-specific genes with expression restricted to immune-privileged sites thus minimizing on-target off-tumor toxicity (**Figure 4A**, **Table S22**).

**Figure 4.**
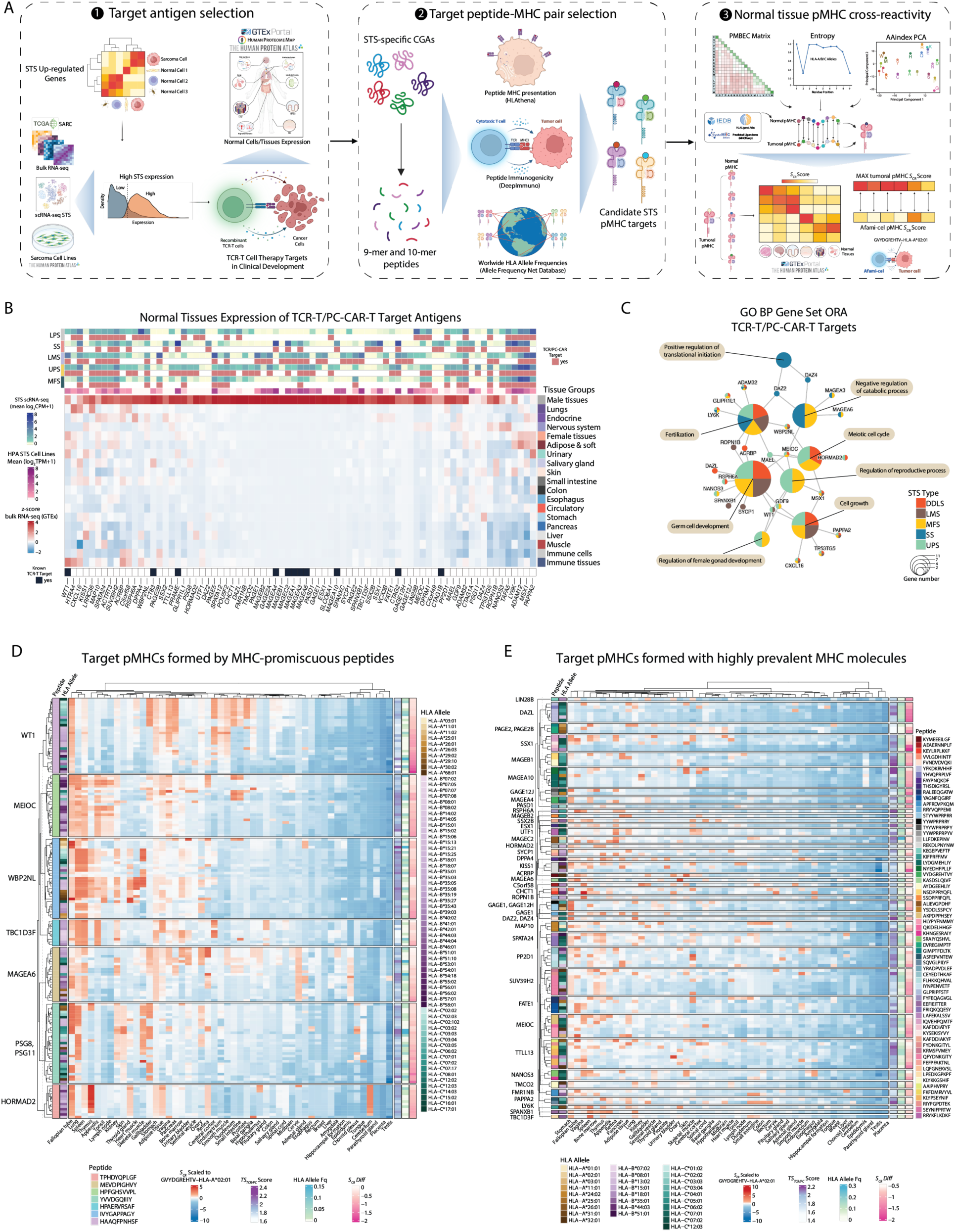
Targeting intracellular antigens in STS: TCR-T and PC-CAR-T cells. (A) TCR-T and PC-CAR-T pMHC target discovery pipeline. (B) Heatmap showing bulk RNA-seq GTEx expression for the 76 target antigens. Top annotation indicates predicted antigen in each STS, with average expression in STS scRNA-seq data and HPA STS cell lines. (C) Gene set ORA using GO BP terms for PC-CAR-T and TCR-T target antigens (FDR < 0.1, minimum and maximum gene set sizes of 2 and 500). (D, E) Heatmaps showing HPA/GTEx tissue-specific S_CR scores for target pMHCs scaled to the S_CR scores of afami-cel’s target, GVYDGREHTV–HLA-A*02:01, obtained as follows: (((S_CR^j (target)-S_CR^j (reference)))/σ_j) ×2 if S_CR^j (target)>S_CR^j (reference), where σ_j is the standard deviation of the S_CR values for the 5,020 target pMHCs in the jth tissue. In (D), pMHCs with S_CR Diff ≤0, and formed by peptides that form pMHCs with ≥10 different HLA alleles in either locus are shown. In (E), only pMHCs with S_CR Diff ≤ 10th percentile and formed by one of the top 10 most prevalent HLA alleles at either locus are shown.

SCAN-ACT identified 76 genes across five STS, with some showing uniformly high expression, and others more specific to one or few histologies (**Figure 4B**, **Table S23**). UPS had the highest number of targets (45), followed by LMS (35), LPS (28), SS and MFS (26), with 7 shared across all histologies (*ACTRT3*, *ADAM12*, *ADAM32*, *CXCL16*, *MAP10*, *MSX1*, and *SUV39H2*, **Figure S7A**). Overall, 34/76 antigens (44.7%) were known CGAs^73^, and gene set ORA identified a roles of the targets in gametogenesis and gonadal cells function (**Figure 4C**). Few of the targets were essential for STS cell line growth in vitro (**Figure S7B**), amplified in human STS (**Figure S7C**, **Table S24**), or involved in SNVs/Indels (**Figure S7E**, **Table S25**). However, 20 gene fusions in STS involved at least one target (**Figure S7D**).

Next, we identified a pool of *pMHCs* suitable as TCR-T or PC-CAR-T targets for each antigen. Specifically, we assessed the HLA presentation and immunogenicity potential of all the 9-mer and 10-mer peptides derived from each antigen, using HLAthena^74^ and DeepImmuno^75^, respectively, focusing on 9-mers and 10-mers as these account for more than 97.5% of HLA class I-bound peptides^76^. Also, we chose the top 10 most frequent HLA alleles in each major geographical area as reported in the Allele Frequency Net Database (AFND)^77^, accounting for 118 HLA alleles (**Table S26**). A major limitation of TCR-T cells is their ability to recognize peptides presented on one or rarely two HLA alleles, making them accessible to a small fraction of the population. To address this, we combined the HLAthena and DeepImmuno predictions with the average global HLA allele frequency of each *pMHC* to obtain a combined TCR-T/PC-CAR-T targetability score, *TS*_*TCR*/*PC*_. We retained the top 5,020 *pMHCs* formed by 348 peptides and 115 HLA alleles with 322 *pMHCs* shared between ≥ 1 antigen (**Table S27**). *ADAM32* had the highest number of targetable *pMHCs* (*n* =136), covering 64 HLA alleles (**Figure S7F**). We found a higher number of targetable *pMHCs* for HLA-C (*n* = 2,041), compared to HLA-B (*n* = 1,540) and HLA-A (*n* = 1,439). The top three alleles by number of targetable *pMHCs* were C*07:01 (*n* =168), C*06:02 (*n* =145), and C*07:02 (*n* =131), while C*04:01 had *pMHCs* with the highest median *TS*_*TCR*/*PC*_ (2.1) (**Figure S7G**).

Because TCR degeneracy allows T-cells to recognize different peptides presented on the same MHC molecule^78^, we designed a *pMHC* cross-reactivity prediction algorithm called PS-TRACT (<u>P</u>eptide <u>S</u>imilarity and <u>T</u>CR <u>R</u>eactivity <u>A</u>nalysis for <u>C</u>ellular <u>T</u>herapies). PS-TRACT measures the similarity of each tumoral peptide to peptides of the same length in the normal MHC ligandome (**Supplementary Data**) while considering the tissue-specific expression and likelihood of presentation of each cross-reactive peptide. Peptide similarity was measured in terms of amino acid physicochemical^79^ and HLA binding energy covariance similarity^80^, given that peptide sequence identity alone cannot fully predict T-cell cross-reactivity^81–83^. Also, because not all peptide residues bear the same importance in mediating TCR recognition^84^, we applied an entropy-based weighting, postulating that residues with higher entropy could be more relevant for *pMHC* TCR recognition^74^.

We validated PS-TRACT on experimentally observed cross-reactive peptides, including tumoral and microbial epitopes^85^, and the *TTN* peptide ESDPIVAQY which was found to be the off-tumor target in cases of fatal myocarditis in patients receiving TCR-T cells directed at the *MAGEA3* epitope EVDPIGHLY^86^ (**Figure S8A**). In each known cross-reactive pair, the target peptide was scored for similarity both against its known cross-reactive peptide and against all the normal peptides of same length binding to the same MHC molecule, used as controls. As expected, known cross-reactive peptides had peptide similarity scores (*S*_*sim*_) ≥ 99.9^th^ percentile (**Supplementary and Source Data**). We applied PS-TRACT to each STS *pMHC* and quantified cross-reactivity scores, *S*_*CR*_, for each tissue in the HPA/GTEx consensus database (**Supplementary Data**).

To test if PS-TRACT can predict tissue-specific TCR cross-reactivity, we applied it to *pMHC* targets for which TCR-T cells have been tested in clinical trials with safety outcome data available^11^ (**Figure S8B**, **Table S28**). We selected the *S*_*CR*_ score for the *pMHC* target of afami-cel, GVYDGREHTV–HLA-A*02:01, as the upper limit for our analyses given afami-cel’s promising safety profile in clinical trials^27,28^. We found that *pMHCs* for which cross-reactivity was observed in clinical trials^11^ had higher tissue-specific *S*_*CR*_ scores compared to GVYDGREHTV– HLA-A*02:01, as well as higher overall *S*_*CR*_ profile difference (*S*_*CR*_*Diff*) with GVYDGREHTV– HLA-A*02:01 when considering all tissues (**Figure S8C**). For example, the *MAGEA3*-derived *pMHC* KVAELVHFL–HLA-A*02:01, targeted by TCR-T cells causing two cases of fatal necrotizing leukoencephalopathy^87^, had a *S*_*CR*_ in the cerebral cortex which was 2.4 standard deviations above the *S*_*CR*_ of GVYDGREHTV–HLA-A*02:01 with the cross-reactive peptide identified as AVADLVHIV from *MCHR2,* which is highly expressed on neurons. Similarly, we observed a high cardiac *S*_*CR*_ score for EVDPIGHLY–HLA-A*01:01 driven by the known cross-reactive *TTN* epitope ESDPIVAQY^86^.

Using PS-TRACT, we found 1113 STS *pMHCs* with *S*_*CR*_*Diff* ≤ 0 across 182 peptides, 69 antigens, and 113 HLA alleles, and of these 74 were shared between ≥ 2 antigens (**Table S27**). We found several peptides predicted to form *pMHCs* with *S*_*CR*_*Diff* ≤ 0 with ≥ 10 different HLA alleles for either the class A, class B, or class C locus (**Figure 4D**). Such MHC-promiscuous peptides could be targetable with a PC-CAR-T recognizing the same peptide presented on different MHC molecules as long as the molecular surface of the resulting *pMHCs* is similar. For example, YVVDGQIIIY derived from both *PSG8* and *PSG11* bound to 17 class C (total allele frequency: 70%), 10 class B (total allele frequency: 8%), and 4 class A alleles (total allele frequency: 6%). Lastly, we found peptides forming *pMHCs* with very low risk of on-target off-tumor toxicity (*S*_*CR*_*Diff* ≤ 10^th^ percentile) with highly prevalent HLA alleles (**Figure 4E**). These peptides could be ideal TCR-T targets, given the increased TCR dependency on HLA restriction, and could be targeted in a larger fraction of the population due to the high prevalence of their HLA alleles. For example, YFKDKRVHHF from *DAZL* and QFYDNKGITY from *SUV39H2* were both targetable on C*07:01 (allele frequency: 14%).

### Validation of peptide-MHC binding for high-priority targets

Most *pMHC* binding prediction tools rely on datasets from mass spectrometric measurements of eluted peptide ligands (EL method), which require further verifications using the actual binding measurements (BA method) to eliminate false identifications^88^. Here, we adopted a recently developed empirical system, IMMUNE (Identifying Massive MHC Utilized Novel Epitopes), to validate the peptide-MHC binding of targets identified with PS-TRACT. IMMUNE employs a yeast display system enabling fast cloning as well as mammal-like post-translational modifications for the ex-vivo expression of complex HLA heterodimers^89,90^. Through anchorage to agglutinin (formed by Aga1p and Aga2p), a peptide-receptive format of HLA-A, B, or C non-covalent heterodimer can be assembled on the cell surface for antibody detection and peptide accommodation (**Figure 5A**). Using A*02:01, A*11:01, B*35:01, and C*12:03 as examples, we observed the entire heavy chain fusion protein represented by its C-terminal HA staining, the soluble form of β2m in association with the heavy chain, and the correctly folded heavy chain/β2m heterodimer represented by the HLA-A, B, C conformation-specific antibody staining, on the surface of correspondingly transformed yeast strains (**Figure S9A**). The successful detection of each component of the separately expressed heavy chain and β2m without any covalent linkers allowed further development of on-yeast peptide binding assays. Without the necessity to purify HLA proteins, the IMMUNE system substantially expedited BA evaluation and synergized with our computational prediction for rapid TCR-T or PC-CAR-T target discovery.

**Figure 5.**
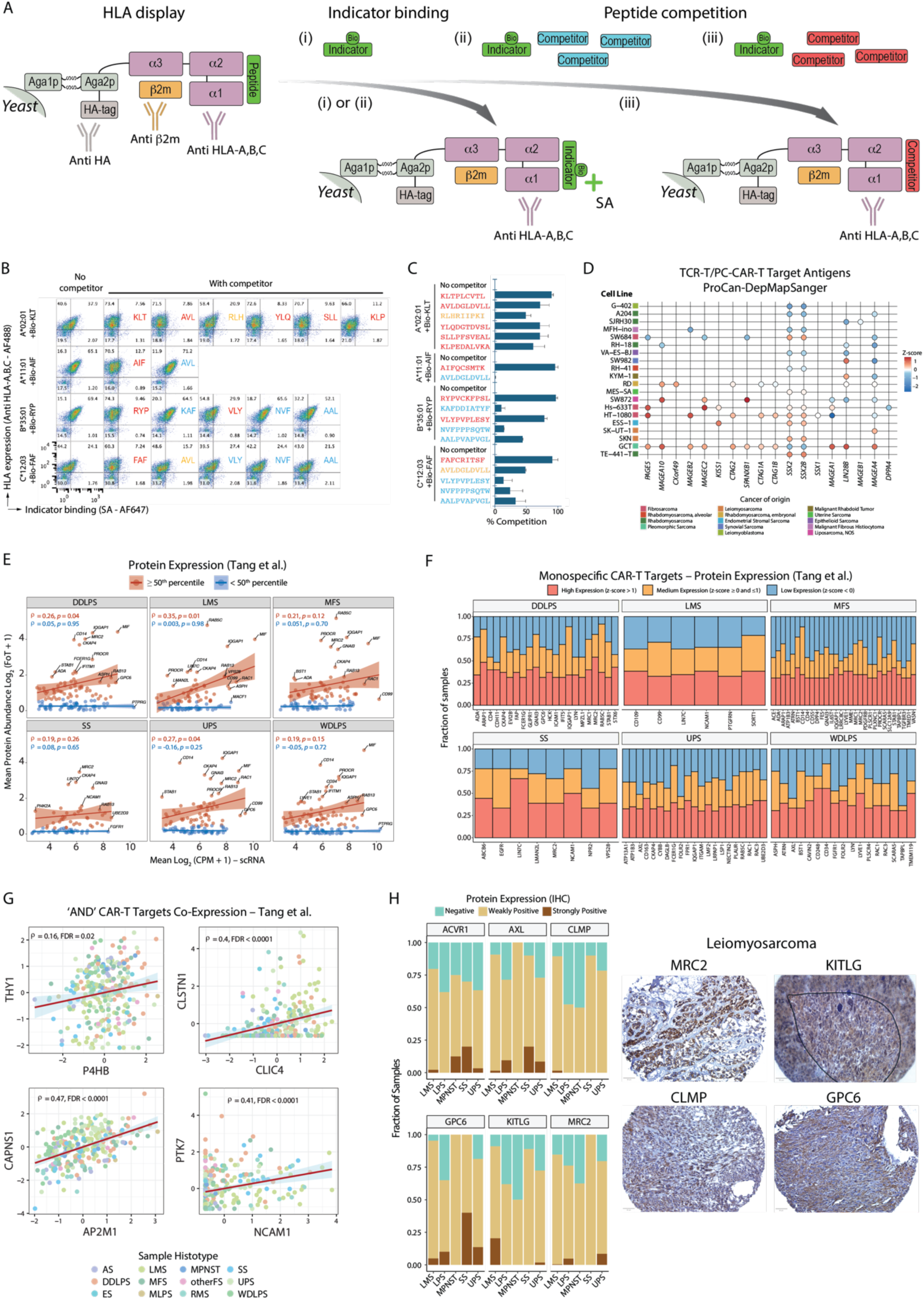
Target validation. (A) Vignette demonstrating the IMMUNE workflow for peptide-MHC binding assessment. (B) IMMUNE results for 14 pMHC targets identified through PS-TRACT, including KAFDDIATYF and RLHRIIPKI from SSXI, AVLDGLDVLL and VLYPVPLESY from PRAME, NVFPPPSQTW and YLQDGTDVSL from ADAM12, and SLLPFSVEAL, KLPEDALVKA and AALPVAPVGL from MSX1. The scatterplots report the mean fluorescence intensity (MFI) for the peptide-HLA binding indicator (x-axis) and for the HLA molecule tested (y-axis). Each peptide is identified by its first three amino acids. Strong binders are indicated in red, while moderate binders are shown in yellow, and weak binders in light blue. The KLT, AIF, RYP, and FAF peptides are used as positive controls for known strong binders for the HLA considered. (C) Bar chart showing the % Competition as measured with IMMUNE, for each tested pMHC target. Strong binders are indicated in red, moderate binders in yellow, and weak binders in light blue. (D) Dot plot showing protein expression in STS cell lines in ProCan-DepMapSanger. (E) Spearman correlation of scRNA-seq expression and protein abundance in the Tang et al. dataset. Only monospecific CAR-T targets for each histotype are plotted. LPS scRNA-seq samples were compared to DDLPS and WDLPS proteomics samples separately. (F) Bar charts showing the fraction of samples in each STS with high, medium, and low expression of selected monospecific CAR-T targets in the Tang et al. dataset. (G) Spearman correlation of protein expression in the Tang et al. dataset for ‘AND’ target pairs. (H) Protein expression by IHC in STS tissue micro-arrays. The bar chart shows the fraction of samples in each STS with negative, weakly positive, and strongly positive expression. Images show tissue sections of LMS samples scored as strongly positive for GPC6, MRC2, CLMP, and KITLG. MPNST: malignant peripheral nerve sheath tumor; MLPS: mucinous liposarcoma; others: other fibrosarcoma; ES: Ewing sarcoma; RMS: rhabdomyosarcoma; AS: angiosarcoma.

By comprehensively analyzing the epitope repertoire of each HLA in the immune epitope database (IEDB), we designed indicator peptides that could be biotinylated without affecting HLA binding. Co-staining of HLA molecules and the exogenously loaded biotinylated indicators confirmed specific binding on the surface of each yeast strain (**Figure S9B**). The capability of IMMUNE to display peptide-receptive HLA mono-alleles represents an ideal system for competitive binding analysis against the indicator. Based on overall *S*_*CR*_ profile and *TS*_*TCR*/*PC*_, we selected 9 peptides forming 14 *pMHC* targets for testing. All test peptides bound A*02:01, including RLHRIIPKI from SSX (**Figure 5B**, **5C**). However, other than AVLDGLDVLL and VLYPVPLESY from PRAME, most peptides showed low to moderately competitive binding for B*35:01 and C*12:03. These results suggest that IMMUNE can be utilized to prioritize pMHCs for targeting.

### Target expression in STS Proteomics Datasets

Next, we analyzed target expression in STS clinical proteomics databases^91,92^, sarcoma cell lines^93,94^, and STS tissue micro-arrays (TMAs). Since none of the TCR-T/PC-CAR-T target antigens were measured in the clinical STS proteomics datasets, we analyzed their expression in STS cell lines, as reported in the Cancer Cell Line Encyclopedia (CCLE)^93^ and in the ProCan-DepMapSanger^94^ repositories. Overall, 33/76 (43.4%) targets were expressed in 21 cell lines spanning 13 STS histotypes (**Figure 5D, Figure S9C**). Of all the targets assessable for expression, *SSX2*/*SSX2B*, *LIN28B*, and *MAGEA4* had the most cell lines with medium-high expression in ProCan-DepMapSanger at 32%, 18% and 14%, respectively. In the Tang et al. STS clinical proteomics dataset^91^, we found overall good correlation between scRNA-seq and protein expression for the STS up-regulated surfaceome genes (**Figure S9D**, **S9E**). This correlation was consistent across histotypes when considering the monospecific CAR-T targets, particularly only those with protein expression ≥ 50^th^ percentile (**Figure 5E**, **Figure S9F**). We found that many targets had high expression in a large fraction of samples (**Figure 5F, Figure S10A, S10C**). Similar results were observed in the Burns et al. dataset (**Figure S10B**).

For the ‘AND’ CAR-T targets with measurable expression of both genes in the Tang et al. dataset, we measured co-expression across all the STS samples (**Table S29**). Of the 1207 ‘AND’ pairs, 274 showed positive correlation in protein expression between the two targets across all STS samples (**Figure 5G**). Of the 438 ‘AND’ CAR-T pairs found in the Burns et al. dataset, 79 had positive correlation in expression across all samples (**Table S29**). Notably, we were unable to distinguish tumor and normal cell protein expression from the bulk analysis, which may confound the correlation.

Lastly, we measured expression of selected targets using in-house TMAs of LMS, SS, MFS, UPS, and MPNST samples (**Figure 5H**). *GPC6* was expressed in 100% (*n* = 10) of SS, 81.4% (*n* = 59) of UPS, and 95% (*n* = 537) of LMS samples, similarly to *MRC2* which was expressed in all SS samples, 84.6% LMS, 79.7% UPS, and 76.1% LPS (*n* = 21) samples. LMS had the highest proportion of positive samples for *KITLG* (91.0%), while SS had the highest number of samples positive for *CLMP* (90.0%), followed by LMS (89.4%). These antigens were selected for IHC assessment due to their overall high STS expression and low normal expression profile.

### Target Prioritization

We calculated a Target Priority Index (TPI) for each cell therapy target class, integrating normal cell/tissue expression with functional, genomics, and proteomics STS data, while considering single-cell tumor expression homogeneity and existing immunotherapeutic agents (**Table S30-S33**). We defined scRNA-seq expression homogeneity as the product of average expression and fraction of cells with non-zero expression (*H*_*RNA*_ score). For monospecific and bispecific CAR-T targets, we used the average *H*_*RNA*_ score of *CD276* across all histotypes as the homogeneity reference, and the average fraction of samples in the Tang et al. proteomics dataset with *CD276* z-score ≥ 0 as the protein expression reference because *CD276* has been tested as a CAR-T target in STS^95^. Using LPS as an example (**Figure 6A**, **6B**), notable monospecific CAR-T targets in the top 100 *TPI*_*CAR*_ included *GPC6* (high protein expression and *H*_*RNA*_), *FGFR1* (essential in LPS cell lines and with ADCs in clinical development^96^), and *TSPAN31* (essential for LPS cell lines and involved by gene fusions and amplifications in LPS).

**Figure 6.**
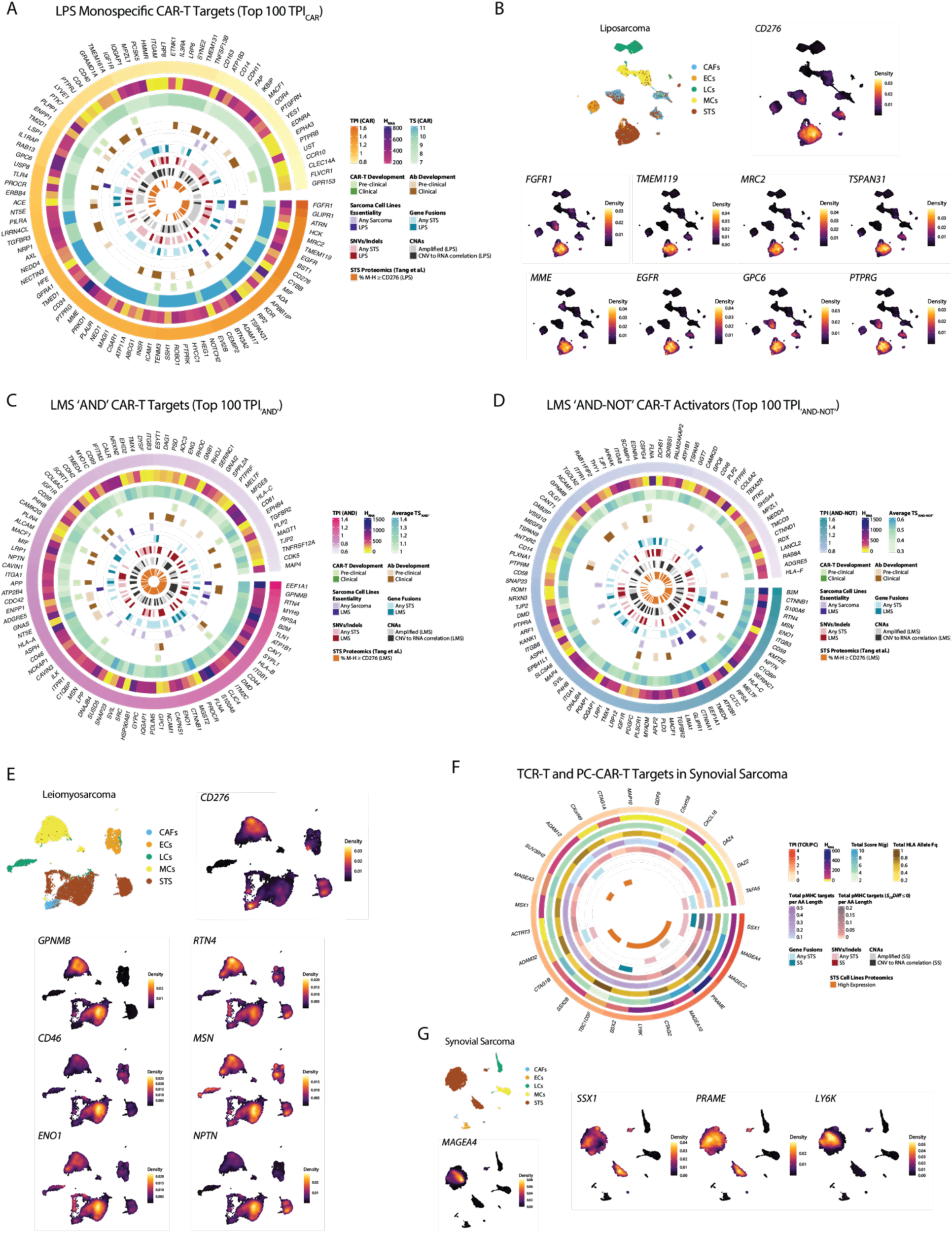
Integrated target prioritization. (A, C, D) Circular heatmaps showing the top 100 CAR-T targets ranked by TPI for LPS (A, monospecific CAR-T targets), and LMS (C, D, bispecific CAR-T targets). (B, E, G) UMAP plots showing expression density of selected monospecific CAR-T targets in LPS (B), bispecific CAR-T targets in LMS (E), and TCR-T/PC-CAR-T target antigens in SS (G). In one UMAP plot per panel, cells are colored based on identity. (F) Circular heatmap showing TPI scoring for TCR-T and PC-CAR-T target antigens in SS.

For bispecific CAR-T targets, we integrated the average *TS*_′*AND*′_ and *TS*_′*AND*-*N*0*T*′_ for all the targetable pairs in which each gene appears into the TPI, prioritizing genes suitable both as ‘AND’ CAR-T targets and ‘AND-NOT’ CAR-T activators. In LMS as an example (**Figure 6C**, **6D, 6E**), notable targets with antibodies developed clinically or pre-clinically include *GPNMB*^97^, *RTN4*^98^ and *CD46*^99^. Considering all the monospecific and bi-specific CAR-T targets identified, 69 had a clinically developed monoclonal antibody (mAb), antibody-drug conjugate (ADC), single-chain variable fragments (scFv), fusion protein, nanobody, or fragment antigen binding region (Fab), while 36 had at least a mAb or ADC in pre-clinical development. Also, of all the CAR-T targets described, 26 and 32 have been previously tested as CAR-T targets either clinically or pre-clinically, respectively.

Finally, we calculated a TPI for TCR-T and PC-CAR-T target antigens, using *MAGEA4 H*_*RNA*_ in SS as the homogeneity reference, given afami-cel’s efficacy in SS. In addition, we considered protein expression in STS cell lines, the total number of target *pMHCs*, and the HLA allele frequency per antigen (**Table S33**). Top targets included *PRAME* and *LY6K*, which have more homogenous expression than *MAGEA4* in SS and a higher total HLA allele frequency. In addition, *SSX1* is recurrently involved in pathogenic fusions in SS and has a higher predicted number of targetable *pMHCs* than *MAGEA4* (**Figure 6F**, **6G**). Given the large number of targets predicted by SCAN-ACT, the TPI will be helpful to identify the most promising tumor-specific antigens for further testing.

## Discussion

The major bottleneck for development of ACTs in solid tumors continues to be the lack of tumor-specific antigens with low expression on normal cells^100,101^. Differently from hematological malignancies, solid tumors lack lineage-specific antigens with homogeneous high-density expression. Also, it remains challenging to ascertain an acceptable level of target expression in normal tissues. To address this, we developed SCAN-ACT, a computational pipeline leveraging scRNA-seq to discover targets for TCR-T cells, and monospecific, bispecific, and peptide-centric CAR-T cells, targeting both intracellular and surface proteins.

We utilized SCAN-ACT to screen for targets in STS, a disease with limited therapeutic options where ACT has proven potential. The characteristic heterogeneity of STS is also an ideal small-scale representation of solid tumor heterogeneity, thus providing a high-bar test case for SCAN-ACT. We screened sarcoma-specific antigens using single-cell DGE analysis between STS cells, non-tumoral cells in the TME, and normal cells across 30 human tissues. scRNA-seq can identify genes that are more selectively expressed in tumoral cells compared to non-tumoral cells in the TME, as opposed to bulk transcriptomics and proteomics^17,18^. Additionally, scRNA-seq allows measurement of target expression heterogeneity, a crucial barrier to ACT efficacy in solid tumors^1^, and the availability of normal tissue scRNA-seq data allows for better estimation of the on-target off-tumor toxicity risk^41,42^. Previous studies demonstrated that target expression even on rare cellular populations in vital organs can mediate CAR-T cell toxicity^102^, and scRNA-seq may better predict such off-tumor effects than bulk RNA-seq^103^.

We extensively validated the off-tumor toxicity risk of the identified targets using publicly available multiomics data in normal cells and tissues, including scRNA-seq, bulk RNA-seq, IHC, and proteomics, with targets already tested in humans as reference. Through this approach, we identified targets with lower predicted risk of on-target off-tumor toxicity than those currently available. We assessed the relevance of the identified targets to STS biology at the transcriptomic, genomic, and proteomic level, as well as functionally, through CRISPR KO screens. This highlighted *TSPAN31* on chromosome 12, which is frequently involved by CNAs in DDLPS and co-amplified with *MDM2* in both WDLPS and DDLPS^104^, indicating a potential oncogenic driver role in LPS.

We found targets not previously studied in STS like *GPC6,* a member of the glypican family which is pathogenic in sarcomas^105^, or *CLMP,* described as tumor suppressor in colon cancer^106^. We show targetable pathways and protein families in STS, including semaphorins and plexins that are physiologically involved in neural development^107^ and poorly studied in STS^108^; the NOTCH pathway, previously targeted in STS^109,110^; the EPH/Ephrin system, described as pathogenic in sarcomas^111^ and physiologically involved in neuron migration, axon guidance, and vascular remodeling^112^; and the WNT/β-Catenin pathway, implicated in sarcomagenesis and previously targeted in STS^113^. Also, we identify stromal antigens, previously targeted to manipulate the TME, as tumor cell intrinsic in STS like *FAP*^43^, *FOLR2*^114^, or *CD248*^115^.

Here we also provide a tool for screening safe and effective *pMHC* targets for ACT in solid tumors. We expand the pool of known CGAs targetable in STS^116^, identifying antigens not previously described such as *SUV39H2* or *MEIOC*, while confirming the role of known CGAs like *PRAME*, *SSX1*, and those of the MAGE and GAGE families^116^. Targetable *pMHCs* were identified using computational tools measuring peptide HLA presentation and immunogenicity. Despite being an in silico analysis, this approach prioritizes peptides likely to trigger cytotoxic T-cell responses while maximizing the number of patients who could benefit from targeting a specific *pMHC*. Almost all TCR-T cells tested in humans so far recognize peptides presented on HLA-A*02:01^11^, which is highly prevalent in Caucasians, and less so in African Americans and Asians and Pacific Islanders^77^. This creates equity and generalizability issues for HLA-restricted ACTs^32^. To address this, we integrated allele frequency into *pMHC* target prioritization, including the most frequent HLA alleles from each major geographical area^77^. We curated a normal MHC ligandome database of 46×10^6^ *pMHCs*, including 630,022 experimentally validated peptides, and developed PS-TRACT, a tool for *pMHC* T-cell cross-reactivity assessment. PS-TRACT overcomes limitations of current methods^72,117^, allowing predictions for 9-mer and 10-mer peptides over 223 HLA alleles and measuring peptide similarity through biophysical properties while applying an entropy-based weighting that prioritizes amino acids critical for TCR recognition. By using a large pool of experimentally observed and computationally predicted normal *pMHCs*, PS-TRACT provides a multi-tissue cross-reactivity risk estimation for each tumoral *pMHC*. Finally, we experimentally measured HLA binding of selected target peptides using a novel yeast display technology, IMMUNE, especially critical for validation of peptide-binding to less common HLA alleles where in silico predictions may be inaccurate.

Compared to prior CAR-T target discovery efforts that either performed DGE analysis between tumoral and normal bulk RNA-seq samples^14,15^ or analyzed patterns of protein expression in tumor samples^45^ or cancer cell lines^16^, we leveraged single-cell DGE analysis with a large cohort of non-tumoral reference cells, allowing a more precise screening for tumor-specific genes.

Unlike prior studies on bispecific CAR-T targets^14,16,45^, we defined tumoral target co-expression at the single cell level and directly measured co-expression in normal cells and tissues using correlation analyses, including normal tissue scRNA-seq and proteomics data. Lastly, compared to prior scRNA-seq studies for CAR-T target discovery^18^, we analyzed a larger number of tumor samples, included more normal reference cells, applied robust single-cell DGE analyses less prone to false discovery^118^, validated the identified targets with genomics, proteomics, transcriptomics, and functional assays in STS, and identified TCR-T/PC-CAR-T and Boolean-logic gated CAR-T cell targets.

In summary, we developed a comprehensive tool for the identification of ACT targets in solid tumors enabling the targeting of both intracellular and membrane proteins. By applying the tool for STS, we nominate and prioritize novel putative targets to expand and accelerate the early success of ACT in this disease. Our pipeline and sarcoma ACT target database are freely accessible for academic use at <u>scanact.stanford.edu</u>.

## Acknowledgments

We thank the patients and families who participated in this study. This work was supported by the My Blue Dots organization (E.J.M.) and the Tad and Diane Taube Family Foundation (M.v.R., E.J.M.). This work used the UCLA Technology Center for Genomics and Bioinformatics and the UCLA Translational Pathology Core Lab. The schematics in Figure 1, 2, 3, 4, and Figure S5 were created with BioRender.com.

## Author contributions

S.T., A.K., and E.J.M. were responsible for conceptualization. S.T., A.K., and E.J.M. were responsible for the methodology. S.T. and E.J.M. were responsible for software. S.T., A.P., A.K., and E.J.M were responsible for formal analysis. S.T., A.S., A.K., and E.J.M. were responsible for data curation. S.T., A.K., and E.J.M. were responsible for writing the original draft. W.J. and A.K. were responsible for the IMMUNE validation and related methods and results. S.T., A.P., A.S., S.V., M.P., N.Q.B, K.N.G., M.v.R., P.H., D.G., S.D., W.J., A.K., and E.J.M. were responsible for review and editing. S.T. A.P., S.A., P.T., S.A., D.G. and E.J.M. were responsible for visualization. P.H., M.v.R., A.K., and E.J.M. were responsible for supervision. A.K. and E.J.M. were responsible for funding acquisition.

## Data and Code Availability

The processed scRNA-seq data new to this paper have been deposited to GEO with accession code GSE279852 (reviewer token: qbejwckcjnkvbgv). The raw scRNA-seq data new to this project is available on SRA with project code PRJNA117421. The previously published scRNA-seq datasets reanalyzed in this project are available in GEO with accession codes GSE212527 and GSE131309. The SCAN-ACT pipeline and code used to create the images in the manuscript are available on Github at https://github.com/StefTesta/SCAN_ACT and as supplementary material. The SCAN-ACT pipeline and the CellSARCTx interactive interface are fully accessible for academic use at https://scanact.stanford.edu.

## Supplemental Information

Document S1. Figures S1-S10.

Document S2. Excel File containing Supplementary Tables S1-S34.

## Methods

### Human subjects

All samples analyzed in this study were collected with informed consent from subjects enrolled on protocols approved by the Institutional Review Boards at Stanford University and University of California Los Angeles.

### Single cell RNA sequencing of STS samples

STS belonging to the following STS histotypes were included in the study: LMS, MFS, UPS, DDLPS, WDLPS, and SS. Pathological diagnosis was confirmed through intuitional pathologist review for the samples analyzed at Stanford University and University of California Los Angeles, and as reported in the original study for the SS samples^33^. For the MFS2 sample, initial tumor biopsy was diagnosed as MFS, while final surgical specimen was classified as UPS. Due to the ambiguity of diagnosis, and the lower representation of MFS samples in our cohort, we decided to analyze this sample as MFS. Also, the UPS6 sample was classified as Spindle Cell Sarcoma versus UPS and was hence analyzed with the other UPS samples in the study.

For samples processed at Stanford University, fresh tumors were minced and enzymatically dissociated using the Human Tumor Dissociation Kit (Miltenyi Biotec) using a GentleMACS Octo Dissociator with heaters (Miltenyi Biotec) following the manufacturer’s instructions. The dissociated cell suspension was then applied to a 70 um MACS SmartStrainer (130-098-462; Miltenyi Biotec). The quality of the suspension was checked by generating a hematoxylin and eosin (H&E) smear and analyzed using a TC10 automated cell counter (BioRad). Samples were further processed by the Stanford Genomics Shared Resource or the using the Chromium 3’ Gene Expression Solution v3.1 (10x Genomics) per the manufacturer’s instructions and sequenced on a NovaSeq 6000 (Illumina). For samples processed at the University of California Los Angeles (MFS1/BO112_sarc04, UPS1/BO112_sarc10, UPS2/BO112_sarc11, UPS3/BO112_sarc13, UPS4/BO112_sarc15, UPS5/BO112_sarc17, LPS1/BO112_sarc07, LPS2/BO112_sarc09, LPS2sx/BO112_sarc09_sx, LPS3sx/BO112_sarc16_sx), these were dissociated by treatment with collagenase D (100mg/mL) and DNase I (100mg/mL) using a 100 *μ*m cell strainer, following the manufacturer’s instructions. After digestion, RBC lysis was performed by adding 1mL of BD lysing buffer (BD Biosciences) and incubating for 5 mins. Then, dead cells were eliminated using a dead cell removal kit (Miltenyi Biotec). Cells were counted using a Countess II cell counter (Thermo Scientific). Samples were then further processed by the UCLA Technology Center for Genomics & Bioinformatics (TCGB) using the Chromium 3’ Gene Expression Solution v3.1 (10x Genomics) per the manufacturer’s instructions and sequenced on a NexSeq500 High Output instrument (Illumina).

### Pre-processing of scRNA-seq STS samples

Sample demultiplexing, read alignment to the GRCh38 human reference genome, and generation of feature-barcode matrices were performed using CellRanger v6.0.0 (10x Genomics). Raw data was pre-processed in R (version 4.2.2.)^119^ using the Seurat package (version 4.3.0)^120^. Quality control was performed separately for each sample and consisted of excluding cells with number of expressed genes < 200 and percentage of mitochondrial genes > 15%. We then identified and excluded doublets using the R package DoubletFinder^121^ (version 2.0.3) applying the standard workflow as described by the developer.

### Sample Integration and cell annotation

Log-normalization was performed independently for each sample using the Seurat function NormalizeData. After this step, the Seurat objects for each independent sample were merged preserving the individual normalization of the data. Sample integration and clustering were performed separately for each histotype, accounting for batch effects and differences in sequencing technologies. The top 2000 highly variable genes of the merged and normalized expression matrices were identified using the function FindVariableFeatures, and then centered and scaled using the ScaleData function. Principal component analysis was then run using the top 2000 highly variable genes previously identified. To integrate multiple samples, the harmony R package^122^ (version 0.1.0) was employed for batch correction. Harmony was ran using both the sample of origin and the sequencing technology variables as arguments of “group.by.vars”, and based on the first 25 PCA dimensions previously identified. UMAP embeddings were then obtained using the first 25 Harmony dimensions, and clusters were obtained by calculating the k-nearest neighbors (k-NN) and a shared nearest-neighbor graph using the Louvain algorithm implemented in the FindClusters function of the Seurat package with a resolution of 1.0. For the LPS, MFS, UPS, and LMS samples, endothelial cells, myeloid cells, lymphoid cells, and CAFs were annotated using the differentially expressed genes discovered with the FindMarkers function in Seurat, which was run with the default parameters except for only.pos = T and min.pct = 0.25. Canonical cell identity markers obtained from the literature were used for cell type annotations. A list of the marker genes for each cell type in our samples is provided in **Table S2**. A small number of cells could not be uniquely assigned to a specific cell identity and were removed from further downstream analysis. UMAP plots were generated with Seurat and scCustomize^123^. Density plots were generated with the R package Nebulosa^124^.

### Public scRNA-seq Datasets

#### Synovial Sarcoma Samples

Previously published synovial sarcoma 10x Genomics and SMART-seq2 scRNA-seq data from Jerby-Arnon et al.^33^ were downloaded from GEO (GSE131309). After applying quality control filtering as described above, the Seurat objects for each sample were integrated using the harmony^122^ pipeline as previously described. Author-provided annotations were used to classify cells as sarcoma cells, endothelial cells, lymphoid cells, myeloid cells, and CAFs.

#### Normal Tissue scRNA-seq Datasets

A list of normal tissue scRNA-seq samples of reference used for differential gene expression was obtained from the HPA website (https://www.proteinatlas.org/about/assays+annotation#singlecell_rna). If for any of the reference datasets listed on the HPA dataset, only one sample was available, additional data was obtained from either Tabula Sapiens^41^ or from the Human Cell Atlas^42^. For each dataset, the raw count matrices were downloaded from the publication-specific repositories as follows: Adipose Tissue (GEO: accession code GSE155960); Bone Marrow (GEO: accession codes GSE159929-GSM4850584, and Tabula Sapiens: https://cellxgene.cziscience.com/e/4f1555bc-4664-46c3-a606-78d34dd10d92.cxg/); Brain (The Allen Brain Map: https://portal.brain-map.org/atlases-and-data/rnaseq/human-m1-10x); Breast (GEO: accession code GSE164898, except for D6-D10 samples); bronchus (figshare: accession code fig11981034); Colon (GEO: accession code GSE116222); Endometrium (GEO: accession code GSE111976); Esophagus (GEO: accession codes GSE159929-GSM4850580 and HCA: https://cellxgene.cziscience.com/e/b07fb54c-d7ad-4995-8bb0-8f3d8611cabe.cxg/); Eye (GEO: accession code GSE137537); Fallopian Tube (GEO: accession code GSE178101, excluding hydrosalpinx samples); Heart Muscle (GEO: accession code GSE109816 and Tabula Sapiens: https://cellxgene.cziscience.com/e/e6a11140-2545-46bc-929e-da243eed2cae.cxg/); Kidney (GEO: accession code GSE131685); Liver (GEO: accession code GSE115469); Lung (Tabula Sapiens: https://cellxgene.cziscience.com/e/0d2ee4ac-05ee-40b2-afb6-ebb584caa867.cxg/); Lymph Node (GEO: accession codes GSE159929-GSM4850583 and Tabula Sapiens: https://cellxgene.cziscience.com/e/18eb630b-a754-4111-8cd4-c24ec80aa5ec.cxg/); Ovary (EMBL-EBI: accession code E-MTAB-8381); Pancreas (GEO: accession code GSE131886); PBMC (GEO: accession code GSE112845 and Tabula Sapiens: https://cellxgene.cziscience.com/e/983d5ec9-40e8-4512-9e65-a572a9c486cb.cxg/); Placenta (EMBL-EBI: accession code E-MTAB-6701); Prostate (Tabula Sapiens: https://cellxgene.cziscience.com/e/d77ec7d6-ef2e-49d6-9e79-05b7f8881484.cxg/); Rectum (GEO: accession code GSE125970); Salivary Gland (Tabula Sapiens: https://cellxgene.cziscience.com/e/f01bdd17-4902-40f5-86e3-240d66dd2587.cxg/); Skeletal Muscle (GEO: accession code GSE143704); Skin (GEO: accession code GSE130973); Small Intestine (GEO: accession code GSE125970); Spleen (GEO: accession codes GSE159929-GSM4850589 and Tabula Sapiens: https://cellxgene.cziscience.com/e/cee11228-9f0b-4e57-afe2-cfe15ee56312.cxg/); Thymus (Tabula Sapiens: https://cellxgene.cziscience.com/e/0ced5e76-6040-47ff-8a72-93847965afc0.cxg/); Testis (GEO: accession code GSE120508); Tongue (Tabula Sapiens: https://cellxgene.cziscience.com/e/55cf0ea3-9d2b-4294-871e-bb4b49a79fc7.cxg/); Vascular (Tabula Sapiens: https://cellxgene.cziscience.com/e/a2d4d33e-4c62-4361-b80a-9be53d2e50e8.cxg/). After download, each dataset was pre-processed in a similar fashion as our own STS samples. The pre-processed samples for a given tissue were then integrated with the harmony pipeline^122^ using the sample variable, and for the SS samples the sequency technology variable as well, for batch correction, as previously described. Then, unsupervised clustering using the Louvain algorithm with a resolution of 1 was applied for all datasets.

Exceptions to this workflow include the brain dataset where the dimensionality reduction and clustering from HPA were used, as well as the endometrium and the bronchus datasets where the author-provided clusters and annotations were used. Author-provided annotations were used whenever possible, and if these were not available, the package SingleR^125^ and common gene markers were used for cluster cell-type annotation, where each cluster was assigned to its most frequent cell type. In SingleR^125^, the Encode^126^ and Blueprint epigenomics^127^ datasets were used for annotation of all datasets except for the pancreas for which the scRNA-seq reference data from Baron et al.^128^ was used instead.

### Identification of tumor cells based on InferCNV

CNVs were calculated using the R package inferCNV^35^. Each sample was independently analyzed using as reference cells the previously annotated CAFs, endothelial cells, myeloid cells, and lymphoid cells from the TME. These were identified based on expression of known marker genes available from the literature as described above. The following parameters were used to run the inferCNV function: analysis_mode=“subclusters”; tumor_subcluster_partition_method=“qnorm”; cluster_by_groups= FALSE; k_obs_groups = 3, denoise=T, HMM=TRUE. All the other parameters were set as default. Only for UPS k_obs_groups was set to 2 since this resulted in better overlap with the classification based on comparison to bulk sarcoma tumors (see below). To define a cell as potentially malignant, we used the CNA states across all genes calculated using a hidden Markov model (HMM) as implemented in the inferCNV package. To reduce the number of false positive CNVs identified, a Bayesian latent mixture model was applied with a default p-value threshold of 0.1 for all samples. For each gene, inferCNV reports a CNA state as one of the following: state 1 corresponding to a complete allele loss, state 2 corresponding to loss of one allele, state 3 corresponding to the neutral state with both allelic copies intact, state 4 corresponding to gain of one allele, state 5 corresponding to gain of 2 alleles, and state 6 corresponding to gain of more than two alleles. For each sample, CNA states of 6 in the output data matrix obtained in the 19^th^ step of the inferCNV workflow were first converted to a state of 5, and then the CNA states were normalized to values ranging from −1 to +1 and centered on 0 using the following formula: (CNA State − 3) / 2. The matrix values were then squared to give the same weight in the analysis to copy gains and copy losses. Then we calculated the overall CNA level of each cell. For each sample, we identified, the CD45^−^ cells with the top 10% CNA level. Cells with normalized PTPRC RNA count = 0 were defined as CD45^−^. The average CNA level for these cells was defined as the CNA profile of the tumor. A spearman correlation coefficient was then calculated for each cell between its overall CNA level and the CNA profile of the tumor. Cells whose spearman correlation coefficient was above the 25^th^ percentile of all the cells in each sample were considered to have detected CNAs and labeled as potentially malignant. The output of the inferCNV analysis used to identify tumor cells is provided in the supplementary data.

### Bulk sarcoma similarity score

Bulk RNA-seq of STS samples from TCGA were downloaded (Tatlow and Piccolo, 2016) and scaled to transcripts per million (TPM) as previously described and used as the bulk tumor references. Cells from scRNA-seq data available from Jerby-Arnon et al.^33^ that were annotated as normal fibroblasts, macrophages, and endothelial cells were used as the normal reference cells. First, the sarcoma histotypes of reference were extracted from TCGA, then the expression matrices for both the bulk RNA-seq sarcoma samples (TPM) and the scRNA-seq samples (raw UMI counts) were subsampled to include only genes in common between each dataset to compare. Then, two spearman correlation coefficients were calculated for a given cell of a scRNA-seq sample, respectively comparing its gene expression profile to the average expression profile of all the bulk RNA-seq tumor samples of reference (*ρ* Tumor) and to the average gene expression profile of all the normal cells in the scRNA-seq dataset used as normal control (*ρ* Normal). Based on these, we defined the similarity score as *ρ* Tumor – *ρ* Normal. Cells with similarity score > 0 were classified as potentially malignant. The InferCNV and similarity to bulk analyses were combined so that a cell was labeled as malignant if it was CD45^−^ (PTPRC RNA count = 0) and had either detected CNV or a similarity score > 0. Cells that could not be classified into a specific cell type based on any of the above methods were labeled as “unknown” and excluded from further downstream analysis.

### Differential Expression Analysis

To limit false discovery of differentially expressed genes in STS using scRNA-seq^118^, we used a pseudobulk DGE method, limma-voom^37^, which has been shown to be statistically more robust and less prone to false discovery than other methods when applied to single cell transcriptomics. Also, for the STS cells versus non-tumoral cells in TME, we complemented the limma-voom results with a new differential distribution testing method, distinct^38^, which was specifically designed for DGE using scRNA-seq.

#### Pseudobulk Limma-Voom

Sarcoma cells and normal cells were identified as above. For each STS histotype, the tumor cells were compared to endothelial cells, myeloid cells, lymphoid cells, and CAFs in the TME, as well as with each normal cell cluster of the normal tissues scRNA-seq samples of reference. For each comparison, first a single integrated Seurat object was created and from this, the raw non-normalized mRNA count matrix was extracted. To minimize variability in DE analysis and reduce false discoveries, the synovial sarcoma samples analyzed with plate sequencing technology and those analyzed with droplet-based technology were tested separately. Also, when comparing tumor samples to normal samples from HPA-TS-HCA, only genes that were present in both gene expression matrices were included in the differential expression test, to avoid identification of falsely up-regulated or down-regulated genes. Based on the information in the Seurat object metadata, the total mRNA counts for each gene were calculated by summing across all the cells of a certain cell type belonging to a specific sample, obtaining a unique pseudobulk for each cell type belonging to a given sample in the analysis. The pseudobulk gene expression matrix was then converted to a DGEList object using the DGEList function in the edgeR^129^ package (version 3.99.4), and subsequently normalized using the calcNormFactors function. Before voom transformation using the voom function from the R package limma^37^ (version 3.57.11), lowly expressed genes were filtered from the expression matrix, using a cutoff of 1 CPM. Counts data was then voom-transformed and a limma workflow was implemented using the sample variable as the blocking variable to control for batch effect. The limma model was then fitted using the patient sex as a covariate. Contrasts were then compared between each normal cell cluster and each sarcoma cluster separately (UPS, MFS, LMS, LPS, and SS). Genes with a Benjamini-Hochberg (BH) corrected p-value < 0.25 and log_2_(FC) ≠ 0 were considered as significantly differentially expressed. We used a more lenient BH-corrected p-value cutoff given the screening and exploratory nature of this part of our analysis.

#### Differential Distribution Testing with distinct

The R package distinct^38^ (version 1.10.2) was used to perform differential gene distribution testing between sarcoma cells and non-tumoral cells of the TME. Since distinct compares empirical cumulative distribution functions for a given gene across cell types, it allows identification of subtle differences in expression that are not captured by other methods which might consider only measures of central tendency in the expression distribution. We hypothesized that this could be particularly useful in comparing transcriptionally similar cells, as STS and non-tumoral mesenchymal cells in TME. Due to its computational load, it resulted unfeasible to use distinct for the HPA-TS-HCA normal cells comparisons. The raw gene expression matrices of sarcoma cells and non-tumoral cells were first processed in Seurat as described above to create a merged Seurat object. Afterwards, a single cell experiment object specific to each tumor versus normal comparison was created using the SingleCellExperiment R package^130^ (version 1.20.1). The expression data was then log normalized using the logNormCounts function from the R package Scuttle^131^ (version 1.8.4). A “batch” variable indicating the sex of the patient was included in the design matrix as a covariate, unless patients were of the same sex. The distinct_test function was then used to run differential distribution testing using default parameters. To obtain the log_2_ FC of the gene expression between sarcoma and normal cells, counts were first converted to CPM using the calculateCPM function from Scuttle^131^, and these were used as input for the log2_FC function in distinct^38^. Then the log_2_FC was then calculated as log_2_ 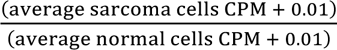, where the average RNA expression in the normal and sarcoma cells were calculated through the distinct function log2_FC. Genes with a log_2_FC ≠ 0 and a “p_adj.glb” value < 0.25 were considered as significantly differentially expressed between sarcoma and normal cells.

#### Identification of differentially expressed genes in STS

For each STS histotype, we defined a gene as up-regulated, if it resulted as differentially up-regulated (log_2_FC > 0, FDR < 0.25) in sarcoma cells in at least one of the comparisons made, either against reference cells in the TME (limma-voom or distinct), or in the HPA-TS-HCA datasets (limma-voom). STS down-regulated genes were instead defined as those that resulted as down-regulated in sarcoma cells (log_2_FC < 0, FDR < 0.25) in at least one of the comparisons made, and that were never up-regulated in sarcoma cells, separately considering the TME and the HPA-TS-HCA tests. For genes with more than one alternative transcript, these were tested separately.

### Hierarchical Clustering of STS versus non-tumoral cells in the TME and normal tissues

To perform hierarchical clustering between STS and non-tumoral cells in the TME as well as normal cells in the HPA-TS-HCA datasets, we used the R package factoextra (version 1.0.7) and base R functions. First, we filtered the CPM normalized scRNA count matrices to retain only genes defined as differentially expressed in STS (either up-or down-regulated), considering the TME and HPA-TS-HCA comparisons separately. Then we merged the count matrices from different samples/tissues, averaged gene expression per cell type and log_2_(+1) transformed the expression values. Pearson correlation was then used to obtain pairwise distances between cell types, separately analyzing STS cells versus non-tumoral cells in the TME, and STS cells versus normal cells in HPA-TS-HCA (supplementary data). The Pearson distance matrices were used to create a dendrogram with the hclust base R function, setting the agglomeration method to “ward.D2”, and this was then visualized using the fviz_dend function from factoextra. The optimal number of clusters, *k*, was obtained by calculating the silhouette score to estimate how similar cells within the same cluster are to each other and how different they are from cells in other clusters, for *k* ∈ {2,…, *n* − 1}, where *n* is the number of cells considered. The silhouette score peaked at *k* = 3 for the TME comparisons, and around *k* = 5 and *k* = 6, for the HPA-TS-HCA comparisons. We chose *k* = 6 for the latter as it provided a more biologically relevant separation of the different cell types.

### List of Surface Proteins

To obtain a comprehensive list of genes coding for proteins that are known to or predicted to be expressed on the plasma membrane, we combined The Cancer Surfaceome Atlas^14^ with the in silico human surfaceome (SURFY)^132^ after conversion of gene names to standard HGNC symbols (**Table S4**).

### Normal Tissue Expression Datasets

To assess the suitability of a given gene as a cellular therapy target, we used several normal tissue and cell datasets to measure RNA and protein expression.

Normal tissue RNA expression was assessed using the bulk RNA-seq dataset from the Human Protein Atlas^39^ (version 22.0, Ensembl version 103.38) accessed at https://www.proteinatlas.org/about/download (file name: rna_tissue_hpa.tsv.zip), and the bulk RNA-seq dataset from the GTEx^66^ accessed at https://gtexportal.org/home/datasets (expression file: GTEx_Analysis_2017-06-05_v8_RNASeQCv1.1.9_gene_tpm.gct; annotation file: GTEx_Analysis_v8_Annotations_SampleAttributesDS.txt).

RNA expression in normal cells was evaluated through the HPA single cell RNA-seq dataset (version 22.0, Ensembl version 103.38), accessed at https://www.proteinatlas.org/about/download (file name: rna_single_cell_type.tsv.zip).

Protein expression in normal tissues was evaluated using three separate datasets:

1. The HPA dataset (version 22.0, Ensembl version 103.38) obtained at (file name: normal_tissue.tsv.zip), which contains protein expression data across 44 normal human tissues obtained through immunohistochemistry.
2. The Human Proteome Map^67^ (HPM) dataset obtained at http://www.humanproteomemap.org/download.php, which reports protein expression data in 30 normal human tissues obtained through LC-MS/MS proteomics.
3. The GTEx Proteomics dataset^133^ which reports protein expression data in 32 normal human tissues, obtained through tandem mass tag (TMT) proteomics. Data was obtained from the Supplemental data (Table S2D) of the corresponding manuscript. Specifically, we used the log_2_ transformed protein abundances relative to the average of the reference in the same run.

#### Dataset pre-processing

Gene names across datasets were standardized using the R package mygene^134^ (version 1.38.0) by converting the gene name aliases to the corresponding standardized HGNC symbol. For the bulk RNA-seq datasets (GTEx and HPA), TPM expression values were converted to log_2_(TPM +1). Then, the expression data was processed differently based on the target discovery pipeline considered. For the permissive on-target off-tumor toxicity filter of the monospecific CAR target discovery pipeline, RNA expression from the GTEx dataset was first averaged across all samples belonging to a given tissue type. Then tissues were organized in tissue groups in a similar fashion as done by MacKay et al.^45^, and the average gene expression per tissue group was calculated (**Tables S5**, **S6**). Similarly, for the HPA dataset, the RNA expression was averaged across tissues belonging to same tissue group. Then, tissue expression in both the GTEx and HPA dataset was categorized as “Medium–High” (M-H) for log_2_(TPM+1) ≥ 4, and “Not Detected–Low” (ND-L) for log_2_(TPM+1) < 4. In all the other pipelines, the expression matrices of the bulk RNA-seq GTEx and HPA dataset were median-centered across tissues (HPA) or samples (GTEx), and then z-scored within genes, before use for downstream analysis.

In the HPA IHC protein expression dataset, the protein expression is reported for each cell type in each tissue as either “Not Detected”, “Low”, “Medium”, or “High”. For use in the permissive on-target off-tumor toxicity filter of the monospecific CAR target discovery pipeline, protein expression for each tissue was classified as follows: “ND-L” if all the cells in a tissue had only “Not Detected” or “Low” expression; “M-H” if all the cells in a tissue had “Medium” or “High” expression, or only “High” expression; “Medium” if all the cells in a tissue had “Medium” expression; “Variable” if the cells in a tissue had either “Not Detected”, “Low”, “Medium” or “High” expression. Tissue group expression was then calculated as the highest tissue expression within a tissue group using the following hierarchy: M-H > Variable > Medium > ND-L. Again, tissue groups were defined as above. For use in all the other pipelines, we first converted the cell type protein expression level from a categorical to a numerical score, where “Not Detected” = 1, “Low” = 2, “Medium” = 3, and “High” = 4. Then, we calculated the average expression value per tissue group for each gene, and then obtained a gene per tissue group expression matrix that was median-centered across tissue groups and z-scored within genes for downstream analysis.

In the HPA single cell RNA-seq dataset, gene expression is reported as normalized TPM (nTPM) for each normal cell cluster in 31 tissues. We first, converted the nTPM values to log_2_(nTPM+1) and then obtained a gene per cell cluster matrix, where columns represented cell clusters in each normal tissue in the dataset. The log_2_(nTPM+1) expression values were then median-centered across cell clusters and z-scored within genes, and this matrix was used for downstream analysis.

The LC-MS/MS proteomics expression data from the HPM was log_2_(+1) transformed and then median-centered across tissues and z-scored within genes, before further analysis. Lastly, in the GTEx TMT proteomics dataset the relative log_2_ transformed protein abundance data provided by the authors was median-centered across tissue samples and z-scored within genes, similarly to the other datasets. All the processed normal tissue and cell expression datasets used in the study are provided in the supplementary data.

### Sarcoma Cell Lines RNA Expression

The HPA cancer cell line expression was downloaded from HPA (version 22.0, Ensembl version 103.38) at https://www.proteinatlas.org/about/download (file: rna_celline_cancer.tsv.zip). The dataset was subsetted to retain only data corresponding to sarcoma cell lines including: GCT, HT-1080, MES-SA, RD, Rh18, Rh30, Rh41, RKN, S-117, SK-LMS-1, SK-UT-1, TE 441.T, TE 617.T, and U-2197. For each gene, HPA reports the average expression as nTPM for all the cell lines of a given cancer type. The nTPM values were averaged for those genes with alternative transcripts, and then converted to log_2_(nTPM+1) before use for downstream analyses. The processed data is provided in the supplementary data.

### Monospecific CAR-T Cell Discovery Pipeline

To define genes suitable for a monospecific CAR-T cell design, first we removed from our initial pool of STS up-regulated genes any genes that were up-regulated more often in normal cells than in sarcoma cells, considering all the comparisons in which that gene appeared. Then this gene pool was filtered to retain only those present in the surfaceome. HLA class I and II genes were also removed from the pool.

To increase tumor-specificity of our candidate targets, we filtered the gene pool to retain only those with high tumor expression in both our scRNA-seq data, in TCGA SARC, and in the HPA sarcoma cell lines (bulk RNA-seq). Tumor expression in our scRNA-seq data was calculated as the average log_2_(CPM+1) across the sarcoma cells of a given STS histotype, where CPM indicates RNA expression in counts per million, while RNA expression in TCGA SARC was calculated as the average log_2_(TPM+1) across all the samples of a given STS histotype. Then, we filtered our candidate pool to retain any gene that had above-median expression in both the SARC TCGA data and in the scRNA-seq data, or above-median expression in the scRNA-seq data and in the HPA sarcoma cell lines. Genes without expression data in SARC TCGA or sarcoma HPA cell lines but with scRNA-seq expression ≥ the 50^th^ percentile were also retained.

Then we assessed the potential for on-target off-tumor toxicity of each candidate target by measuring its RNA and protein expression in normal tissues and cells and comparing it to that of targets for which CAR-T cells are currently available for clinical use or in clinical development. The list of tested CAR-T targets was obtained from the previous work from MacKay et al^45^ (**Table S7**). Specifically, we identified genes at risk for on-target off-tumor toxicity by applying both a *permissive* and a *stringent filter*.

For our *permissive filter*, we used the HPA IHC protein database, as well as the GTEx and HPA bulk RNA-seq databases. Specifically, protein and RNA expression were categorized as described above, and then the number of tissue groups (TGs) with medium-high (M-H) expression was measured for the tested CAR-T targets. We used the 90^th^ percentile of the number of TGs with M-H expression for the tested CAR-T targets in each of the three datasets as cutoffs. A gene was considered to have passed the filter in a dataset if it had number of TGs with M-H expression below the respective cutoff.

For our *stringent filter*, we assessed the risk of on-target off-tumor toxicity both at the single cell level, using the HPA scRNA-seq dataset, and by measuring protein expression at the tissue level, using the HPA, HPM, and the GTEx protein expression datasets. For this analysis, we used the HPA protein numerical scoring classification obtained as described above. Also, for the HPA scRNA-seq dataset we used the average cell type expression across all tissues. Again, we used the normal cell/tissue expression of tested CAR-T targets as a reference to score our candidate genes against.

For a gene to have an acceptable on-target off-tumor toxicity risk based on its expression in the HPA scRNA-seq dataset, it had to meet two conditions. First, we categorized gene expression as “high” for z-score> 1, and “low” for z-score ≤ 1. Based on this, genes from our pool that had “high” expression in ≤14 cell types, which corresponds to the 75^th^ percentile of the number of cell types with “high” expression for the tested CAR-T targets, were initially retained. Then, we calculated the maximum expression value per cell type for the tested CAR-T targets and retained only those genes that had RNA expression below this threshold in each cell type. The following cell types were not considered: lymphoid immune cells (T-cells, NK-cells, B-cells, plasma cells), due to the potentially higher tolerability of their depletion; fibroblasts, adipocytes, and endometrial stromal cells due to potential high transcriptional similarity to the STS in the study; and cells in the gonads and placenta (ovarian stromal cells, granulosa cells, oocytes, peritubular cells (testis), cytotrophoblasts, extravillous trophoblasts, Hofbauer cells, syncytiotrophoblasts, Leydig cells, spermatocytes, late spermatids, spermatogonia, early spermatids, and Sertoli cells) being sites of immune privilege. Only those genes that met both criteria were retained based on their single cell RNA expression in normal cells.

For each of the 3 protein expression datasets, we calculated the maximum expression values per tissue (GTEx and HPM) or tissue group (HPA) for the tested CAR-T targets and used these as thresholds so that only the candidate targets with protein expression below the threshold in each of the tissues or tissue groups studied were retained. The following tissues and tissue groups were not considered: fetal tissues, male or female reproductive tissues, placenta, CD4 T-cells, CD8 T-cells, NK-cells, B-cells, spleen, immune cells and immune tissue. Notably, for the GTEx proteomics dataset, we used the average z-scored protein expression across all samples belonging to a given tissue.

For each candidate gene (*g*), we then combined the results of both filters by calculating a monospecific CAR Targetability Score, *TS_CAR_*, as follows:

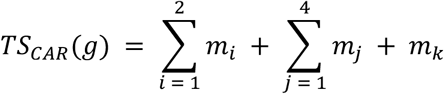

Where:

- *m*_*i*_ represents the score from the 2 bulk RNA-seq datasets (HPA and GTEx categorical) and is calculated as +1 if the gene passes the filter, –1 if it does not, and 0 if the gene is not present in the dataset.
- *m*_j_ represents the score from any of the 4 protein expression datasets (HPA categorical, GTEx proteomics, HPM, and HPA numeric) and is calculated as +2 if the gene passes the filter, –2 if it does not, and 0 if it is not present in the dataset.
- *m*_*k*_ represents the score from the HPA scRNA-seq dataset and is calculated as +1.5 if the gene passes the filter, –1.5 if it does not, and 0 if it is not present in the dataset.

Any gene with TS*_CAR_* ≥ 5 which corresponds to the 25^th^ percentile of the *TS_CAR_* for the tested CAR-T targets was considered as a suitable target for a monospecific CAR-T cell. The results of the pipeline are reported in **Table S8**.

### Gene set over-representation analyses

Gene set over-representation analysis (ORA) was performed using the R packages, clusterProfiler^135^ (version 4.10.0), ReactomePA^136,137^ (version 1.46.0), and org.Hs.eg.db^138^ (version 3.18.0). Before analysis, the gene symbols were converted to Entrez gene IDs. Then, we analyzed all the candidate targets for monospecific CAR-T cells for each STS, using the function enrichGO for Gene Ontology (GO) term enrichment, the function enrichKEGG for Kyoto Encyclopedia of Genes and Genomes (KEGG) pathways enrichment, the function enrichPathway for enrichment of Reactome pathways, and the function enrichWP for enrichment of WikiPathways terms. The p-value cutoff for significance was set to 0.05 and the Bonferroni-Hochberg method was used for FDR correction, while minimum and maximum gene sets sizes were set respectively to 10 and 500. We then performed a clustered ORA on the monospecific CAR-T targets by STS histotype. In this case we used the compareCluster function from clusterProfiler and within this we used the same enrichment functions as described above, depending on the reference database used, again setting FDR cutoff to 0.05 and minimum and maximum gene sets sizes to 10 and 500. When performing the cluster ORA analysis on the TCR-T and PC-CAR targets, the FDR cutoff was instead set to 0.1 due to smaller gene pool to test, while the minimum and maximum gene sets sizes were set to 2 and 500, respectively.

### TCGA Pan-Cancer Atlas

RNA expression data for the Pan-Cancer Atlas TCGA^139^ samples (batch normalized RSEM values) was obtained from cBioPortal^140,141^ for the genes of interest. The downloaded data is available in the Source Data. Data for the following cancer types was analyzed: KICH: Kidney Chromophobe; ACC: Adrenocortical carcinoma; LIHC: Liver hepatocellular carcinoma; UVM: Uveal Melanoma; PCPG: Pheochromocytoma and Paraganglioma; KIRP: Kidney renal papillary cell carcinoma; LAML: Acute Myeloid Leukemia; THYM: Thymoma; SKCM: Skin Cutaneous Melanoma; KIRC: Kidney renal clear cell carcinoma; READ: Rectum Adenocarcinoma; LGG: Brain Lower Grade Glioma; CHOL: Cholangiocarcinoma; COAD: Colon Adenocarcinoma; PRAD: Prostate Adenocarcinoma; STAD: Stomach Adenocarcinoma; UCEC: Uterine Corpus Endometrial Carcinoma; CESC: Cervical squamous cell carcinoma and endocervical adenocarcinoma; BLCA: Bladder Urothelial Carcinoma; ESCA: Esophageal Carcinoma; DLBC: Diffuse Large B-Cell Lymphoma; TGCT: Testicular Germ Cell Tumors; OV: Ovarian serous cystadenocarcinoma; LUSC: Lung Squamous Cell Carcinoma; LUAD: Lung Adenocarcinoma; HNSC: Head and Neck Squamous Cell Carcinoma; BRCA: Breast Invasive Carcinoma; PAAD: Pancreatic Adenocarcinoma; GBM: Glioblastoma Multiforme; THCA: Thyroid Carcinoma; UCS: Uterine Carcinosarcoma; SARC: Sarcoma; MESO: Mesothelioma.

### Cell Line Gene Dependency

The DepMap project^57,71^ reports the results of CRISPR KO experiments on 1100 human cell lines. To assess the essentiality of our candidate targets for sarcoma cell lines (including both soft tissue sarcomas as well as chondrosarcoma, osteosarcoma and Ewing’s sarcoma), we used the “CRISPRGeneEffect.csv” file obtained from the DepMap website, containing data regarding the effect of the KO of each gene in each cell line. Gene effects less than –1 indicate a significant reduction in cell growth and proliferation, as this was the median gene effect estimate of genes known to be essential for cell growth and survival.

### Soft Tissue Sarcoma Genomics Data

Data on copy number alterations (CNAs) and mutations in soft tissue sarcomas were obtained from the TCGA SARC firehose database^36^ and from the MSK-IMPACT^58^ sarcoma study. Both datasets were downloaded from cBioPortal^140,141^. The file named “data_linear_cna.txt” was used to analyze linear CNA data from TCGA firehose, while the file named “data_cna.txt” was used to determine amplification status of a given gene in each sample for both the TCGA and MSK-IMPACT databases. For the Pearson correlation analysis between linear CNA and RNA expression, the TCGA SARC bulk RNA-seq expression data obtained as above, was used. Correction for multiple hypothesis was obtained using the BH method setting the threshold for significance to 0.01.

Data regarding single nucleotide variants (SNVs) and small insertions/deletions (Indels) was retrieved from the “data_mutations.txt” files for both the TCGA and the MSK-IMPACT cohorts. Lastly, data on gene fusions in soft tissue sarcomas was obtained by integrating data in the “data_sv.txt” file obtained from cBioPortal for the MSK-IMPACT cohort, data from the study from Gao et al.^142^ on recurrent gene fusions in TCGA tumor samples, and from a search through the Mitelman Database^61,143^ (https://mitelmandatabase.isb-cgc.org/mb_search) that included the following sarcoma types: Ewing sarcoma; Atypical lipomatous tumor/atypical lipoma/well-differentiated liposarcoma; Alveolar rhabdomyosarcoma; Myxoinflammatory fibroblastic sarcoma; Undifferentiated pleomorphic sarcoma; Liposarcoma, dedifferentiated; Inflammatory myofibroblastic tumor/myofibroblastic sarcoma; Soft tissue tumor, NOS; Clear cell sarcoma; Undifferentiated round cell sarcoma; Undifferentiated sarcoma; Mesenchymal tumor, NOS; Fibrosarcoma; Embryonal rhabdomyosarcoma; Spindle cell/sclerosing rhabdomyosarcoma; Myxofibrosarcoma; Endometrial stromal sarcoma; Pleomorphic rhabdomyosarcoma; Liposarcoma, NOS; Alveolar soft part sarcoma; Leiomyosarcoma; Sclerosing epithelioid fibrosarcoma; Malignant soft tissue tumor, NOS; Angiosarcoma; Rhabdomyosarcoma, NOS; Epithelioid sarcoma; Dermatofibrosarcoma protuberans/Bednar tumor/giant cell fibroblastoma; Low-grade fibromyxoid sarcoma; Synovial sarcoma; Histiocytic sarcoma/malignant histiocytosis; Liposarcoma, myxoid/round cell; Liposarcoma, pleomorphic. The combined fusion data in STS is available from **Table S15**.

### Logic-gated ‘AND’ CAR Targets Discovery Pipeline

To discover gene pairs suitable for a bispecific logic-gated ‘AND’ CAR design, first we selected genes up-regulated in sarcoma cells, separately for each STS, as described above. Next, we removed from this pool of STS up-regulated genes any genes that were up-regulated more often in normal cells than in sarcoma cells, considering all the comparisons in which that gene appeared. Then, we filtered this gene pool to include only highly expressed genes, as defined in the monospecific CAR-T cell target discovery pipeline. Then, for the selected genes we computed all possible pairs and measured gene co-expression at the single cell level by running the CS-CORE algorithm from the R package CSCORE^65^ (version 0.0.0.9). Specifically, we ran CS-CORE on all the tumor cells of a given sarcoma histotype and corrected p-values for multiple hypothesis testing using the BH method. Any non-significant correlations were converted to a score of 0, using an adjusted p-value significance threshold of 0.05. For a given gene pair (*g_i_, gj*), CS-CORE computes a correlation score, which we will refer to as *ρ_Tumor_*(*g_i_, gj*), and which can assume values between –1 (lowest likelihood of co-expression) and +1 (highest likelihood of co-expression).

Then, for each gene pair we assessed co-expression in normal cells and tissues, with the caveat that genes for an ideal target pair of an ‘AND’ logic-gated CAR would have mutually exclusive expression across all normal cells and tissues. To assess this, we used, the GTEx and HPA bulk RNA-seq databases, the HPA scRNA-seq database, the HPA IHC protein expression database, as well as the TMT GTEx proteomics and the HPM LC-MS/MS proteomics datasets. For the bulk RNA-seq datasets (GTEx and HPA), the median-centered, z-scored numerical gene expression matrices were used, where expression per each tissue (HPA), or tissue sample (GTEx) was measured. In this case, we used Pearson correlation to assess gene co-expression across normal tissues. Correction for multiple hypothesis testing was performed using the BH method, and for pairs with a corrected p-value ≥ 0.05 the correlation coefficient *ρ* was set to 0. Similarly, for the scRNA-seq HPA database, we used Pearson correlation to measure gene co-expression across normal cell clusters, with non-significant correlations set to *ρ* = 0, using a BH-corrected p-value threshold of 0.05. For the GTEx TMT proteomics database, we first removed any genes with more than 25% missing values across all tissue samples analyzed. Then, we applied the Pearson correlation as above to assess protein co-expression across normal tissue samples, again using a BH-corrected p-value threshold of 0.05 for significance and setting *ρ* = 0 for any non-significant correlations.

For the HPA IHC database (numerical score data) and for the HPM proteomics database, given the low number of comparisons, we did not use Pearson correlation but instead calculated a modified odds ratio (OR) for each gene pair (*g_i_, gj*) as follows:

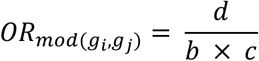

Where:

- *d* is the number of tissues/cells where both *g_i_* and *g_i_* have z-score > 1
- *b* is the number of tissues/cells where *g_i_* has z-score > 1 and *g_j_* has z-score ≤ 1
- *c* is the number of tissues/cells where *g_j_* has z-score > 1 and *g_i_* has z-score ≤ 1

In this case, pairs with mutually exclusive protein expression were considered those where *OR*_*mod*_ = 0. Of note, if both *d* and *b* × *c* were 0, the OR was set to 0.

Then, for each gene pair (*g*_*i*_, *g*_j_) we combined results from each normal tissue/cell database to obtain a score that estimates gene co-expression across normal tissues and cells, *N*b*g*_*i*_, *g*_j_c, as follows:

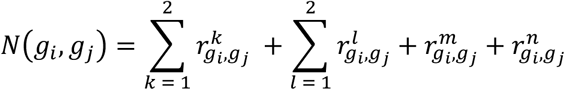

Where:

- 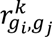 represents the score from the 2 bulk RNA-seq datasets and is calculated as:

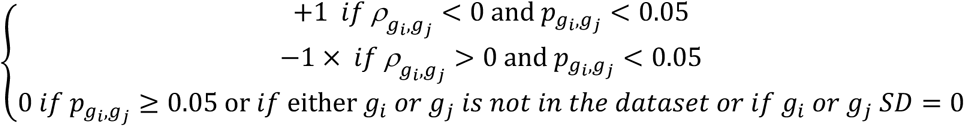
- 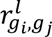 represents the score from the HPA IHC and the HPM proteomics datasets and is calculated as:

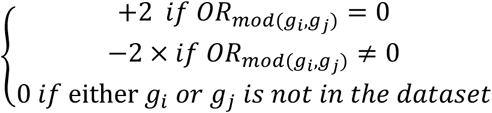
- 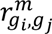 represents the score from the HPA scRNA-seq dataset and is calculated as:

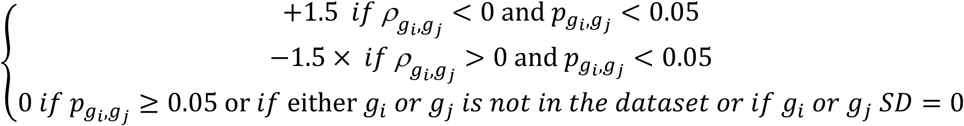
- 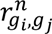 represents the score from the GTEx proteomics dataset and is calculated as:

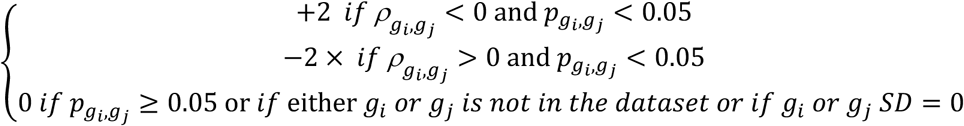
- 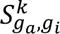 is the Pearson correlation coefficient in a dataset between the two genes in a pair and *p_g*i*,gj_* is the respective BH-corrected p-value.

The raw score *N*(*g*_*i*_, *g*_j_) was then normalized and scaled as follows:

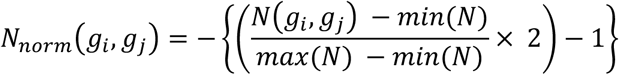

Where:

- *N*(*g*_*i*_, *g*_j_) is the raw score for a given gene pair (*g*_*i*_, *g*_j_)
- min(*N*) and max(*N*) are the maximum and minimum raw scores across all gene pairs for a given STS histotype, respectively.

This normalization ensures that a score between 0 and 1 indicates higher likelihood of gene co-expression across normal tissues/cells and conversely a score between −1 and 0 indicates a lower likelihood of co-expression.

Finally, for each gene pair (*g*_*i*_, *g*_j_) we calculated an ‘AND’ logic-gated Targetability Score, called *TS*_′*AND*′_ score, as follows:

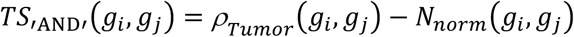

Gene pairs with *TS*_′AND′_ score ≥ 1 were considered to have higher fitness as targets for an ‘AND’-gated CAR design. The results of the pipeline by STS histotype are reported in **Table S17**.

### Logic-gated ‘AND–NOT’ CAR Targets Discovery Pipeline

To discover gene pairs for an ‘AND–NOT’ bispecific CAR design, we applied a similar logic to that used for discovery of gene pairs for an ‘AND’-gated CAR. In this case, we looked for genes with high expression in sarcoma cells that would function as activators, and for genes with low to no expression in sarcoma cells but high co-expression with the paired activator in normal cells/tissues that would function as inhibitors, thus preventing on-target off-tumor toxicity.

To do this, first we obtained a list of highly expressed sarcoma up-regulated genes for each STS histotype, removing any genes up-regulated in normal cells more often than STS cells, as described above, and considered these as candidate activators. Then, to define a list of possible inhibitors with little to no expression in sarcoma cells, for each STS histotype we selected surfaceome genes with scRNA-seq expression below the 50^th^ percentile among the sarcoma down-regulated genes as defined in our differential gene expression pipeline. Single cell RNA expression was again measured as the average log_2_(CPM+1) across all the tumor cells in a sarcoma histotype.

We then calculated all the possible unique pairs between the candidate activators and inhibitors and measured their co-expression in normal cells/tissues using the same datasets used for the ‘AND’-gated CAR targets discovery pipeline. Specifically, for the GTEx and HPA bulk RNA-seq datasets, as well as for the scRNA-seq HPA and the GTEx TMT proteomics datasets we used Pearson correlation to measure co-expression of the two genes in each pair, and applied BH multiple hypothesis testing correction setting the threshold for significance to a p-value < 0.05. For non-significant correlations, we then set *ρ* = 0. For the GTEx TMT proteomics dataset, again we first removed any genes with more than 25% missing values across all tissue samples analyzed before performing correlation testing.

For the HPA IHC and the HPM proteomics databases, instead of performing Pearson correlation analysis we calculated a modified odds ratio (OR) for each gene pair (*g*_*a*_, *g*_*i*_) between an activator, *g*_*a*_, and an inhibitor, *g*_*i*_ as follows:

Where:

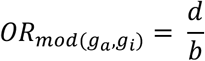

- *d* is the number of tissues where both the activator *g*_*a*_ and the inhibitor *g*_*i*_ have z-score > 1
- *b* is the number of tissues where the activator *g*_*a*_ has z-score > 1 and the inhibitor *g*_*i*_ has z-score ≤ 1

Pairs where *OR*_*mod*_ = *Inf* were considered as suitable for an ‘AND–NOT’ CAR design. In cases where both *d* and *b* were 0, we set *OR*_*mod*_ to *Inf*.

Then, for each gene pair (*g*_*a*_, *g*_*i*_) we combined results from each normal tissue/cell database to calculate an ‘AND–NOT’ logic-gated Targetability Score, *TS*_′*AND*–*N*0*T*′_ as follows:

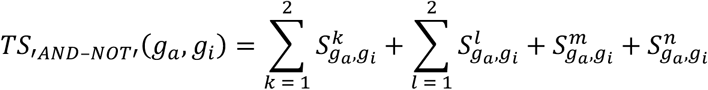

Where:

- 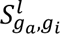 represents the score from the 2 bulk RNA-seq datasets and is calculated as:

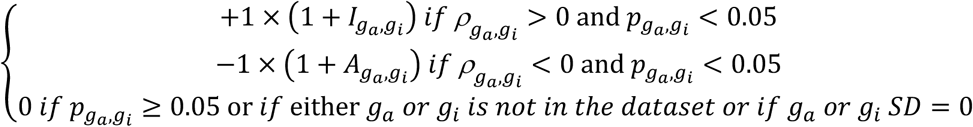
- 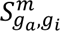 represents the score from the HPA IHC and the HPM proteomics datasets and is calculated as:

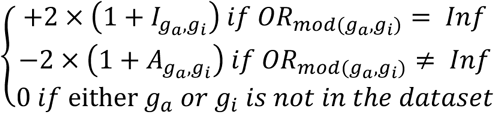
- 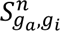 represents the score from the HPA scRNA-seq dataset and is calculated as:

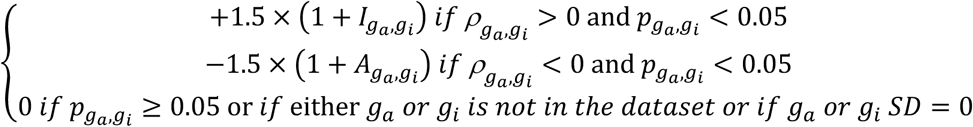
- *S*^*n*^ represents the score from the GTEx proteomics dataset and is calculated as:

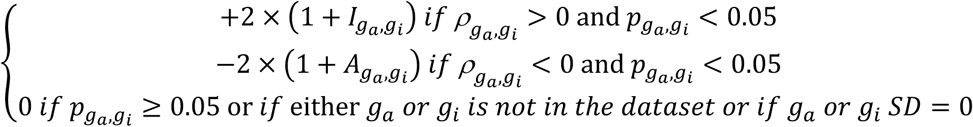
- *I*_g*a*,g*i*_ is the proportion of comparisons in a dataset where the expression of the inhibitor is equal or above that of the activator (*z* − *score*_*gi*_ ≥ *z* − *score*_*ga*_)
- *A*_g*a*,g*i*_ is the proportion of comparisons in a dataset where the expression of the activator is above that of the activator (*z* − *score*_g*a*_ > *z* − *score*_g*i*_)
- *ρ*_g*a*,g*i*_ is the Pearson correlation coefficient in a dataset between the activator and the inhibitor and *p*_g*a*,g*i*_ is the respective BH-corrected p-value.

The *TS*_‘AND–NOT’_ scores were then normalized as follows:

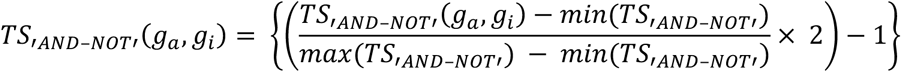

Where:

- *TS*_′*AND*–*N*0*T*′_(*g*_*a*_, *g*_*i*_) is the raw score for a given gene pair (*g*_*a*_, *g*_*i*_)
- *min*(*TS*_′*AND*–*N*0*T*′_) and *max*(*TS*_′*AND*–*N*0*T*′_) are the maximum and minimum raw scores across all gene pairs for a given STS histotype, respectively.

In each sarcoma histotype, gene pairs with values above the 99^th^ percentile of the normalized *TS*_′*AND*–*N*0*T*′_ score were considered as those suitable for an ‘AND-NOT’-gated CAR design. The results of the pipeline by STS histotype are reported in **Table S18**.

### HPA Single Cell Gene Clusters and Signatures

HPA categorizes genes based on their expression profile across different normal cell types in their scRNA-seq dataset^144^. Specifically, genes are categorized into 80 clusters, with one or more clusters corresponding to a specific cell type. We downloaded the gene cluster assignments directly from the HPA website and grouped gene clusters specific to a given cell type together to obtain cell type-specific gene signatures that were used to annotate genes in each bispecific CAR target pair. Also, the expression of these gene signatures in our STS scRNA-seq data was measured using the Seurat function AddModuleScore and shown in **Figure S7**. The HPA single cell gene clusters and signatures are provided in the Supplementary Data.

### T cell receptor-engineered T cell and Peptide-Centric CAR Target Discovery Pipeline

To identify targets for TCR-T or PC-CAR-T cells in STS, we obtained a list of up-regulated genes for each of the STS histotype studied. This was defined as any gene, either coding for intracellular or surface proteins, that was up-regulated in at least one of the DET comparisons made. Within this gene pool we then identified genes with high STS expression, defined in this case as those with expression ≥ 25^th^ percentile in both our scRNA-seq data and in the SARC TCGA samples, or those genes with expression ≥ 25^th^ percentile in both the sarcoma HPA cell lines and our scRNA-seq data. Genes without expression data in SARC TCGA or sarcoma HPA cell lines but with scRNA-seq expression ≥ the 25^th^ percentile were also considered to have high STS expression.

Then we compared the expression in normal cells and tissues of each of the up-regulated genes to that of known TCR-T cell therapy targets that are currently in clinical development identified through a search on clinicaltrials.gov with the term “solid tumor” for the Condition/disease field, and “TCR-T cells” or “cellular therapy” for the Intervention/treatment field (**Table S22**). Specifically, only genes belonging to the cancer germline antigen (CGA) class were chosen as reference due to their preferential expression in immune-privileged sites like the gonads.

Normal cell and tissue expression was measured again using the HPA scRNA-seq dataset (average cell type expression), the HPA and GTEx bulk RNA-seq datasets (average tissue group expression), the GTEx TMT proteomics dataset (average tissue expression), the HPM LC-MS/MS proteomics dataset, or the HPM IHC data, obtained as previously described.

For each gene (*g*) we calculated a summative score *N*(*g*), representing its normal cell/tissue RNA and protein expression as compared to that of the CGAs used as TCR-T cell targets, and calculated as follows:

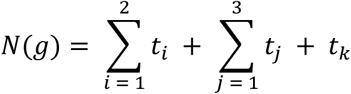

Where:

- *t*_*i*_ is the score calculated by comparing the candidate target expression to that of the reference targets in the HPA and GTEx bulk RNA-seq datasets and is calculated as +1 if the candidate has expression below the maximum expression value for the reference targets in each tissue group, 0 if the candidate has expression above the maximum value for the reference targets in at least one of the tissue groups or if the gene is not present in the parent dataset;
- *t*_j_ is the score calculated by comparing the candidate target expression to that of the reference targets in the HPA IHC, GTEx, and HPM proteomics datasets and is calculated as +2 if the candidate has expression below the maximum expression value for the reference targets in each tissue or tissue group, 0 if the candidate has expression above the maximum value for the reference targets in at least one of the tissues or tissue groups or if the gene is not present in the parent dataset;
- *t*_*k*_ is the score calculated by comparing the candidate target expression to that of the reference targets in the HPA scRNA-seq dataset and is calculated as +1.5 if the candidate has expression below the maximum expression value for the reference targets in each cell type and 0 if the candidate has expression above the maximum value for the reference targets in at least one of the cell types or if the gene is not present in the parent dataset.

Genes with a *N*(*g*) score ≥ 3.5 and with high STS expression as defined above were retained for further analysis. Due to the low number of genes with *N*(*g*) ≥ 3.5 meeting the combined high expression criteria, possibly related to RNA expression dilution in bulk tissue analysis (TCGA SARC) and differences between in vivo and in vitro analysis (HPA cell lines), genes that had *N*(*g*) score ≥ 3.5 and scRNA-seq expression ≥ the 25^th^ percentile, regardless of TCGA SARC or HPA cell line expression, were also considered for further analysis as below (**Table S23**).

#### Global HLA Allele Frequency Estimation

To estimate the global HLA allele frequencies at the HLA-B, HLA-C, and HLA-A loci, we downloaded allele frequencies for all the population studies available in the Allele Frequency Net Database (AFND)^77^. Then, we used the python package HLAfreq^145^ to combine allele frequencies across populations, first within countries and then across countries to obtain an average world allele frequency. Analyses were ran using default parameters. The population size of each country was use as weight in the global allele frequency calculation, as described by the developer. Population sizes by country updated to 2022 were downloaded from the World Bank (https://data.worldbank.org/indicator/SP.POP.TOTL?most_recent_year_desc=false), or from the United Nations Populations Division (Martinique, Turkey, Taiwan), or from the European Union Commission (Madeira, Azores).

#### Peptide-MHC targetability assessment

Next for each candidate target antigen, we used the MSiC model from HLAthena^74^ to infer presentation and binding to HLA class I molecules of all the possible 9-mer and 10-mer peptides derived from each target. The amino acid sequence of each candidate target in FASTA format was obtained from UniProt^146^. For our analysis, we chose the top 10 most frequent HLA class I alleles by geographical region worldwide as reported in the Allele Frequency Net Database (AFND)^77^ https://www.allelefrequencies.net/top10freqs.asp, and then applied DeepImmuno^75^ to measure the predicted immunogenicity, the ability to elicit a CD8^+^ immune response, of each peptide-MHC pair (*pMHC*). Then, for each HLA allele we calculated the average worldwide allele frequency across all the populations in which that allele was detected in the AFND (**Table S26**). Finally, for each *pMHC* of each candidate target gene *g*, a targetability score, *TS*_*TCR*/*PC*_, was calculated to assess its suitability as a target for either TCR-T or PC-CAR-T cells as follows:

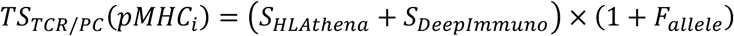

Where *pMHC*_*i*_ represents the *i^th^ pMHC* pair of a target gene *g, S*_*HLAt*ℎ*ena*_ is the score from the MSiC HLAthena model for *pMHC*_*i*_, *S*_*DeepImmuno*_ is the immunogenicity score assigned by DeepImmuno for *pMHC*_*i*_, and *F*_*allele*_ is the average worldwide frequency for the HLA allele in *pMHC*_*i*_. For each target antigen, we then ranked peptides based on their cumulative HLA allele frequency calculated as the sum of the allele frequencies of the HLA alleles in all of the *pMHCs* formed by each peptide, and then retained for cross-reactivity assessment only those *pMHCs* originating from peptides with the top five highest cumulative HLA allele frequency, and with *TS*_*TCR*/*PC*_ ≥ the 99^th^ percentile, and *S*_*HLAt*ℎ*ena*_ ≥ 0.85. The results for HLAthena and DeepImmuno for all the target antigens, as well as the *TS*_*TCR*/*PC*_ scores, are available in the supplementary data.

#### Peptide-MHC cross-reactivity assessment

For the *pMHCs* of each target antigen selected as described above, we assessed the likelihood of cross-reactivity with *pMHCs* derived from other proteins expressed in normal cells and tissues. To do this, we developed an algorithm called PS-TRACT (<u>P</u>eptide <u>S</u>imilarity and <u>T</u>CR <u>R</u>eactivity <u>A</u>nalysis for <u>C</u>ellular <u>T</u>herapies), that considers the similarity of each tumoral *pMHC* to a pool of normal *pMHCs* of reference while accounting for the likelihood of presentation of the cross-reactive *pMHCs* in normal tissues. Specifically, first we obtained a list of *pMHCs* experimentally detected in normal tissues/cells (*observed normal ligandome*) by integrating data from IEDB^147^, systeMHC^148^, and the HLA Ligand Atlas^149^. For IEDB, the full dataset was downloaded from https://iedb.org/database_export_v3.php (file: “mhc_ligand_full.csv.csv”), and epitopes were filtered to retain only linear peptides derived from human proteins and binding to MHC class I, with any qualitative binding measurement other than “Negative”, and of either 9 or 10 amino acids in length. Epitopes where MHC binding was measured with any of the following methods were also removed: half maximal effective concentration (EC50), dissociation constant KD (∼EC50), and dissociation constant KD (∼IC50). Lastly, only epitope binding data for alleles in standard four-digit typing HLA allele nomenclature were retained. For the HLA Ligand Atlas, the 2020.12 data release was obtained from https://hla-ligand-atlas.org/downloads. Epitope data was filtered to retain only 9-mer and 10-mer peptides binding to HLA class I or HLA class I and II, categorized as either strong or weak binders, and only those binding to HLA alleles reported in standard four-digit typing nomenclature. For systeMHC, data for the following SYSMHCIDs were obtained from https://systemhc.sjtu.edu.cn/datasets: SYSMHC00002, SYSMHC00003, SYSMHC00004, SYSMHC00007, SYSMHC00026, SYSMHC00028, SYSMHC00029, SYSMHC00031, SYSMHC00037, SYSMHC00039, SYSMHC00040, SYSMHC00044, SYSMHC00047, SYSMHC00052, SYSMHC00053, SYSMHC00055, SYSMHC00057, SYSMHC00064, SYSMHC00073, SYSMHC00088, and SYSMHC00089. In this case, only 9-mer and 10-mer peptides binding HLA class I and derived from human proteins were retained. To expand the pool of *pMHCs* presented in normal tissues, we ran MHCflurry^150^ for all the possible 9-mer and 10-mer peptides derived from the whole human proteome (*predicted normal ligandome*), using all the HLA alleles tested in our *pMHC* targetability pipeline as described above. Protein FASTA sequences for 20,409 proteins total were obtained from UniProt and mhcflurry-predict-scan was used to obtain binding, processing, and presentation predictions, with results filtered for affinity percentile < 2.0. Data from the observed and predicted normal ligandomes were combined to obtain a unified non-redundant reference dataset of unique *pMHC*/protein combinations (*unified normal ligandome*, supplementary data). Those *pMHCs* with MHCflurry presentation score ≤ 0.5 were removed from the unified normal ligandome unless they were listed in the observed ligandome. Then for each selected tumoral peptide we measured its similarity with normal peptides accounting for both their physicochemical and HLA binding properties. Specifically, given a tumoral peptide *p*_*t*_ and a normal reference peptide *p*_*n*_ both binding to an HLA allele *A*, peptide similarity *S*_*sim*_was calculated as:

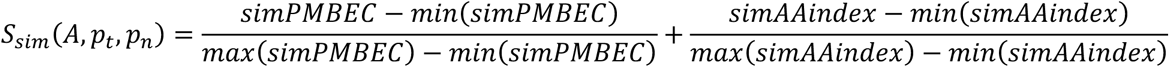

Where *simPMBEC* is a similarity score based on the peptide:MHC binding energy covariance (PMBEC) matrix^80^ which measures amino acid similarity based on their effect on peptide HLA binding. Specifically, *simPMBEC* was calculated as:

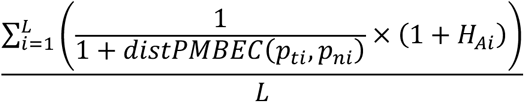

Where *distPMBEC* = *max*(*PMBEC*) − *PMBEC*, *PMBEC* is the value in the PMBEC matrix for the pair of amino acids at the *i*^*th*^ position of *p*_*t*_ and *p*_*n*_, *L* is the peptide length (9 or 10 amino acids), and *H*_*Ai*_ is the entropy at the *i*^*th*^ peptide residue calculated for all the peptides of same length as *p*_*t*_ binding to the HLA allele *A* in both the observed and predicted ligandome. This was obtained using the MolecularEntropy function from the R package HDMD^151^ (version 1.2), with values ranging from 0 to 1, indicating respectively no amino acid sequence variability and maximum sequence variability at the *i*^*th*^ peptide residue.

On the other end, *simAAindex* was calculated as follows:

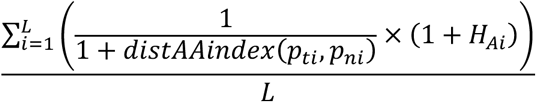

Where *distAAindex* is the Euclidean distance calculated between the pair of amino acids at the *i*^*th*^ position of *p*_*t*_ and *p*_*n*_, considering the first nine principal components of a PCA dimensionality reduction of 554 physicochemical amino acid properties available from AAindex^79^ and obtained through the R package ftrCOOL^152^ (version 2.0.0), *L* is the peptide length, and *H*_*Ai*_ is the residue entropy calculated as above. The results of the peptide similarity analysis for the candidate *pMHCs* for each target antigen are available in the supplementary data.

To define a significant threshold for *S*_*sim*_, we tested PS-TRACT on known cross-reactive peptide pairs^85^ and found that a similarity score of 0.95 was the lowest observed for cross-reactive peptides. Also, we ran PS-TRACT between the *MAGEA3*-derived peptide EVDPIGHLY and the *TTN*-derived peptide ESDPIVAQY, which was found to be the off-tumor target leading to cases of fatal myocarditis in patients that received EVDPIGHLY-targeted TCR-T cells^86^, obtaining a peptide similarity score of 1.19. Based on these results, we chose a threshold of *S*_*sim*_ > 1.15 as this provided the highest specificity in identifying known cross-reactive peptides (**Figure S11A**). Thus, after running PS-TRACT on each candidate tumoral *pMHC*, we selected only the corresponding normal *pMHCs* with *S*_*sim*_ > 1.15 for further cross-reactivity assessment.

Specifically, for each of these highly similar normal *pMHCs* we calculated a tissue-specific cross-reactivity score, *S*_*CR*_, that considers the tissue abundance of the antigen from which the cross-reactive peptide originates, the likelihood of presentation based on MHCflurry, and the presence of the peptide in the observed ligandome:

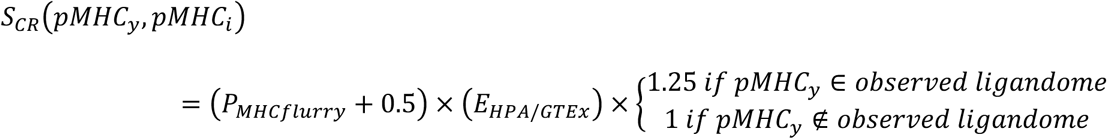

Where *pMHC*_*y*_ is the *y*^*th*^ cross-reactive *pMHC* for a given tumoral *pMHC*_*i*_, *P*_*MHCflurry*_ is the MHCflurry presentation score for *pMHC*_*y*_, and *E*_*HPA*/*GTEx*_ is the median-centered, z-scored RNA expression for the parent antigen of the *pMHC*_*y*_ peptide in the HPA and GTEx bulk RNA-seq consensus datasets (source: https://www.proteinatlas.org/about/download, file: rna_tissue_consensus.tsv.zip). Then for each tumoral *pMHC* in each of the 50 normal tissues, we selected the maximum *S*_*CR*_ score among all the filtered cross-reactive normal *pMHCs*, and this vector of *S*_*CR*_ scores was used to define the cross-reactivity profile of the tumoral target *pMHC* considered.

Finally, we benchmarked the cross-reactivity profile of each candidate tumoral *pMHC* pair against that of the *pMHC* targeted by afami-cel (GVYDGREHTV–HLA-A*02:01) which was used as reference given afami-cel’s promising safety and tolerability profile observed in clinical trials^27,28^. Specifically, for each candidate tumoral peptide-MHC pair *pMHC*_*i*_, we calculated the difference *S*_*CR*_*Diff*(*i*) between its *S*_*CR*_ cross-reactivity profile and that of GVYDGREHTV–HLA-A*02:01, as follows:

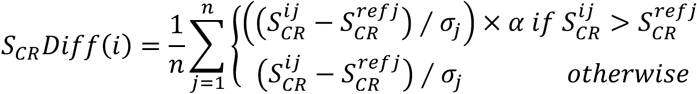

Where 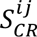 is the *S*_*CR*_ score in the *j*^*th*^ tissue of *pMHC*_*i*_, while 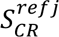 is the *S*_*CR*_ score in the *j*^*th*^ tissue of the reference *pMHC* (GVYDGREHTV–HLA-A*02:01), *n* is the number of tissues considered, *σ*_j_ is the standard deviation of the *S*_*CR*_ scores for all the *pMHC* pairs in the *j*^*th*^ tissue, and *α* = 2. Target tumoral *pMHCs* with differential *S*_*CR*_ profile compared to GVYDGREHTV–HLA-A*02:01 (*S*_*CR*_*Diff*) ≤ 0 were considered to have acceptable risk of cross-reactivity.

### Clinical STS proteomics datasets

Two clinical STS MS proteomics datasets were used to validate the expression of the identified cell therapy targets at the protein level, one from Tang et al.^91^ and one from Burns et al.^92^ The dataset from Tang et al. was obtained from the supplementary material of the relative paper (“Proteomic_matrix(tumor)” and “Clinical_information” tables). The relative fraction of total (Fot) protein intensities for the Tang et al. dataset were log_2_(+1) transformed and then median centered across samples and z-scored across all proteins for standardization. Protein symbols were standardized to the current HGNC symbols before analysis. The proteomics dataset from Burns et al. was obtained from the supplementary material (Supplementary Data 2). The expression matrix in this case was used as is given that protein expression values were already log-transformed, and median centered across samples and z-scored across proteins. Protein symbols were standardized to the current HGNC symbols before analysis. For SCAN-ACT implementation in other cancers, we obtained pre-processed mass spectrometry-based proteomics data from either LinkedOmicsKB^153^, the Proteomic Data Commons (PDC)^154^, or from a literature search (**Table S34**). Most of the data was derived from studies conducted by Clinical Proteomic Tumor Analysis Consortium (CPTAC) or the International Cancer Proteogenome Consortium (ICPC). For data obtained from LinkedOmicsKB, which was provided as log_2_ reference-normalized protein abundance, we applied median-centering across samples and z-scoring across proteins. In contrast, for data from the PDC, where the log_2_ reference-normalized protein abundance was already standardized across proteins, we performed only median-centering across samples. For the gastric adenocarcinoma dataset, expression data from different technical replicates of the same patient was averaged before analysis. For other cancers, expression data provided in the supplementary materials of the related publications was used^155–158^. For the prostate cancer dataset, which was provided as log2 reference-normalized protein abundance, we applied median-centering across samples and z-scoring across proteins. For the melanoma and esophageal cancer datasets, only median-centering across samples was necessary, as the data was already standardized across proteins. The bladder cancer expression data was used as provided by the authors, as it was already standardized across both samples and proteins. Prior to analysis, gene symbols across all datasets were updated to the current HGNC version. Proteins with missing values in 50% or more samples were removed, and the remaining missing values were imputed with a value of 1e-6. For genes with multiple protein isoforms, the isoform with the highest expression was used.

### STS cell line proteomics data

Protein expression data in soft tissue sarcoma cell lines was obtained from the Cancer Cell Line Encyclopedia^93^ (CCLE) and the ProCan-DepMapSanger repository. For CCLE, the normalized log_2_ transformed relative protein intensities were obtained from https://gygi.hms.harvard.edu/publicaShtions/ccle.html (file: protein_quant_current_normalized.csv). Gene symbols were standardized to the current HGNC symbols, and for different protein isoforms for a given gene symbol we retained the isoform with maximum expression (supplementary data). Then, data was median-centered across all cell lines and z-scored within genes before use for downstream analysis. The following “soft tissue” cell lines were considered in the CCLE dataset for further analysis: “A-204”, “RD”, “RH-41”, “HT-1080”, “RH-30”, and “KYM-1”. The ProCan-DepMapSanger dataset was obtained from the figshare repository associated with the corresponding publication: https://doi.org/10.6084/m9.figshare.19345397 (file: ProCan-DepMapSanger_protein_matrix_8498_averaged.txt). Similarly to the CCLE dataset, first we standardized gene symbols to the current HGNC nomenclature, then for genes with more than one protein isoform we retained the isoform with highest expression (supplementary data).

Since the data was already provided as log_2_ transformed relative protein intensities, the protein per cell line expression matrix was median-centered across cell lines and z-scored within genes before further analysis. The following soft tissue sarcoma cell lines were considered for downstream analysis: “A204”, “ESS-1”, “G-402”, “GCT”, “Hs-633T”, “HT-1080”, “KYM-1”, “MES-SA”, “MFH-ino”, “RD”, “RH-18”, “RH-41”, “RKN”, “SJRH30”, “SK-LMS-1”, “SK-UT-1”, “SKN”, “SW684”, “SW872”, “SW982”, “TE-441-T”, and “VA-ES-BJ”.

### IHC of Selected CAR-T Targets

IHC was performed by the Stanford Human Pathology/Histology Service Center for ACVR1, AXL, CLMP, GPC6, KITLG, and MRC2. Previously obtained tissue microarrays constructed using one to seven 1-mm FFPE cores per patient were used for the analysis, including UPS, LPS, LMS, SS, and MPNST samples. The following antibodies were used for the analysis: ACVR1 Rabbit IgG (cat. HPA007505, Sigma Aldrich, MO, USA), AXL Rabbitt IgG (cat. HPA037422, Sigma Aldrich, MO, USA), GPC6 (cat. HPA017671, Sigma Aldrich, MO, USA), KITLG (cat. HPA070395, HPA017671), CLMP (cat. 16127-1-AP, Thermo Fisher Scientific, MA, USA), and MRC2 (cat. HPA041991, Sigma Aldrich, MO, USA). All the antibodies were used at a dilution titer of 1:50 except for the CLMP antibody (1:400), and the KITLG antibody which was used at a 1:100, 1:200, or 1:50 dilution depending on the TMA analyzed. Tumor cell staining intensity was scored using a three-point scale (0, negative; 1, weakly positive; and 2, strongly positive). Staining for FAP was performed at the UCLA pathology department on a tissue microarray of 60 undifferentiated pleomorphic sarcomas in triplicate (no previous treatment), of which 57 samples were evaluable. FAP immunostaining was performed on Leica Bond platform using a polyclonal rabbit anti-FAP alpha antibody (Abcam, ab207178). All slides were scanned at 20X magnification. The percentage of FAP expressing cells was quantified using the HALO® image analysis platform based on automatic cell segmentation (Indica Labs). Missing or folded tissue cores were excluded. Replicate cores were averaged.

### Measurement of peptide-MHC binding through IMMUNE

#### Plasmid construction

To construct plasmids driving the expression of non-covalent HLA-I heterodimers, the commercially available backbone vector pRS315 was modified^89^. Briefly, the entire expression cassette containing genes encoding the Aga2 leader sequence, the ectodomain of a particular HLA-I heavy chain allele (A*02:01 or A*11:01 or B*35:01 or C*12:03), the Aga2p yeast native protein subunit, an HA epitope tag, and the MFα terminator was cloned to the downstream of the GAL10 promoter in pRS315, so that the expressed HLA-I heavy chain ectodomain was fused to the N-terminus of Aga2p for yeast surface display. Subsequently, a second expression cassette containing genes encoding the Aga2 leader sequence, β2m, a C-myc epitope tag and the MFα terminator was cloned to the downstream of the GAL1 promoter in the same pRS315-based vector, allowing the secretion of the β2m domain for HLA heavy chain/β2m dimerization and surface detection. In addition, the Aga2p subunit was further removed from the above plasmids, which resulted in control plasmids yielding no HLA detection on the surface of yeast. Gene synthesis and subcloning services were provided by GenScript (Jiangsu, China).

#### Electrotransformation of yeast cells and expression of HLA

The *Saccharomyces cerevisiae* yeast parent strain, EBY100^159^ was used for generation of daughter strains displaying each of the aforementioned HLA alleles. The constructed plasmids were transformed into EBY100 one at a time by electroporation, following the MicroPulser electroporation mannual (BioRad). Transformed cells that received the corresponding plasmid were recovered on selected dextrose-based (SD) medium agar plates minus the nutrient corresponding to the selection marker gene within the plasmid. Single colonies on SD agar plates appeared after 2-3 days of incubation at 30°C. To induce yeast expression of target proteins, a single yeast colony was inoculated with liquid SD medium and cultured in a shaking tube at 30°C until it reached the exponential growth phase. A sufficient amount of yeast cells was then transferred from the SD medium to selected galactose-based (SG) medium for protein expression under the promotion of GAL1 or GAL10. Typically, an overnight shaking and induction at 30°C can promote effective expression of the protein of interest in yeast for surface detection by flow cytometry.

#### Immunofluorescent labeling and flow cytometric analysis

A sufficient amount of galactose-induced yeast cells was collected by centrifugation and labeled by primary antibodies in PBS + 0.5% bovine serum albumin (BSA) for 1 h on ice. The primary antibody was rabbit anti-HA tag monoclonal antibody (mAb clone C29F4, Cell signaling), mouse anti-human β2m mAb (clone 2M2, Biolegend), or mouse anti-HLA-A, B, C conformation-specific mAb (clone W6/32, Biolegend). After the primary staining, the cells were washed at least twice with ice cold PBS + 0.5% BSA and labeled by secondary antibodies on ice for another 1 h. The secondary antibody was Alexa Fluor 488 conjugated goat anti-rabbit IgG (H+L) highly cross-adsorbed secondary antibody, Alexa Fluor 488 or Alexa Fluor 647 conjugated goat anti-mouse IgG (H+L) highly cross-adsorbed secondary antibody (Thermo Fisher Scientific). After the secondary staining, cells were washed with ice cold PBS + 0.5% BSA and resuspended in PBS + 0.5% BSA for flow cytometric analysis. A FACSCalibur flow cytometer (BD biosciences) was used to acquire flow data. At least 10000 cell events were collected for each sample and gated through forward and side scattering. FL1 and FL4 fluorescence channels were selected for Alexa Fluor 488 and Alexa Fluor 647, respectively, according to the fluorophore selected at the secondary labeling step as mentioned above. Flowjo (BD biosciences) was used for data analysis.

#### Peptide design and synthesis

To design indicator peptides appropriate for evaluation of HLA peptide binding on the surface of yeast cells and for downstream competitive assay development, the reported epitope repertoire for each aforementioned HLA allele from the Immune Epitope Database (https://www.IEDB.org) was carefully analyzed. First, an epitope peptide was considered only if it contained a cysteine residue at P4 or P5 or P6 anchor position available for site-specific modification, i.e., biotinylation or fluorophore conjugation. After that, the chosen peptide was further analyzed and validated 1) based on its predicted %Rank_EL (<2) and %Rank_BA scores (<2) from the NetMHCpan-4.1 webserver, and 2) based on its sequence similarity to the peptides with available X-ray structure in complex with the given HLA. For example, Bio-KLT (amino acids: KLTPL{Cys-Biotin}VTL), Bio-AIF (aa: AIFQ{Cys-Biotin}SMTK), Bio-RYP (aa: RYPV{Cys-Biotin}KFPSL), and Bio-FAF (aa: FAF{Cys-Biotin}RITSF) were designed as the indicator peptide for A*02:01, A*11:01, B*35:01, and C*12:03, respectively. All peptides were synthesized (GenScript) with over 95% purify.

#### On yeast peptide binding

About half a million yeast cells displaying a peptide-receptive HLA were collected by centrifugation, resuspended in 40 μl of PBS, and incubated with 0.5 uM biotinylated indicator peptides ([peptide]>>[HLA]) overnight prior to fluorescent labeling for flow cytometry. After peptide binding reached equilibrium, yeast cells were washed twice with ice-cold PBS + 0.5% BSA and then simultaneously stained for bound biotinylated indicator peptides and HLA molecules on the surface using Alexa Fluor 647 conjugated streptavidin and HLA conformation-specific mAb W6/32 followed by Alexa Fluor 488 conjugated goat anti-mouse IgG (H+L) highly cross-adsorbed secondary antibody (Thermo Fisher Scientific). Double stained cells were then washed and resuspended in ice-cold PBS + 0.5% BSA for analysis on a BD FACSCalibur. Flow cytometric data was analyzed using FlowJo.

#### On yeast peptide competition

To evaluate the competitive binding of a test peptide against the indicator peptide to HLA on the surface of yeast cells, about half a million yeast cells were collected and incubated with the biotinylated peptide as mentioned above, but with addition of a 20-fold more concentrated test peptide in its non-biotinylated format. The peptide competition assay was also performed under a similar condition as mentioned above, allowing reaction to reach equilibrium. Yeast cells were then collected, washed, and co-stained for bound biotinylated indicator peptides and HLA molecules using the aforementioned fluorescent labeling method for flow cytometric analysis. The %Binding in the presence of competitors was quantified as [(MFI_with competitor_−BG)/(MFI_no competitor_−BG)]×100%, where MFI_with competitor_ represents the average fluorescence intensity in the presence of test peptide (a competitor), MFI_no competitor_ represents the average fluorescence intensity in the absence of competitors, and BG represents the background fluorescence intensity. The %Competition was calculated as 100%−%Binding to reveal binding of the test peptide competitor to HLA molecules displayed on yeast. MFI was determined using appropriately gated cell populations.

### Target Priority Index

For each target discovery pipeline, we calculated a final Target Priority Index (TPI) that integrates normal tissues/cells target expression, scRNA-seq tumor expression and homogeneity, STS protein expression, DepMap sarcoma cell line essentiality, incidence of SNVs/Indels, gene fusions and CNAs in STS involving the target considered, and the presence of clinically or pre-clinically developed immunotherapeutic agents against each target (for monospecific and bispecific CAR-T cells targets only).

Homogeneity of expression at the single-cell level in each STS was calculated as the product of the average tumor expression in log_2_(CPM+1) and the percentage of tumor cells with log_2_(CPM+1) > 0, which we defined as *H*_*RNA*_ score:

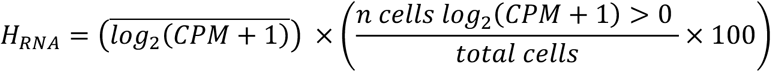

To assess if a gene has been previously studied as an immunotherapy target, we focused on therapeutic monoclonal antibodies (mAb), ADCs, single-chain variable fragments (scFv), fusion proteins, nanobodies, and fragment antigen binding regions (Fab), as well as CAR-T cells. Specifically, we downloaded a list of clinically developed mAbs, ADCs, scFvs, Fabs, nanobodies and fusion proteins from Thera-SAbDab^160^ (https://opig.stats.ox.ac.uk/webapps/sabdab-sabpred/therasabdab, accessed on 07/16/2024, Supplementary Data), retaining only agents developed for human use, and removing all those agents with clinical development phase reported as “TBC”, “Preregistration”, or “Preregistration (w)”. We complemented Thera-SAbDab with a PubMed search for pre-clinically developed therapeutic antibodies or ADCs against any genes in the surfaceome, using the following search query: “gene AND antibody, OR gene AND antibody drug conjugate, OR gene AND ADC OR gene AND antibody-drug conjugate” (**Table S30**). Targets with clinical development phase reported as “TBC”, “Preregistration”, or “Preregistration (w)” in Thera-SAbDab were included as pre-clinical targets, unless they were described as clinical targets. As far CAR-T cells, for clinically developed targets we used the previously defined list of targets as per MacKay et al^45^, while for pre-clinically developed targets we performed a PubMed search for all the genes in the surfaceome using the following search query: “gene AND CAR-T OR gene AND chimeric antigen receptor T-cells OR gene AND chimeric antigen receptor”, **Table S30**).

#### Monospecific CAR-T Cell Target TPI Calculation

To obtain the TPI for monospecific CAR-T targets, we used the list of targets defined based on the *TS*_*CAR*_ as described above. Then, for these targets we normalized the *TS*_*CAR*_ to assume values between 0 and 1 as follows:

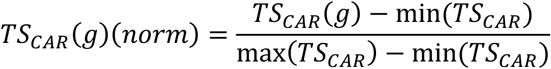

Where *TS*_*CAR*_(*g*) is the *TS*_*CAR*_ for a target *g*, and min(*TS*_*CAR*_) and max(*TS*_*CAR*_) represents the minimum and maximum *TS*_*CAR*_ values, respectively, for all the final targets being considered for the STS histotype analyzed.

Then, for each target *g*, we obtained a summative validation score, *V*_*CAR*_, calculated as the sum of different scores assigned for each validation feature, defined as follows:

- If the *H*_*RNA*_ score for *g* is equal or greater than the average *H*_*RNA*_ for *CD276* across the five STS studied, *g* is assigned a score of 2, otherwise *g* is assigned a score of 0. *CD276* was used as reference as it has been clinically (NCT04433221) and pre-clinically^95^ studied as a CAR-T target in STS.
- For each gene *g* and STS histotype we measured the percentage of samples in the STS proteomics Tang et al. dataset with medium-high (M-H) expression (z-score ≥ 0). DDLPS and WDLPS samples were considered as one group (LPS) for this analysis. If the percentage of samples of the STS analyzed with M-H expression for *g* was greater than or equal to the average percentage of samples with M-H expression for CD276 across SS, MFS, UPS, LMS and LPS, then we assigned a score of 2, otherwise we assigned a score of 0.
- If *g* was reported as target for clinically developed mAbs, ADCs, scFv, Fabs, nanobodies or fusion proteins, we assigned a score of 1. If instead *g* was only studied preclinically as target for therapeutic antibodies or ADCs, we assigned a score of 0.5. Otherwise, we assigned a score of 0.
- If *g* was reported as target for clinically developed CAR-T cells, we assigned a score of 1, while we assigned a score of 0.5 if *g* was only explored as a CAR-T target preclinically, and a score of 0 if it was never tested as a CAR-T target.
- If *g* resulted as essential (Chronos DepMap score ≤ – 1) in sarcoma cell lines derived from the STS histotype studied, we assigned a score of 1, otherwise we assigned a score of 0.5 if *g* was essential in any sarcoma cell line, or a score of 0 if *g* was not essential in any sarcoma cell lines.
- If *g* was involved in at least one gene fusion in the STS histotype studied we assigned a score of 1, otherwise we assigned a score of 0.5 if *g* was involved in at least one gene fusion in any STS, or a score of 0 if *g* was never involved in gene fusions in STS.
- If *g* was involved by at least one SNV or Indel in the STS histotype studied we assigned a score of 1, otherwise we assigned a score of 0.5 if *g* was involved by at least one SNV/Indel in any STS histotype, or a score of 0 if *g* was never involved by SNVs/Indels.
- If *g* was involved by CNAs in the STS histotype studied and there was a positive correlation between its linear CNV level and RNA expression in TCGA, we assigned a score of 1, otherwise if it was involved by CNAs either in TCGA or MSK-IMPACT in the same STS histotype studied we assigned a score of 0.5. Instead, if *g* was never involved by CNAs in STS we assigned a score of 0.

We then normalized *V*_*CAR*_ to assume values ranging from 0 to 1 as follows:

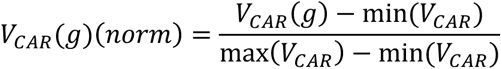

Finally, we calculated the TPI for each gene *g* as follows:

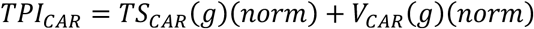

The TPI for the monospecific CAR-T targets is reported in **Table S31**.

#### Bispecific CAR-T Cells Targets TPI Calculation

For bispecific CAR-T cells targets we obtained a TPI for logic-gated ‘AND’ CAR-T cells (*TPI*_′*AND*′_), and a separate TPI for ‘AND-NOT’ CAR-T cells (*TPI*_′*AND*-*N*0*T*′_). Specifically, we defined suitable target pairs for both ‘AND’ and ‘AND-NOT’ CAR-T cells as described above, and then for each ‘AND’ CAR-T target and ‘AND-NOT’ CAR-T activator we calculated the average *TS*_′*AND*′_ and *TS*_′*AND*–*N*0*T*′_ scores considering all the target pairs in which that gene appears. We then normalized the average *TS*_′*AND*′_and *TS*_′*AND*–*N*0*T*′_ for each gene *g* to assume values ranging from 0 to 1 as follows:

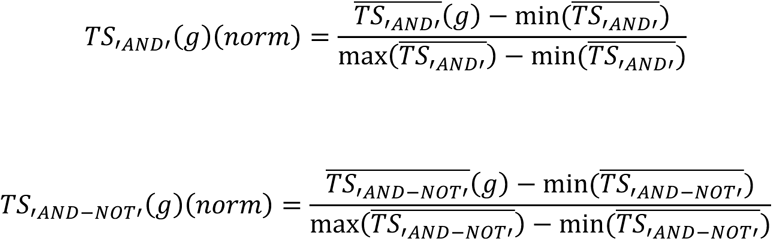

Then, for each target *g* we calculated *V*_*CAR*_(*g*) and *V*_*CAR*_(*g*)(*norm*) as described above, and finally obtained the TPIs as follows:

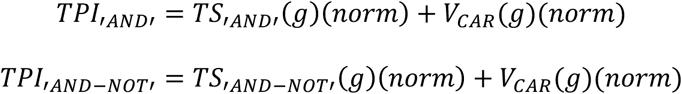

The TPIs for the bispecific CAR-T targets are reported in **Table S32**.

#### TCR and PC-CAR-T Cells Targets TPI Calculation

For each of the defined target antigens *g* for TCR-T and PC-CAR-T cells in each STS, we first normalized the *N*(*g*) score, indicative of the normal tissue and cell expression of *g* as previously defined, to assume values between 0 and 1:

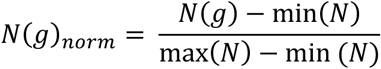

Then, for each target *g* we obtained a summative validation score, *V*_*TCR*/*PC*_, calculated as the sum of the following scores:

- If the *H*_*RNA*_ score for *g* is equal or greater than the *H*_*RNA*_ for *MAGEA4* in SS, *g* is assigned a score of 2, otherwise is assigned a score of 0. *MAGEA4* was used as reference in this case since it is the source antigen for the target of afami-cel, the first TCR-T cell therapy to receive FDA approval in soft tissue sarcomas.
- If *g* had a z-score ≥ 0 in at least two separate STS cell lines in either CCLE or in the ProCan-DepMapSanger datasets, we assigned a score of 1, otherwise we assigned a score of 0.
- If *g* resulted as essential (Chronos DepMap score ≤ – 1) in sarcoma cell lines derived from the STS histotype studied, we assigned a score of 1, otherwise we assigned a score of 0.5 if *g* was essential in any sarcoma cell line, or a score of 0 if *g* was not essential in any sarcoma cell lines.
- If *g* was involved in at least one gene fusion in the STS histotype studied we assigned a score of 1, otherwise we assigned a score of 0.5 if *g* was involved in at least one gene fusion in any STS, or a score of 0 if *g* was never involved in gene fusions in STS.
- If *g* was involved by at least one SNV or Indel in the STS histotype studied we assigned a score of 1, otherwise we assigned a score of 0.5 if *g* was involved by at least one SNV/Indel in any STS histotype, or a score of 0 if *g* was never involved by SNVs/Indels.
- If *g* was involved by CNAs in the STS histotype studied and there was a positive correlation between its linear CNV level and RNA expression in TCGA, we assigned a score of 1, otherwise if it was involved by CNAs either in TCGA or MSK-IMPACT in the same STS histotype studied we assigned a score of 0.5. Instead, if *g* was never involved by CNAs in STS we assigned a score of 0.

We then normalized *V*_*TCR*/*PC*_ to assume values ranging from 0 to 1 as follows:

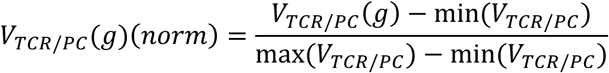

Next, for each target antigen we considered the target *pMHCs* defined based on their *TS*_*TCR*/*PC*_ as reported in **Table S27**. Specifically, for each target antigen *g* we calculated the average HLA allele frequency at each locus, by considering all the peptides derived from each antigen binding to HLA-A, HLA-B, or HLA-C class alleles, as follows:

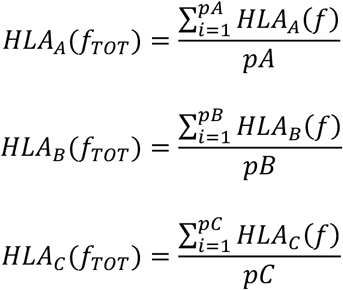

Where *pA*, *pB*, *pC* are respectively the number of *pMHC* pairs including class A, class B and class C alleles for the target *g*, while *HLA*_*A*_(*f*), *HLA*_*B*_(*f*), and *HLA*_*C*_(*f*) represent the HLA allele frequencies of the *i*^*th*^*pMHC* considered. We then obtained the total HLA allele frequency, *HLA*_*T*0*T*_(*g*) for each target *g* by adding *HLA*_*A*_(*f*_*T*0*T*_), *HLA*_*B*_(*f*_*T*0*T*_), and *HLA*_*C*_(*f*_*T*0*T*_), and normalized this to obtain values ranging from 0 to 1, as follows:

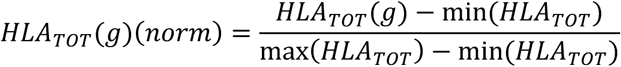

Next, for each target *g* we calculated the total number of *pMHC* targets (*pMHC*_*T*0*T*_) and the number of *pMHC* targets with *S*_*CR*_*Diff* ≤ 0 (*pMHC*_*SCR*_). We divided these by the target antigen total number of amino acids (*L*_*AA*_) to account for differences in protein size, and then normalized them to obtain values ranging from 0 to 1, as follows:

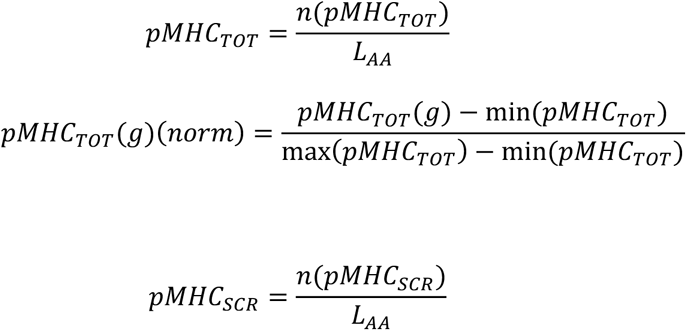

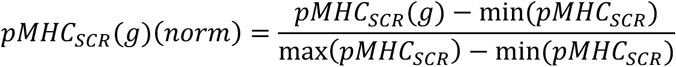

Finally, we calculated the TPI for each target *g* as follows:

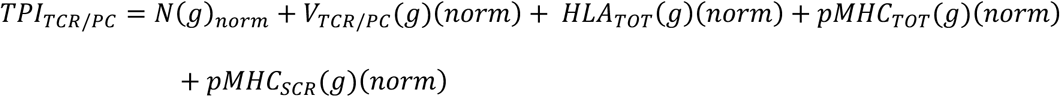

The TPIs for the TCR-T and PC-CAR-T cells targets are reported in **Table S33**.

