## Supplementary Figures 1-10 for "SCAN-ACT: Adoptive T Cell Therapy Target Discovery Through Single-Cell Transcriptomics"

This document includes **Supplementary Figures 1-10**.

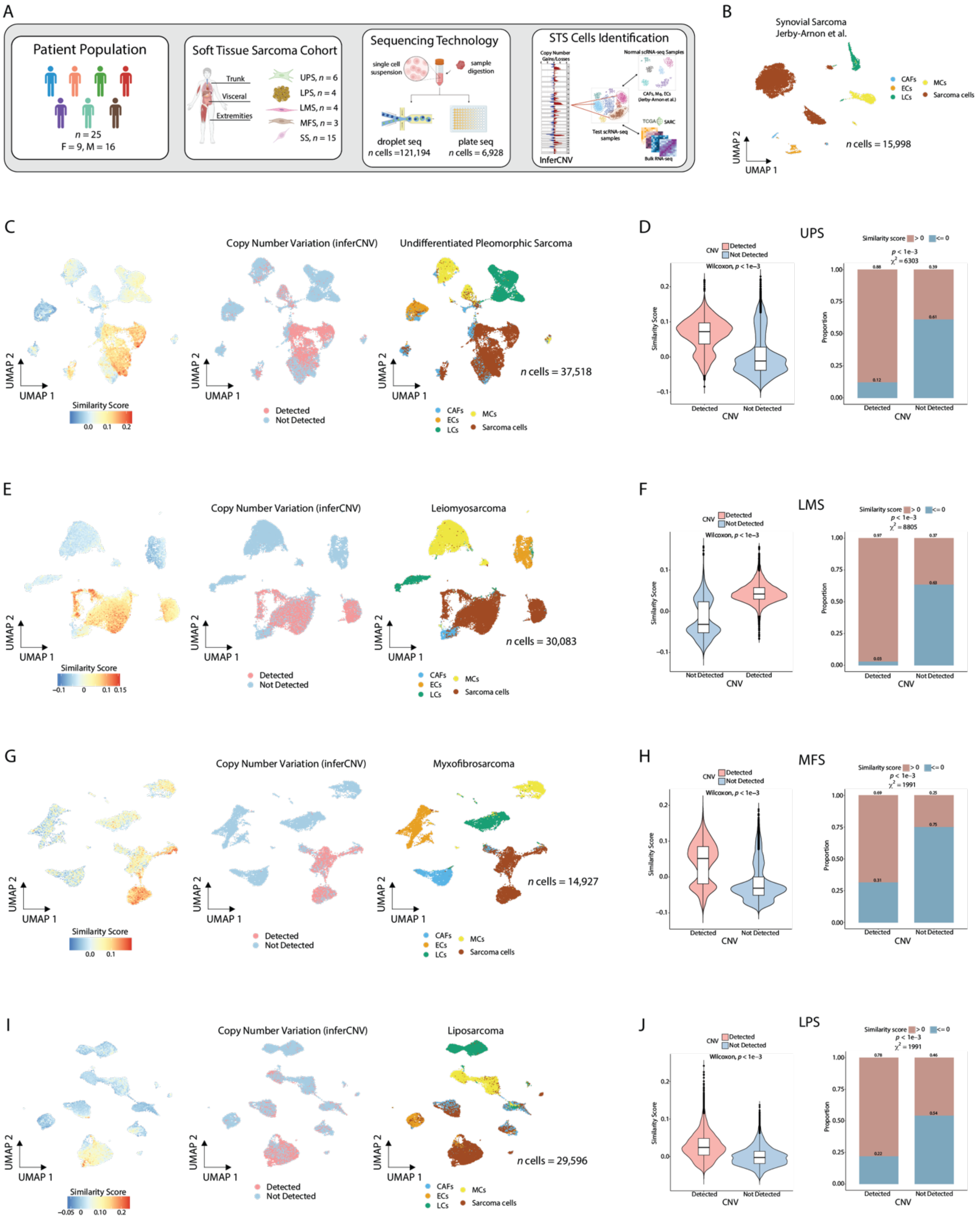

A
Patient Population
n = 25
F = 9, M = 16
Soft Tissue Sarcoma Cohort
Trunk
Visceral
Extremities
UPS, n = 6
LPS, n = 4
LMS, n = 4
MFS, n = 3
SS, n = 15
Sequencing Technology
single cell suspension
sample digestion
droplet seq
n cells =121,194
plate seq
n cells = 6,928
STS Cells Identification
InferCNV
B
Synovial Sarcoma
Jerby-Arnon et al.
CAFs
ECs
LCs
MCs
Sarcoma cells
n cells = 15,998
C
Copy Number Variation (inferCNV)
Undifferentiated Pleomorphic Sarcoma
Similarity Score
Detected
Not Detected
n cells = 37,518
D
UPS
E
Copy Number Variation (inferCNV)
Leiomyosarcoma
n cells = 30,083
F
LMS
G
Copy Number Variation (inferCNV)
Myxofibrosarcoma
n cells = 14,927
H
MFS
I
Copy Number Variation (inferCNV)
Liposarcoma
n cells = 29,596
J
LPS

**Figure S1**. **Single cell RNA sequencing of soft tissue sarcoma patient samples**.

**(A)** Scheme illustrating scRNA-seq of STS sarcoma samples and tumor cell annotation strategy. **(C, E, G, I)**. UMAP plots showing the samples in the study with cells colored based on the transcriptomic similarity score (*left*), CNV status (*center*), or assigned cell identity (*right*). **(B)** UMAP plot showing cell identity annotation for the SS samples. **(D**, **F**, **H**, **J)** Violin plots showing the distribution of the similarity score (left) and stacked bar charts showing the proportion of cells with similarity score $> 0$ and $\leq 0$ (right) for CD45- cells with and without CNVs detected by inferCNV in the UPS (**D**), LMS (**F**), MFS (**J**), and LPS (**H**) samples. A Wilcoxon rank-sum test was used to compare the two distributions in the violin plots and a Chi-square test was used to compare proportions in the bar charts.

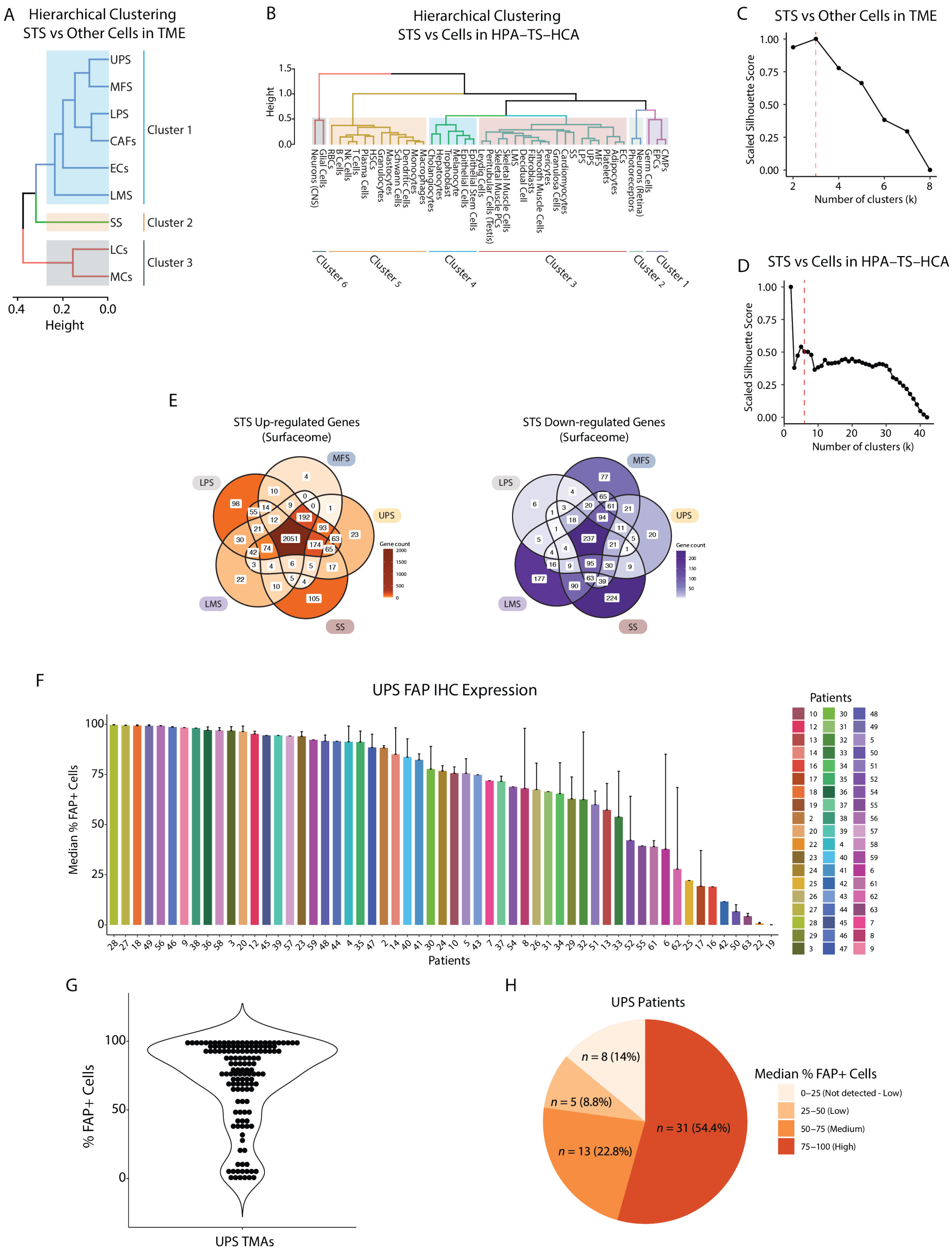

A
Hierarchical Clustering
STS vs Other Cells in TME
UPS
MFS
LPS
CAFs
ECs
LMS
SS
LCs
MCs
Cluster 1
Cluster 2
Cluster 3
0.4
0.2
0.0
Height
B
Hierarchical Clustering
STS vs Cells in HPA–TS–HCA
Height
1.5
1.0
0.5
0.0
Neurons (CNS)
Glial Cells
RBCs
B Cells
NK Cells
T Cells
Plasma Cells
HSCs
Granulocytes
Mastocytes
Schwann Cells
Dendritic Cells
Monocytes
Macrophages
Cholangiocytes
Hepatocytes
Trophoblast
Melanocyte
Epithelial Cells
Leydig Cells
Peritubular Cells (Testis)
Epithelial Stem Cells
Skeletal Muscle PCs
Skeletal Muscle Cells
LMS
Decidual Cell
Fibroblasts
Smooth Muscle Cells
Pericytes
Granulosa Cells
Cardiomyocytes
SS
LPS
UPS
MFS
Platelets
Adipocytes
ECs
Photoreceptors
Neurons (Retina)
Germ Cells
EPCs
CMPs
Cluster 6
Cluster 5
Cluster 4
Cluster 3
Cluster 2
Cluster 1
C
STS vs Other Cells in TME
Scaled Silhouette Score
1.00
0.75
0.50
0.25
0.00
2
4
6
8
Number of clusters (k)
D
STS vs Cells in HPA–TS–HCA
Scaled Silhouette Score
1.00
0.75
0.50
0.25
0.00
0
10
20
30
40
Number of clusters (k)
E
STS Up-regulated Genes
(Surfaceome)
STS Down-regulated Genes
(Surfaceome)
LPS
MFS
UPS
LMS
SS
Gene count
F
UPS FAP IHC Expression
Median % FAP+ Cells
100
75
50
25
0
Patients
G
% FAP+ Cells
100
50
0
UPS TMAs
H
UPS Patients
n = 8 (14%)
n = 5 (8.8%)
n = 13 (22.8%)
n = 31 (54.4%)
Median % FAP+ Cells
0–25 (Not detected - Low)
25–50 (Low)
50–75 (Medium)
75–100 (High)

**Figure S2**. **Analysis of differentially expressed genes in STS and FAP IHC expression in UPS.**

**(A)** Results of unsupervised hierarchical clustering of STS versus non-tumoral cells in the TME. The clustering analysis identified three cell clusters. **(B)** Results of unsupervised hierarchical clustering of STS versus normal cells in HPA-TS-HCA. The clustering analysis identified six cell clusters. **(C)** Plot showing the variation of the silhouette score for different number of clusters (k) for the comparisons of STS versus non-tumoral cells in the TME. **(D)** Plot showing the variation of the silhouette score for different number of clusters (k) for the comparisons of STS versus cells in the HPA-TS-HCA. **(E)** Venn diagrams showing the number of STS up-regulated and down-regulated genes shared between histotypes considering only surfaceome genes. **(F)** Bar chart showing the median % of cells expressing FAP as measured through IHC on 138 TMAs from 57 UPS patients. **(G)** Violin plot showing the distribution of the % of $FAP^{+}$ cells in the 138 UPS TMAs obtained from 57 patients. **(H)** Pie chart showing the fraction of patients with Not Detected-Low FAP expression (0-25% $FAP^{+}$ cells), Low expression (25-50 % $FAP^{+}$ cells), Medium expression (50-75 % $FAP^{+}$ cells), and High expression (75-100 % $FAP^{+}$ cells). Expression is measured as the median value considering all the TMAs for each patient.

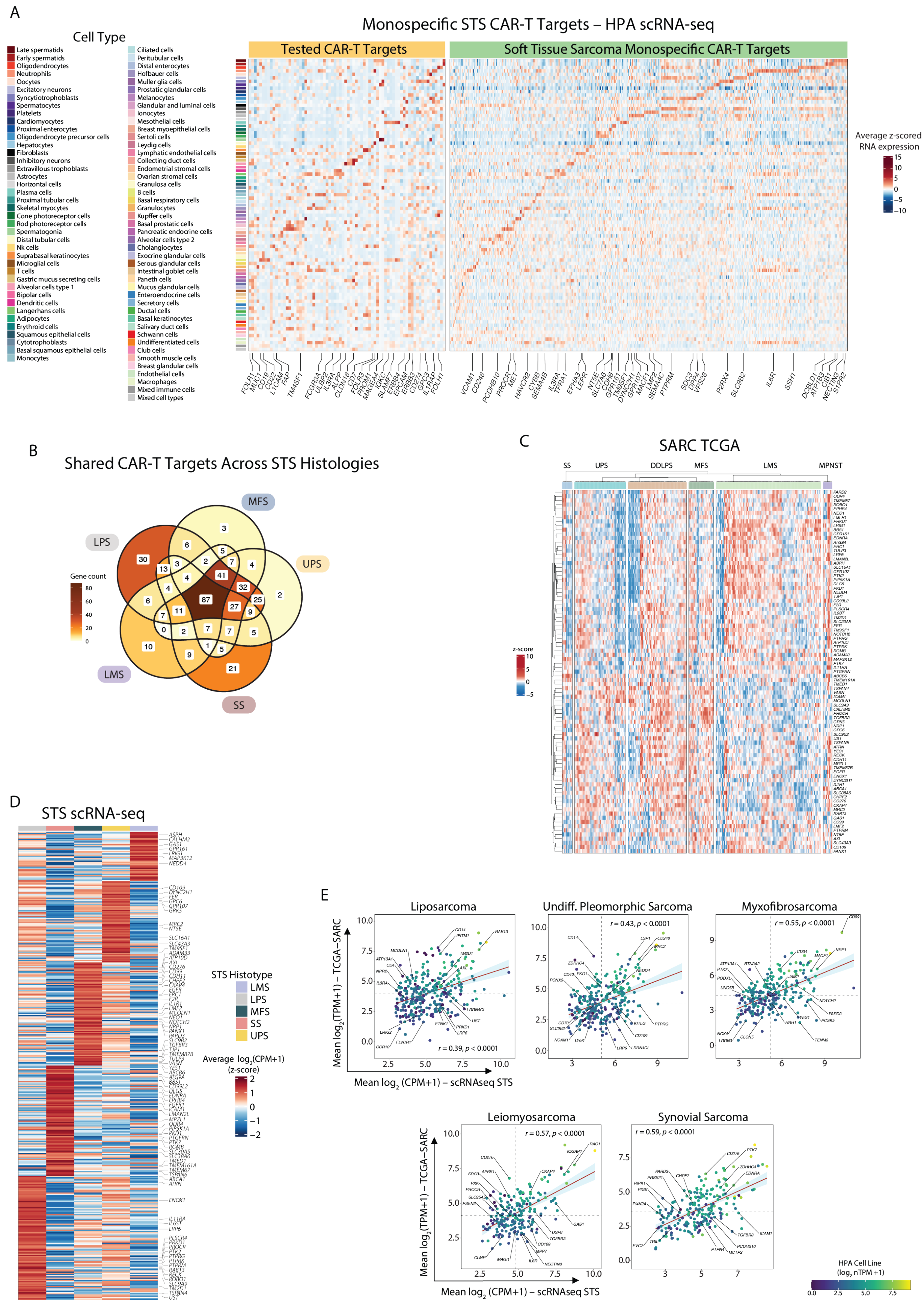
A
Monospecific STS CAR-T Targets – HPA scRNA-seq
Cell Type
Tested CAR-T Targets
Soft Tissue Sarcoma Monospecific CAR-T Targets
Average z-scored RNA expression
B
Shared CAR-T Targets Across STS Histologies
MFS
LPS
UPS
LMS
SS
Gene count
C
SARC TCGA
SS
UPS
DDLPS
MFS
LMS
MPNST
z-score
D
STS scRNA-seq
STS Histotype
LMS
LPS
MFS
SS
UPS
E
Liposarcoma
Undiff. Pleomorphic Sarcoma
Myxofibrosarcoma
Leiomyosarcoma
Synovial Sarcoma
Mean log2(TPM+1) – TCGA–SARC
Mean log2 (CPM+1) – scRNAseq STS
HPA Cell Line (log2 nTPM +1)

**Figure S3**. **Monospecific CAR-T targets**.

**(A)** scRNA-seq expression of monospecific CAR targets in the HPA scRNA-seq database. Heatmap shows the average z-scored and median-centered pseudo-bulk RNA expression for tested CAR-T cells targets (upper panel) and the monospecific STS CAR targets identified in this study (bottom panel) as measured in the HPA scRNA-seq database. RNA expression values were first pseudo-bulked by cell cluster, and then averaged across clusters and tissues for a given cell type. **(B)** Venn diagram showing the number of targets shared between the five STS histotypes. **(C)** Expression of the monospecific CAR targets common to MFS, LMS, SS, LPS, and UPS in TCGA SARC. RNA expression in $\log_2$(TPM+1) was median-centered across samples and z-scored across genes before plotting. **(D)** Expression of the candidate monospecific CAR targets in scRNA-seq STS samples. RNA expression in $\log_2$(CPM+1) was averaged across cells of the same type before median-centering and z-scoring. **(E)** Pearson correlation between average scRNA-seq expression and average bulk RNA-seq in matched STS samples from TCGA SARC, for CAR-T targets in each STS. Dots are colored based on gene expression in HPA sarcoma cell lines. CMPs: Common Myeloid Progenitor Cells; EPCs: Erythroid Progenitor Cells; HSCs: Hematopoietic Stem Cells; RBCs: Red blood cells; ECs: Endothelial Cells; MCs: Myeloid Cells; LCs: Lymphoid Cells; CAFs: Cancer associated fibroblasts.

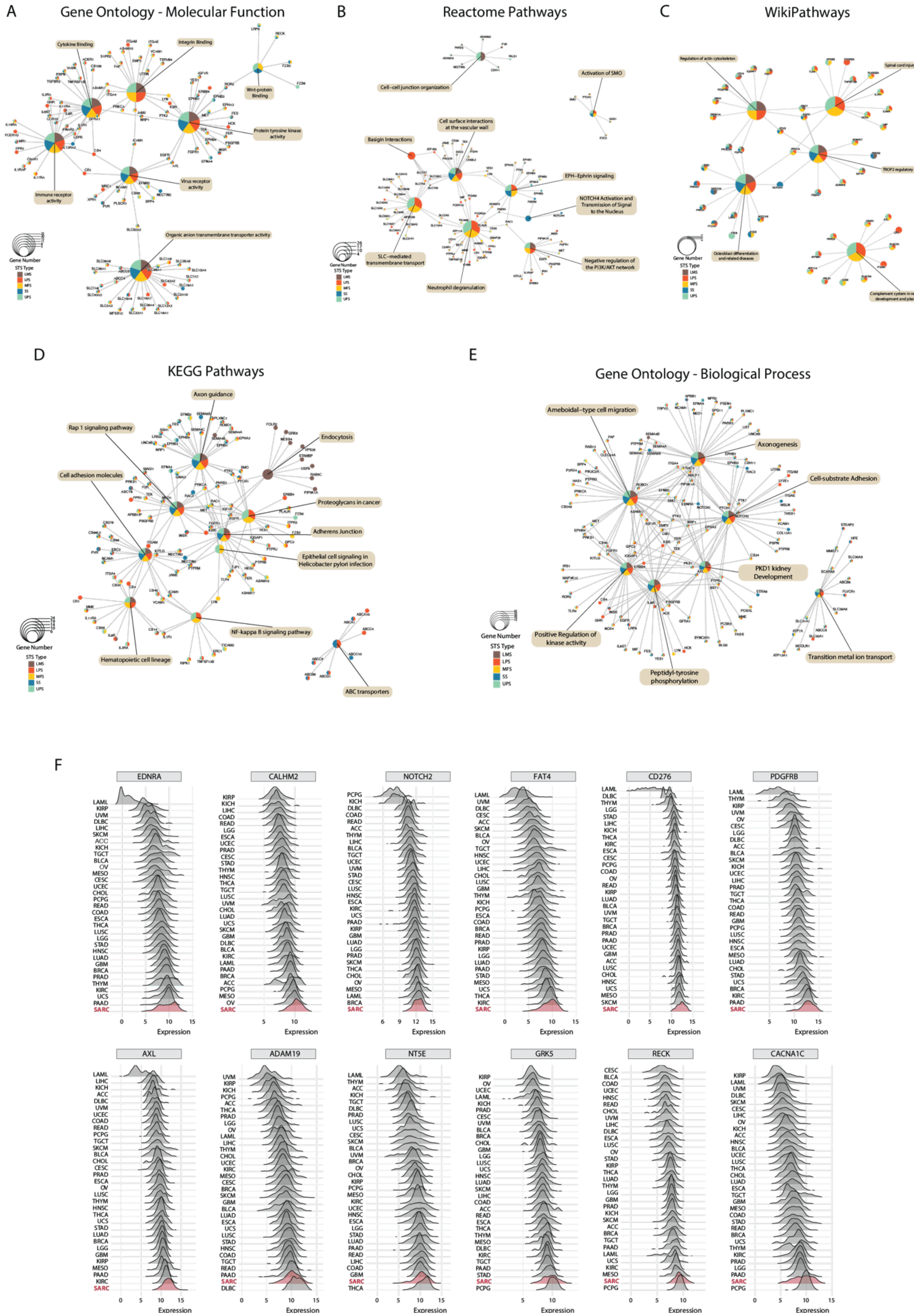
A
Gene Ontology - Molecular Function
Cytokine Binding
Integrin Binding
Wnt-protein Binding
Protein tyrosine kinase activity
Immune receptor activity
Virus receptor activity
Organic anion transmembrane transporter activity
B
Reactome Pathways
Cell–cell junction organization
Activation of SMO
Cell surface interactions at the vascular wall
Basigin Interactions
EPH–Ephrin signaling
NOTCH4 Activation and Transmission of Signal to the Nucleus
SLC–mediated transmembrane transport
Negative regulation of the PI3K/AKT network
Neutrophil degranulation
C
WikiPathways
Regulation of actin cytoskeleton
Spinal cord injury
TROP2 regulatory signaling
Osteoblast differentiation and related diseases
Complement system in neuronal development and plasticity
D
KEGG Pathways
Axon guidance
Rap 1 signaling pathway
Endocytosis
Cell adhesion molecules
Proteoglycans in cancer
Adherens Junction
Epithelial cell signaling in Helicobacter pylori infection
NF-kappa B signaling pathway
Hematopoietic cell lineage
ABC transporters
E
Gene Ontology - Biological Process
Ameboidal–type cell migration
Axonogenesis
Cell-substrate Adhesion
PKD1 kidney Development
Positive Regulation of kinase activity
Peptidyl-tyrosine phosphorylation
Transition metal ion transport
Gene Number
STS Type
LMS
LPS
MFS
SS
UPS
F
EDNRA
CALHM2
NOTCH2
FAT4
CD276
PDGFRB
AXL
ADAM19
NT5E
GRK5
RECK
CACNA1C
SARC
Expression

**Figure S4. Gene set over-representation analysis (ORA) of monospecific CAR-T targets by sarcoma histotype and TCGA Pan-Cancer Atlas expression for selected monospecific CAR-T targets.**

**(A-E)** Gene concept networks showing results of gene set ORA using **(A)** GO Molecular Function terms **(B)**, Kyoto Encyclopedia of Genes and Genomes (KEGG) pathways, **(C)** WikiPathways terms, **(D)** Reactome pathways, and GO Biological Process terms **(E)**. (**F**) Ridge plots showing RNA expression in $log_2$(TPM+1) across different cancers in the Pan-Cancer Atlas from TCGA for selected monospecific CAR-T cells targets. Soft tissue sarcoma samples (SARC) are highlighted in bold red.

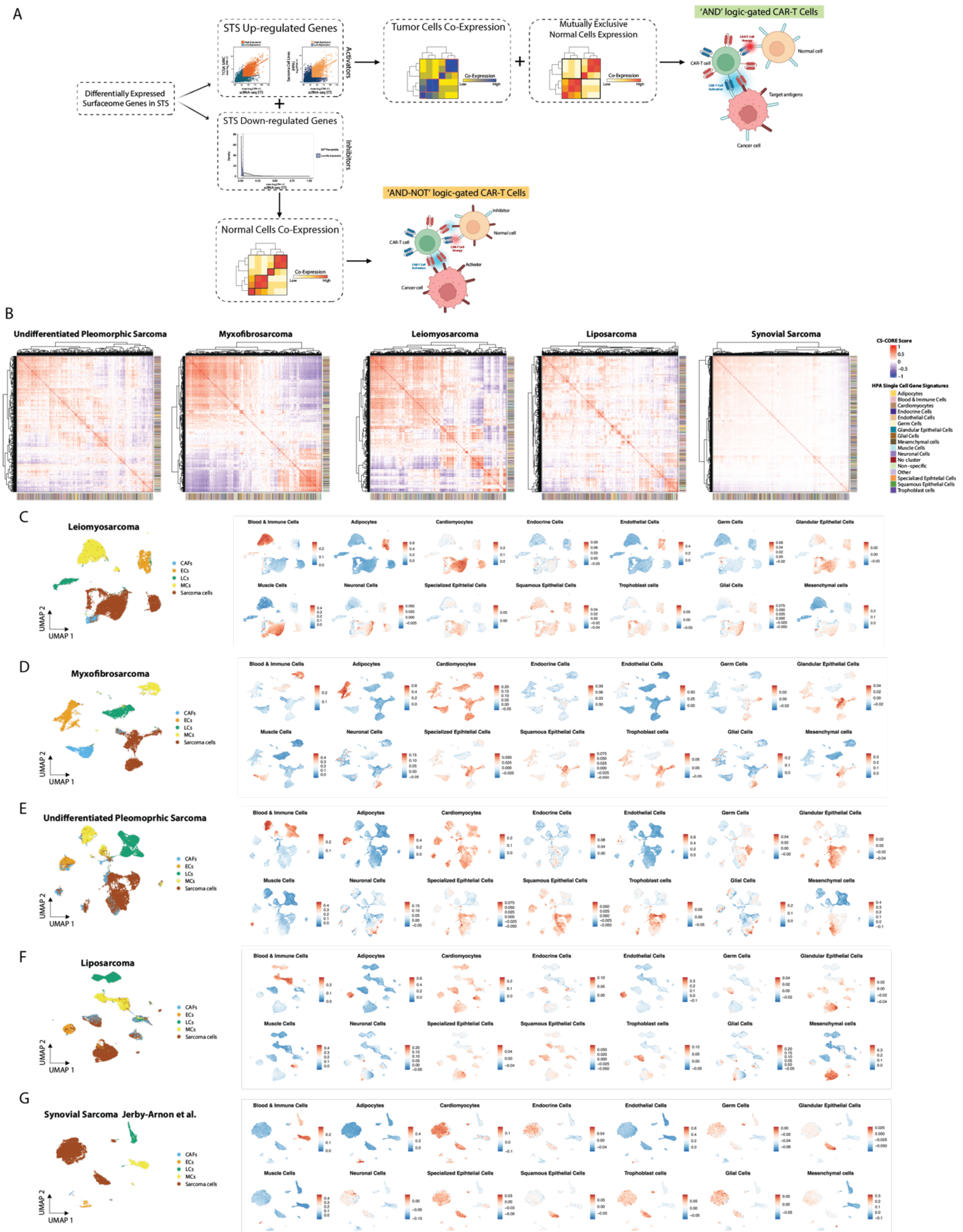
A
Differentially Expressed Surfaceome Genes in STS
STS Up-regulated Genes
Activators
STS Down-regulated Genes
Inhibitors
Tumor Cells Co-Expression
Mutually Exclusive Normal Cells Expression
'AND' logic-gated CAR-T Cells
Normal cell
CAR-T cell
Target antigens
Cancer cell
Normal Cells Co-Expression
'AND-NOT' logic-gated CAR-T Cells
Inhibitor
Activator
B
Undifferentiated Pleomorphic Sarcoma
Myxofibrosarcoma
Leiomyosarcoma
Liposarcoma
Synovial Sarcoma
CS-CORE Score
HPA Single Cell Gene Signatures
C
Leiomyosarcoma
D
Myxofibrosarcoma
E
Undifferentiated Pleomoprhic Sarcoma
F
Liposarcoma
G
Synovial Sarcoma Jerby-Arnon et al.
CAFs
ECs
LCs
MCs
Sarcoma cells
UMAP 2
UMAP 1
Blood & Immune Cells
Adipocytes
Cardiomyocytes
Endocrine Cells
Endothelial Cells
Germ Cells
Glandular Epithelial Cells
Muscle Cells
Neuronal Cells
Specialized Epithelial Cells
Squamous Epithelial Cells
Trophoblast cells
Glial Cells
Mesenchymal cells

**Figure S5**. **Bispecific CAR-T cell targets in STS**.

**(A)** Illustration showing the steps of the bispecific CAR targets pair discovery pipeline. **(B)** Heatmaps showing the CS-CORE scores for the highly expressed STS up-regulated genes used to define the 'AND' CAR gene pairs. Columns are colored based on the HPA single-cell signature to which each gene belongs to. **(C-G)** UMAP plots showing the expression of cell-specific gene signatures in LMS **(C)**, MFS **(D)**, UPS **(E)**, LPS **(F)**, and SS **(G)**. The gene signatures used represent all the genes belonging to a given HPA scRNA-seq cell type-specific signature.

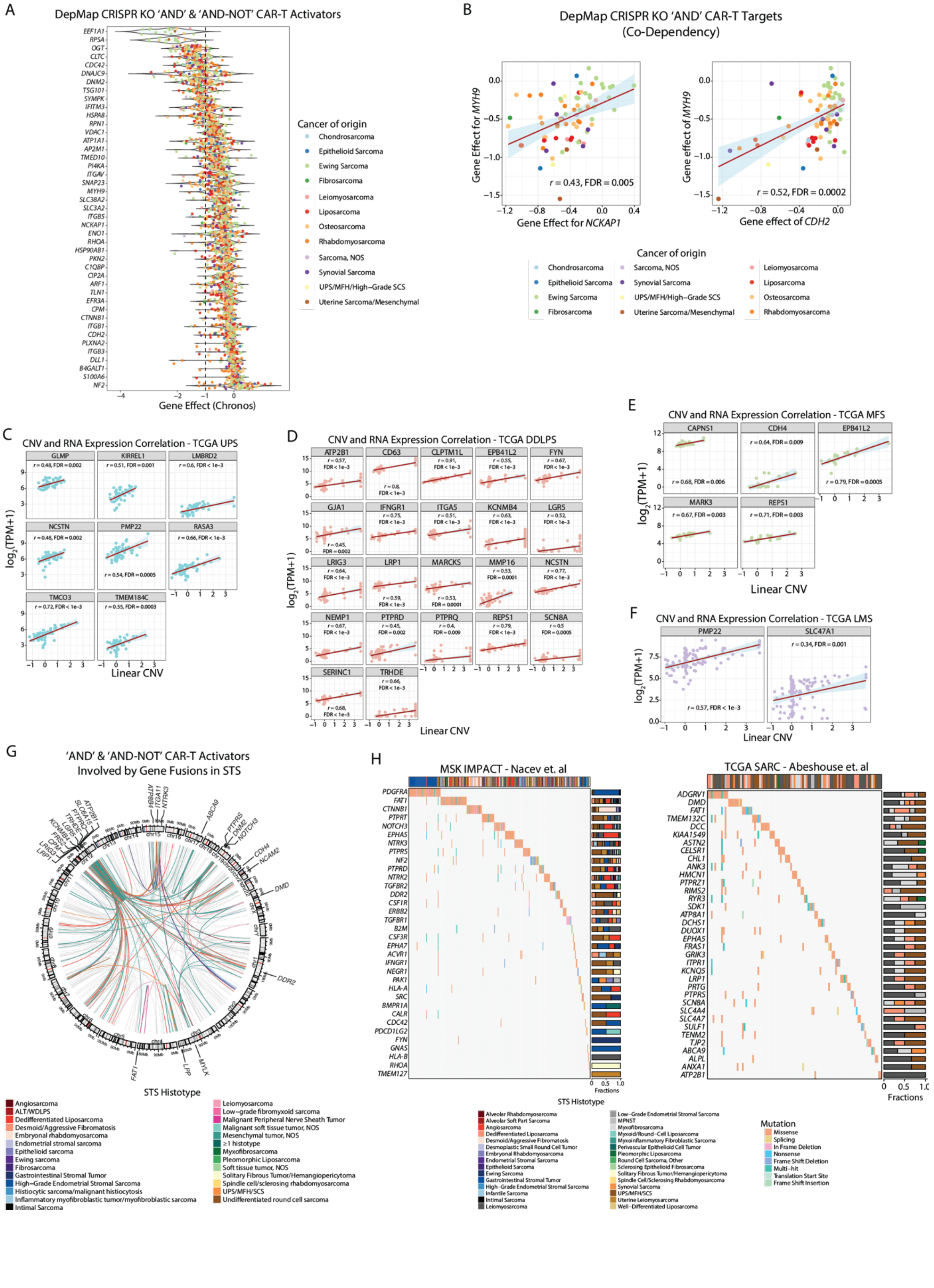
A
DepMap CRISPR KO 'AND' & 'AND-NOT' CAR-T Activators
EEF1A1
RPSA
OGT
CLTC
CDC42
DNAJC9
DNM2
TSG101
SYMPK
IFITM3
HSPA8
RPN1
VDAC1
ATP1A1
AP2M1
TMED10
PI4KA
ITGAV
SNAP23
MYH9
SLC38A2
SLC3A2
ITGB5
NCKAP1
ENO1
RHOA
HSP90AB1
PKN2
C1QBP
CIP2A
ARF1
TLN1
EFR3A
CPM
CTNNB1
ITGB1
CDH2
PLXNA2
ITGB3
DLL1
B4GALT1
S100A6
NF2
Gene Effect (Chronos)
Cancer of origin
Chondrosarcoma
Epithelioid Sarcoma
Ewing Sarcoma
Fibrosarcoma
Leiomyosarcoma
Liposarcoma
Osteosarcoma
Rhabdomyosarcoma
Sarcoma, NOS
Synovial Sarcoma
UPS/MFH/High-Grade SCS
Uterine Sarcoma/Mesenchymal
B
DepMap CRISPR KO 'AND' CAR-T Targets
(Co-Dependency)
Gene Effect for MYH9
Gene Effect for NCKAP1
r = 0.43, FDR = 0.005
Gene effect of MYH9
Gene effect of CDH2
r = 0.52, FDR = 0.0002
Cancer of origin
Chondrosarcoma
Epithelioid Sarcoma
Ewing Sarcoma
Fibrosarcoma
Sarcoma, NOS
Synovial Sarcoma
UPS/MFH/High-Grade SCS
Uterine Sarcoma/Mesenchymal
Leiomyosarcoma
Liposarcoma
Osteosarcoma
Rhabdomyosarcoma
C
CNV and RNA Expression Correlation - TCGA UPS
GLMP
KIRREL1
LMBRD2
NCSTN
PMP22
RASA3
TMCO3
TMEM184C
log2(TPM+1)
Linear CNV
D
CNV and RNA Expression Correlation - TCGA DDLPS
ATP2B1
CD63
CLPTM1L
EPB41L2
FYN
GJA1
IFNGR1
ITGA5
KCNMB4
LGR5
LRIG3
LRP1
MARCKS
MMP16
NCSTN
NEMP1
PTPRD
PTPRQ
REPS1
SCN8A
SERINC1
TRHDE
log2(TPM+1)
Linear CNV
E
CNV and RNA Expression Correlation - TCGA MFS
CAPNS1
CDH4
EPB41L2
MARK3
REPS1
log2(TPM+1)
Linear CNV
F
CNV and RNA Expression Correlation - TCGA LMS
PMP22
SLC47A1
r = 0.34, FDR = 0.001
r = 0.57, FDR < 1e-3
log2(TPM+1)
Linear CNV
G
'AND' & 'AND-NOT' CAR-T Activators
Involved by Gene Fusions in STS
STS Histotype
Angiosarcoma
ALT/WDLPS
Dedifferentiated Liposarcoma
Desmoid/Aggressive Fibromatosis
Embryonal rhabdomyosarcoma
Endometrial stromal sarcoma
Epithelioid sarcoma
Ewing sarcoma
Fibrosarcoma
Gastrointestinal Stromal Tumor
High-Grade Endometrial Stromal Sarcoma
Histiocytic sarcoma/malignant histiocytosis
Inflammatory myofibroblastic tumor/myofibroblastic sarcoma
Intimal Sarcoma
Leiomyosarcoma
Low-grade fibromyxoid sarcoma
Malignant Peripheral Nerve Sheath Tumor
Malignant soft tissue tumor, NOS
Mesenchymal tumor, NOS
≥1 histotype
Myxofibrosarcoma
Pleomorphic Liposarcoma
Soft tissue tumor, NOS
Solitary Fibrous Tumor/Hemangiopericytoma
Spindle cell/sclerosing rhabdomyosarcoma
UPS/MFH/SCS
Undifferentiated round cell sarcoma
H
MSK IMPACT - Nacev et. al
PDGFRA
FAT1
CTNNB1
PTPRT
NOTCH3
EPHA5
NTRK3
PTPRS
NF2
PTPRD
NTRK2
TGFBR2
DDR2
CSF1R
ERBB2
TGFBR1
B2M
CSF3R
EPHA7
ACVR1
IFNGR1
NEGR1
PAK1
HLA-A
SRC
BMPR1A
CALR
CDC42
PDCD1LG2
FYN
GNAS
HLA-B
RHOA
TMEM127
Fractions
TCGA SARC - Abeshouse et. al
ADGRV1
DMD
FAT1
TMEM132C
DCC
KIAA1549
ASTN2
CELSR1
CHL1
ANK3
HMCN1
PTPRZ1
RIMS2
RYR3
SDK1
ATP8A1
DCHS1
DUOX1
EPHA5
FRAS1
GRIK3
ITPR1
KCNQ5
LRP1
PRTG
PTPRS
SCN8A
SLC4A4
SLC4A7
SULF1
TENM2
TJP2
ABCA9
ALPL
ANXA1
ATP2B1
Fractions
STS Histotype
Alveolar Rhabdomyosarcoma
Alveolar Soft Part Sarcoma
Angiosarcoma
Dedifferentiated Liposarcoma
Desmoid/Aggressive Fibromatosis
Desmoplastic Small Round Cell Tumor
Embryonal Rhabdomyosarcoma
Endometrial Stromal Sarcoma
Epithelioid Sarcoma
Ewing Sarcoma
Gastrointestinal Stromal Tumor
High-Grade Endometrial Stromal Sarcoma
Infantile Sarcoma
Intimal Sarcoma
Leiomyosarcoma
Low-Grade Endometrial Stromal Sarcoma
MPNST
Myxofibrosarcoma
Myxoid/Round-Cell Liposarcoma
Myxoinflammatory Fibroblastic Sarcoma
Perivascular Epithelioid Cell Tumor
Pleomorphic Liposarcoma
Round Cell Sarcoma, Other
Sclerosing Epithelioid Fibrosarcoma
Solitary Fibrous Tumor/Hemangiopericytoma
Spindle Cell/Sclerosing Rhabdomyosarcoma
Synovial Sarcoma
UPS/MFH/SCS
Uterine Leiomyosarcoma
Well-Differentiated Liposarcoma
Mutation
Missense
Splicing
In Frame Deletion
Nonsense
Frame Shift Deletion
Multi-hit
Translation Start Site
Frame Shift Insertion

**Figure S6. Functional characterization of bispecific CAR-T cell targets in STS**.

**(A)** Violin plot showing the effect on cell growth of the knock-out (KO) of 'AND' CAR targets and 'AND-NOT' CAR activators in sarcoma cell lines as reported in the DepMap public database. Only targets that are essential for cell growth in at least one sarcoma cell line are shown. The gene KO effect is quantified based on the Chronos algorithm with more negative values corresponding to genes that are more essential for cell growth. The vertical dashed line intersecting at –1 indicates the median Chronos value for the known essential genes. **(B)** Scatter plots showing CRISPR KO co-dependency plots for the 'AND' CAR pairs *MYH9-NCKAP* (left) and *MYH9-NCKAP* (right). Pearson correlation was used, and the p-values were corrected for multiple hypothesis testing using the BH method with significance set to FDR < 0.01. **(C**, **D**, **E**, **F)** Scatter plots showing the Pearson correlation between linear CNV level and RNA expression for UPS (**C**), DDLPS (**D**), MFS (**E**), LMS (**F**) samples in TCGA for selected 'AND' CAR and 'AND-NOT' CAR (activators) targets. RNA expression is reported in $\log_2$(TPM+1). The p-values shown were adjusted for multiple hypothesis testing using the BH method with significance set to FDR < 0.01. **(G)** Circos plot showing gene fusions in STS from the Mitelman database, MSK-IMPACT, and TCGA SARC cohorts, for which one of the involved genes is a bispecific CAR target. Colored links show fusions involving the top 30 genes most recurrently involved in gene fusions among all bispecific CAR targets. The link colors indicate the STS histotype in which the fusion has been identified. **(H)** Oncoplots showing recurring SNVs/Indels for bispecific CAR targets in the MSK-IMPACT (*left*) and TCGA cohorts (*right*). The stacked bar charts show the mutation frequency per each gene across different soft tissue sarcoma histologies. Only the most frequently mutated genes across all samples in each cohort are shown. ALT: Atypical Lyopmatous Tumor; WDLPS: Well-differentiated Liposarcoma; GIST: Gastrointestinal Stromal Tumor; MPNST: Malignant Peripheral Nerve Sheath Tumor; IMT: Inflammatory Myofibroblastic Tumor; UPS:

Undifferentiated Pleomorphic Sarcoma; MFH: Malignant Fibrous Histiocytoma; SCS: High-grade Spindle Cell Sarcoma.

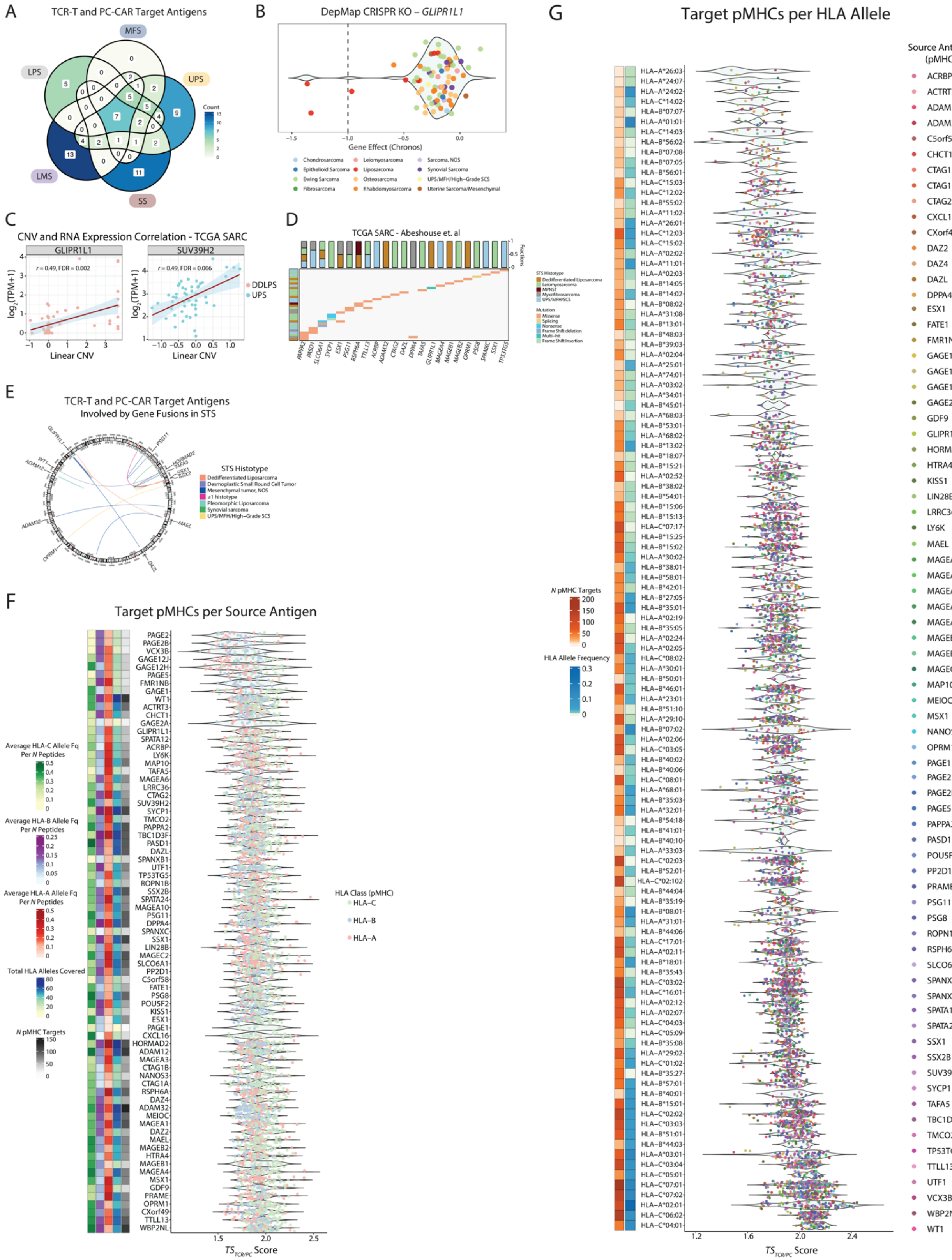

A
TCR-T and PC-CAR Target Antigens
B
DepMap CRISPR KO – GLIPR1L1
Gene Effect (Chronos)
C
CNV and RNA Expression Correlation - TCGA SARC
Linear CNV
D
TCGA SARC - Abeshouse et. al
E
TCR-T and PC-CAR Target Antigens
Involved by Gene Fusions in STS
STS Histotype
F
Target pMHCs per Source Antigen
HLA Class (pMHC)
G
Target pMHCs per HLA Allele
Source Antigen (pMHC)

**Figure S7**. **Characterization of TCR-T and PC-CAR target antigens and derived target pMHCs**.

**(A)** Venn diagram showing the number of PC-CAR and TCR-T target antigens shared between the different STS studied. **(B)** Violin plot showing the CRISPR KO effect of *GLIPR1L1* on STS cell lines. The gene KO effect is quantified using the Chronos algorithm with more negative values corresponding to genes more essential for cell growth. The vertical dashed line intersecting at –1 indicates the median Chronos value for known essential genes. Dots are colored based on the cell line cancer of origin. **(C)** Scatter plots showing Pearson correlation between CNA level and RNA expression in TCGA DDLPS and UPS samples for *GLIPR1L1* and *SUV39H2*, respectively. P-values were adjusted for multiple hypothesis testing using the BH method, and significance was set at FDR < 0.01. **(D)** Oncoplot showing recurring SNVs/Indels for TCR-T and PC-CAR target antigens in TCGA SARC. The stacked bar charts show the mutation frequency per each gene across different STS histotypes. Only the most frequently mutated genes across all samples are shown. **(E)** Circos plot showing gene fusions in STS from the Mitelman database, MSK-IMPACT and TCGA SARC, for which one of the involved genes is a TCR-T and PC-CAR target antigen. The link colors indicate the STS histotype in which the fusion was identified. **(F)** Violin plot showing the distribution of $TS_{TCR/PC}$ scores for the candidate $pMHCs$ selected for each antigen. Dots are colored based on the HLA class restriction of each $pMHC$. The heatmap on the left shows the total number of target $pMHCs$ per antigen, the total number of HLA alleles covered, and the cumulative HLA-A, HLA-B, and HLA-C allele frequency for each antigen. The cumulative allele frequencies for each HLA locus were calculated by first isolating all $pMHC$ pairs that bind to HLA alleles of a given locus (A, B, or C) for each antigen. Then, for each locus, the HLA allele frequencies of these $pMHCs$ were summed and divided by the number of unique peptides that form all the $pMHCs$ binding to HLA alleles of that locus. **(G)** Violin plot showing the distribution of $TS_{TCR/PC}$ scores for the candidate $pMHCs$ grouped by their HLA allele restriction. The dots are colored based on the source tumor antigen of the $pMHCs$. The heatmap on the left shows the total number of $pMHCs$ per each HLA allele, and the average worldwide allele frequency for each HLA allele.

A

### Peptide Similarity Analysis of Known Cross-Reactive Peptide Pairs

| Source Antigen | Target Peptide | Cross-Reactive Peptide | Normalized PMBEC Similarity | Normalized AAindex Similarity | Peptide Similarity Score | Cross-Reactive Peptide Organism | HLA |
|---|---|---|---|---|---|---|---|
| MAGEA-6 | YIFATCLGLS | YIFAACLLLI | 1 | 1 | 2 | Mycobactermium Penetrantis | HLA-A*02:01 |
| C1QTNF12 | LLGPQLVLL | LLGPLLVLL | 0.908869595 | 0.88228284 | 1.791152436 | HBV | HLA-A*02:01 |
| EPCAM | VVAGIVVLV | VVAGIIALV | 0.913012115 | 0.770224445 | 1.68323656 | HIV-1 | HLA-A*02:01 |
| MDK | ALLALTSAV | ALMAFTSAV | 0.912451862 | 0.766779325 | 1.679231188 | HCV | HLA-A*02:01 |
| MDK | LLLTLLALL | LLLTLLLLL | 0.789303717 | 0.884015257 | 1.673318974 | Adenovirus | HLA-A*02:01 |
| DPYSL2 | IPRRTTQRI | IPRRIRQRI | 0.8906457 | 0.769828303 | 1.660474003 | HIV-1 | HLA-B*07:02 |
| gp100 | MLGTHTMEV | MLGTHAMLV | 0.8710389 | 0.770499778 | 1.641538678 | HCMV | HLA-A*02:01 |
| NDUFS2 | VSDGSSRPY | SSDNSSRPY | 0.869510415 | 0.767433998 | 1.636944413 | HIV-1 | HLA-A*01:01 |
| HEPACAM | RLAPFVYLL | RLAPFGYKI | 0.851321611 | 0.677718908 | 1.529040519 | Encephalitis virus | HLA-A*02:01 |
| CD274 | LLNAFTVTV | LLNAFAIAV | 0.86741129 | 0.65556493 | 1.52297622 | HIV-1 | HLA-A*02:01 |
| TERT | RLVDDFLLV | RLVDEFLAI | 0.836655457 | 0.679907523 | 1.516562981 | HIV-1 | HLA-A*02:01 |
| ISG15 | MLAGNEFQV | MLAGNAFTA | 0.81887112 | 0.680004889 | 1.498876008 | Calicivirus | HLA-A*02:01 |
| MART-1 | AAGIGILTV | IAGIGILAI | 0.772540075 | 0.679898621 | 1.452438696 | Pseudorabies | HLA-A*02:01 |
| CEA | IMIGVLVGV | IMVGALIGV | 0.791737024 | 0.650953595 | 1.442690619 | HIV-1 | HLA-A*02:01 |
| MART-1 | AAGIGILTV | GAGIGVLTA | 0.731202806 | 0.682462736 | 1.413665542 | Bacillus Polymyxa | HLA-A*02:01 |
| MART-1 | AAGIGILTV | QAGIGILLA | 0.720063659 | 0.679487922 | 1.399551581 | E. Coli | HLA-A*02:01 |
| CNIH4 | LLNLPVATW | HCNLSVATW | 0.665910794 | 0.674457652 | 1.340368446 | HIV-1 | HLA-B*58:01 |
| CCR9 | AIADLLFLV | RLRDLLFLV | 0.613536037 | 0.677826374 | 1.291362411 | HIV-1 | HLA-A*02:01 |
| TMEM161A | ALGGLLTPL | CLGGLLTMV | 0.551620235 | 0.686133548 | 1.237753783 | EBV | HLA-A*02:01 |
| TMEM161A | ALGGLLTPL | LLGGLLTMV | 0.530080979 | 0.686820687 | 1.216901666 | E. Coli | HLA-A*02:01 |
| MAGEA-3 | EVDPIGHLY | ESDPIVAQY | 0.649505777 | 0.54274076 | 1.192246536 | Homo Sapiens (Titin) | HLA-A*01:01 |
| MART-1 | AAGIGILTV | GAGIGVAVL | 0.59185572 | 0.457640834 | 1.049496554 | HSV-2 | HLA-A*02:01 |
| MART-1 | AAGIGILTV | GIGIGVLAA | 0.559592626 | 0.472963967 | 1.032556593 | HSV-1 | HLA-A*02:01 |
| MART-1 | AAGIGILTV | LIVIGILIL | 0.521802407 | 0.475771354 | 0.997573761 | Adenovirus | HLA-A*02:01 |
| MART-1 | EAAGIGILTV | MLSGIGIFFI | 0.523095177 | 0.425643599 | 0.948738776 | Chlamydia Trachomatis | HLA-A*02:01 |

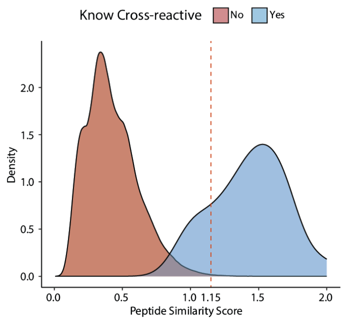


B

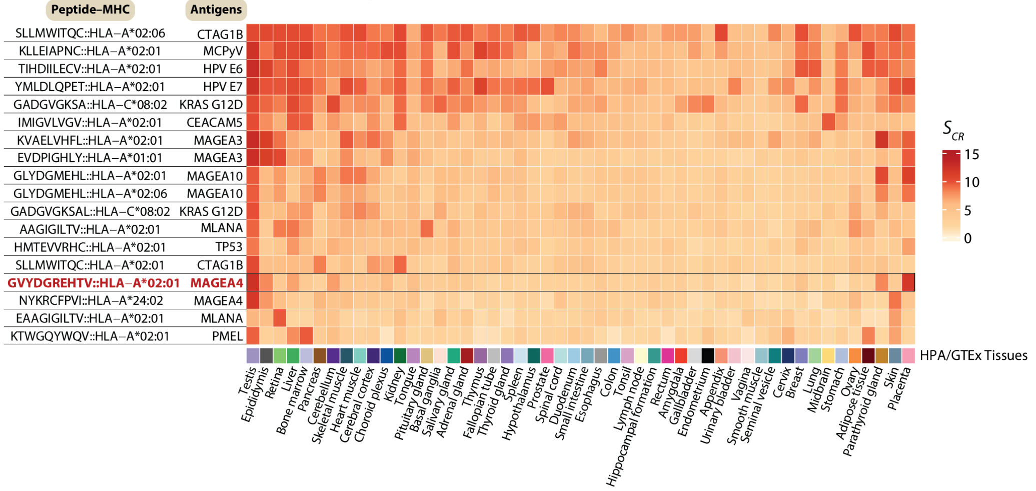


C

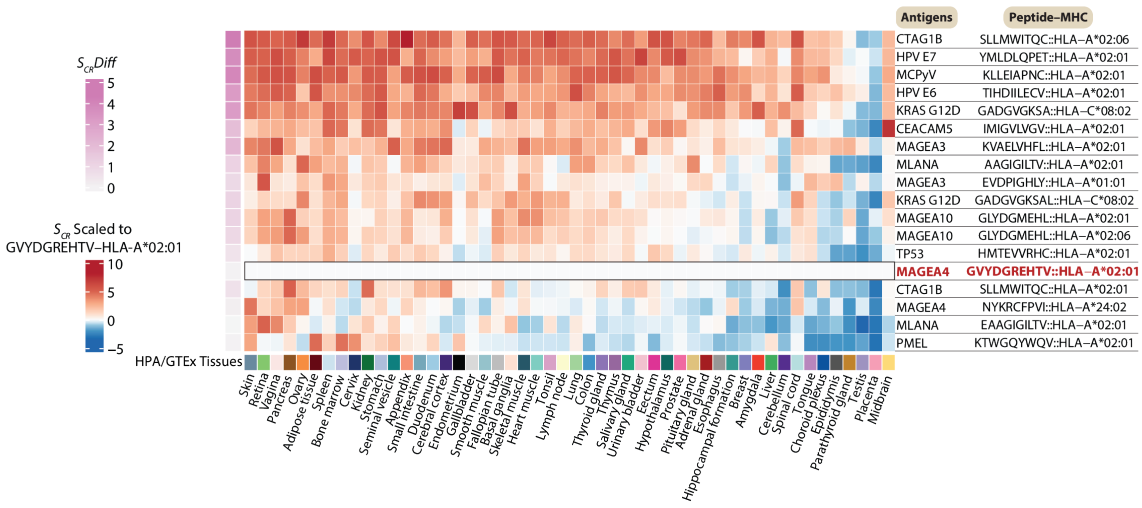

**Figure S8**. **Validation of peptide similarity and normal tissue cross-reactivity algorithms**.

**(A)** The table on the *left* shows the results of the peptide similarity analysis on known pairs of cross-reactive peptides. The density plot on the *right* shows the distribution of the peptide similarity scores ($S_{sim}$) obtained between known cross-reactive peptide pairs (blue curve) and those obtained between each tested peptide and all the peptides of same length binding to the same HLA allele as reported in the normal ligandome (red curve). The vertical line shows the $S_{sim}$ score cutoff of 1.15, used to define highly similar peptides as it provided the highest specificity in identifying known cross-reactive peptides. **(B)** Heatmap showing the HPA/GTEx tissue-specific $S_{CR}$ scores for $pMHCs$ targeted by TCR-T cells that have been tested in clinical trials, and for which safety outcomes are available. For each $pMHC$, the parent antigen is displayed. Afami-cel's target, GVYDGREHTV–HLA-A*02:01, is highlighted for comparison. **(C)** Heatmap showing the HPA/GTEx tissue-specific $S_{CR}$ scores for each known $pMHC$ target scaled to the $S_{CR}$ scores of afami-cel's target, GVYDGREHTV–HLA-A*02:01, and obtained as follows: $\left(\frac{\left(S_{CR}^{j}(target)-S_{CR}^{j}(reference)\right)}{\sigma_j}\right) \times 2 \; if \; S_{CR}^{j}(target) > S_{CR}^{j}(reference)$, where $\sigma_j$ is the standard deviation of the $S_{CR}$ values for all the target $pMHCs$ in the $j^{th}$ tissue. $S_{CR}Diff$: differential $S_{CR}$ profile compared to afami-cel's target, GVYDGREHTV–HLA-A*02:01.

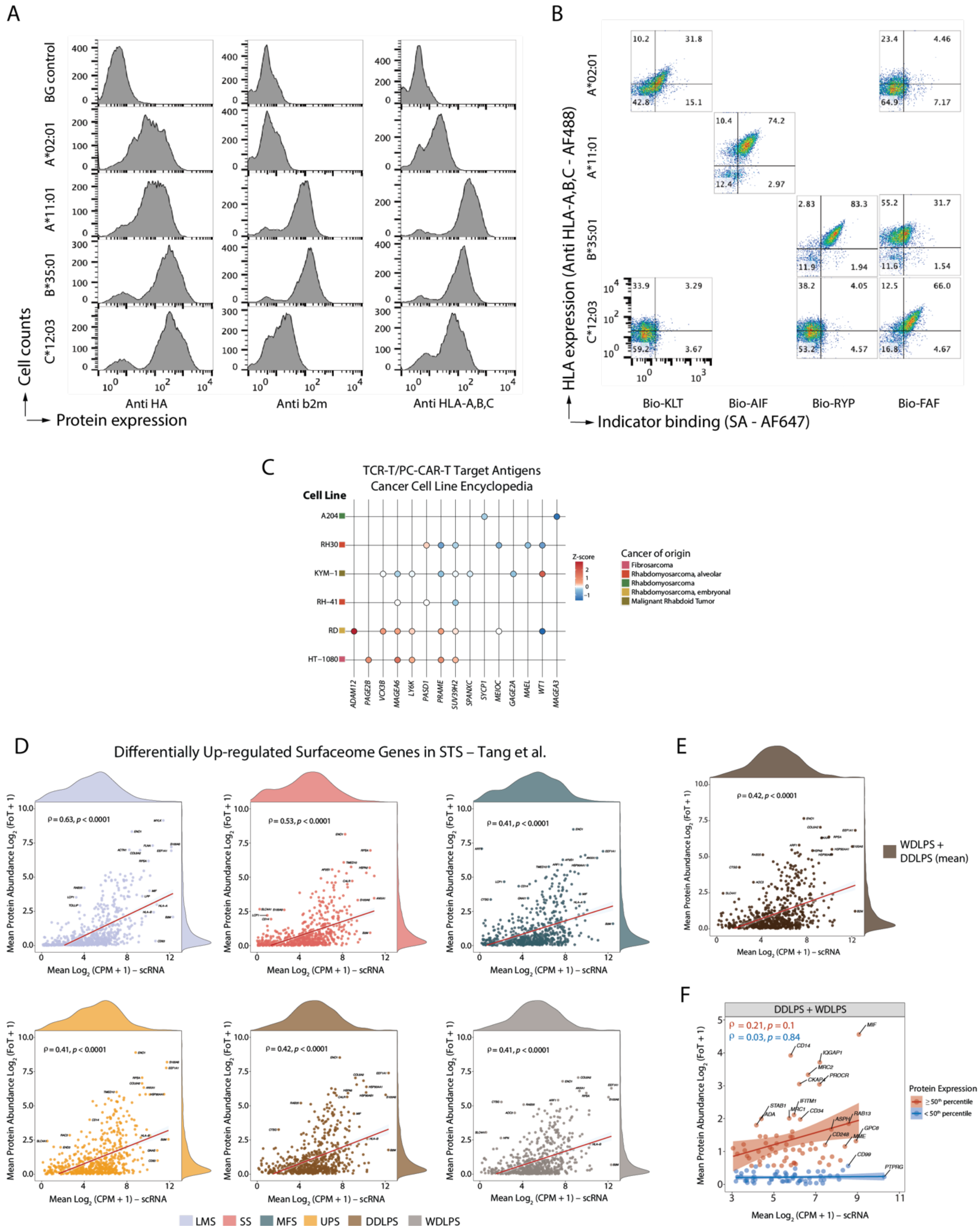
A
BG control
A*02:01
A*11:01
B*35:01
C*12:03
Cell counts
Protein expression
Anti HA
Anti b2m
Anti HLA-A,B,C
B
HLA expression (Anti HLA-A,B,C - AF488)
Indicator binding (SA - AF647)
Bio-KLT
Bio-AIF
Bio-RYP
Bio-FAF
10.2
31.8
42.8
15.1
23.4
4.46
64.9
7.17
10.4
74.2
12.4
2.97
2.83
83.3
11.9
1.94
55.2
31.7
11.6
1.54
33.9
3.29
59.2
3.67
38.2
4.05
53.2
4.57
12.5
66.0
16.8
4.67
C
TCR-T/PC-CAR-T Target Antigens
Cancer Cell Line Encyclopedia
Cell Line
A204
RH30
KYM−1
RH−41
RD
HT−1080
ADAM12
PAGE2B
VCX3B
MAGEA6
LY6K
PASD1
PRAME
SUV39H2
SPANXC
SYCP1
MEIOC
GAGE2A
MAEL
WT1
MAGEA3
Z-score
Cancer of origin
Fibrosarcoma
Rhabdomyosarcoma, alveolar
Rhabdomyosarcoma
Rhabdomyosarcoma, embryonal
Malignant Rhabdoid Tumor
D
Differentially Up-regulated Surfaceome Genes in STS – Tang et al.
ρ = 0.63, p < 0.0001
ρ = 0.53, p < 0.0001
ρ = 0.41, p < 0.0001
ρ = 0.41, p < 0.0001
ρ = 0.42, p < 0.0001
ρ = 0.41, p < 0.0001
Mean Protein Abundance Log2 (FoT + 1)
Mean Log2 (CPM + 1) – scRNA
LMS
SS
MFS
UPS
DDLPS
WDLPS
E
ρ = 0.42, p < 0.0001
WDLPS + DDLPS (mean)
F
DDLPS + WDLPS
ρ = 0.21, p = 0.1
ρ = 0.03, p = 0.84
MIF
CD14
IQGAP1
MRC2
PROCR
CKAP4
STAB1
IFITM1
ADA
MRC1
CD34
ASPH
RAB13
CD248
GPC6
MME
CD99
PTPRG
Protein Expression
≥ 50th percentile
< 50th percentile

**Figure S9**. **Validation of peptide-MHC targets and protein to RNA correlation in the Tang et al. dataset.**

**(A)** Histograms showing the mean fluorescence intensity (MFI) for antibodies binding the HA tag, soluble β2m, and either HLA-A*02:01, A*11:01, B*35:01, or C*12:03, used in the IMMUNE workflow. **(B)** Scatter plots showing MFI for binding indicator to either HLA-A*02:01, A*11:01, B*35:01, or C*12:03, for the control peptides KLTPLCVTL, AIFQCSMTK, RYPVCKFPSL, and FAFCRITSF. Peptides with the highest binding for a given HLA allele were used as positive controls to assess the HLA binding of test peptides, as shown in Figure 5B. **(C)** Dot plot showing protein expression in STS cell lines in the CCLE database. **(D)** Scatter plots showing Spearman correlation between single cell RNA and protein expression in the Tang et al. STS proteomics dataset for LMS, UPS, SS, MFS, DDLPS, WDLPS. The genes plotted for each sarcoma histotype are the up-regulated surfaceome genes for each histotype that are also measured in the Tang et al. dataset. RNA expression is the average $\log_2$(CPM+1) across all tumor cells for a given STS histotype. Protein expression is reported as the average $\log_2$(FoT+1) protein abundance across all samples for a given STS histotype. **(E)** Scatter plots showing Spearman correlation between single cell RNA and protein expression in the Tang et al. STS proteomics dataset for both WDLPS and DDLPS, considering the differentially up-regulated surfaceome genes in liposarcoma (LPS). RNA expression is the average $\log_2$(CPM+1) across all LPS cells in our scRNA-seq data. Protein expression is reported as the average $\log_2$(FoT+1) protein abundance across all DDLPS and WDLPS samples. **(F)**. Scatter plot showing Spearman correlation analysis between single cell RNA expression and protein abundance in the Tang et al. STS proteomics dataset for both WDLPS and DDLPS, considering only the monospecific CAR targets defined in LPS. RNA expression is the average $\log_2$(CPM+1) across all LPS cells in the scRNA-seq dataset. Protein expression is reported as the average $\log_2$(FoT+1) protein abundance across all

DDLPS and WDLPS samples. Genes are separated based on the median protein abundance.

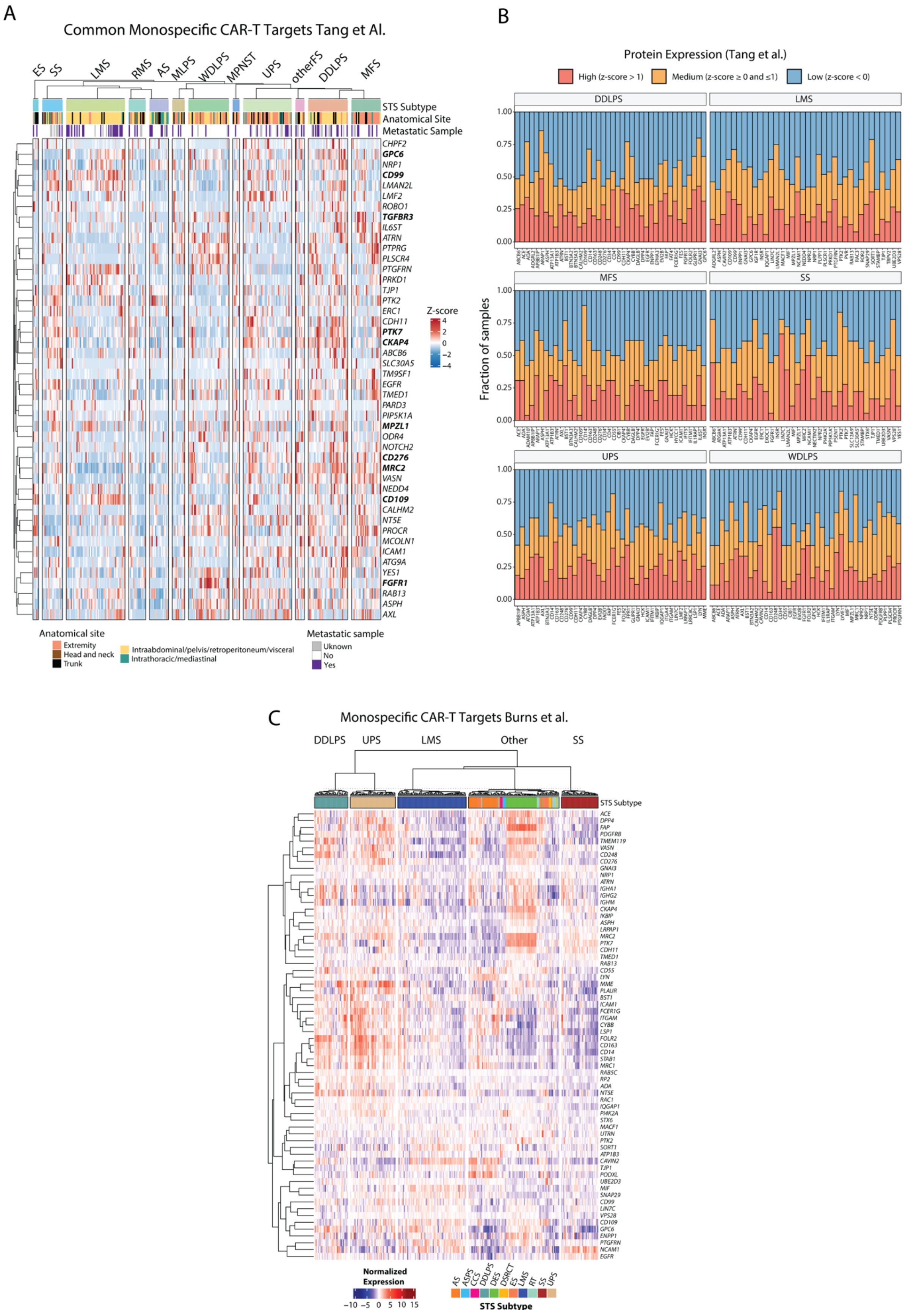
A
Common Monospecific CAR-T Targets Tang et Al.
ES
SS
LMS
RMS
AS
MLPS
WDLPS
MPNST
UPS
otherFS
DDLPS
MFS
STS Subtype
Anatomical Site
Metastatic Sample
CHPF2
GPC6
NRP1
CD99
LMAN2L
LMF2
ROBO1
TGFBR3
IL6ST
ATRN
PTPRG
PLSCR4
PTGFRN
PRKD1
TJP1
PTK2
ERC1
CDH11
PTK7
CKAP4
ABCB6
SLC30A5
TM9SF1
EGFR
TMED1
PARD3
PIP5K1A
MPZL1
ODR4
NOTCH2
CD276
MRC2
VASN
NEDD4
CD109
CALHM2
NT5E
PROCR
MCOLN1
ICAM1
ATG9A
YES1
FGFR1
RAB13
ASPH
AXL
Z-score
4
2
0
−2
−4
Anatomical site
Extremity
Head and neck
Trunk
Intraabdominal/pelvis/retroperitoneum/visceral
Intrathoracic/mediastinal
Metastatic sample
Uknown
No
Yes
B
Protein Expression (Tang et al.)
High (z-score > 1)
Medium (z-score ≥ 0 and ≤1)
Low (z-score < 0)
DDLPS
LMS
MFS
SS
UPS
WDLPS
Fraction of samples
1.00
0.75
0.50
0.25
0.00
C
Monospecific CAR-T Targets Burns et al.
DDLPS
UPS
LMS
Other
SS
STS Subtype
Normalized Expression
−10 −5 0 5 10 15
AS
ASPS
CCS
DDLPS
DES
DSRCT
ES
LMS
RT
SS
UPS
STS Subtype

**Figure S10. Expression of monospecific CAR-T targets in the Tang et al. and Burns et al. STS clinical proteomics datasets**.

**(A)** Protein expression in the Tang et al. dataset for monospecific CAR-T targets common to all histotypes. Expression is z-scored and median-centered on the whole dataset. Notable genes are in bold. **(B)** Stacked bar chart shows the fraction of samples with high (z-score > 1), medium ($0 \leq$ z-score $\leq 1$), and low (z-score $< 0$) expression in the DDLPS, WDLPS, MFS, UPS, LMS, and SS samples of the Tang et al. STS dataset for the monospecific CAR targets. Only the top 40 proteins in each histotype with fraction of samples with medium and high expression ≥ 40% are shown. **(C)** Protein expression in the Burns et al. dataset for all the monospecific STS CAR-T targets identified. Expression is normalized through z-scoring and median-centering. MPNST: malignant peripheral nerve sheath tumor; MLPS: mucinous liposarcoma; others: other fibrosarcoma; ES: Ewing sarcoma; RMS: rhabdomyosarcoma; AS: angiosarcoma; ASPS: alveolar soft part sarcoma; CCS: clear cell sarcoma; DES: desmoid tumor; DSRCT desmoplastic small round cell tumor; RT: rhabdoid tumor.
